# Development of a NanoBRET assay platform to detect intracellular target engagement in living cells for the Glycine Transporter 1 (SLC6A9)

**DOI:** 10.64898/2026.10.05.756331

**Authors:** Malene H. Jensen, Ioannis P. Avgeris, Nur M. Kocaturk, Amalie N. L. Mikkelsen, Bente Frølund, William Farnaby, Petrine Wellendorph, Jonas S. H. Mortensen, Francesco Bavo

## Abstract

Glycine transporter 1 (GlyT1) is a clinically investigated target whose conformational plasticity enables ligands to engage distinct extracellular and intracellular binding sites. However, methods to directly quantify intracellular binding remain limited and largely rely on radioligand-based approaches. Here, we report a NanoBRET-based binding assay to quantify intracellular GlyT1 target engagement in living cells using a fluorescent tracer derived from the GlyT1 inhibitor Cmpd1. With a structure-based approach followed by pharmacological characterization we identified a suitable meta-substituted exit vector for linker and fluorophore attachment while maintaining GlyT1 binding. A series of fluorescent probes was synthesized and evaluated, leading to the development of GlyT1-probe which displayed high-affinity binding and enabled robust NanoBRET measurements. Unlike the functional uptake assay, the NanoBRET assay distinguished intracellular from extracellular GlyT1 inhibitors and resolved structure-activity relationships at the intracellular binding site. This assay provides a quantitative approach to interrogate conformation-selective GlyT1 target engagement in living cells, facilitating the characterization of GlyT1 ligands targeting the intracellular site.

## INTRODUCTION

Solute carriers (SLCs) constitute the largest superfamily of transmembrane proteins and mediate the movement of diverse substrates across cellular membranes.^1^ Members of the solute carriers 6 (SLC6) family are secondary active co-transporters responsible for membrane uptake of neurotransmitters and amino acids.^2^ Within the SLC6 family, the Glycine Transporter 1 (SLC6A9) is a Na^+^/Cl^+^ dependent symporter primarily localized at the plasma membrane of neurons and astrocytes, where it contributes to regulating glycine levels in the brain^3^ Given the role of glycine as an obligate co-agonist at NMDA receptors, GlyT1 has been extensively investigated as a therapeutic target for schizophrenia, with the GlyT1 inhibitors bitopertin and iclepertin advancing to Phase III clinical trials.^4^ More recently, peripheral inhibition of GlyT1 has gained clinical relevance,^5^ with bitopertin being repurposed for the treatment of erythropoietic protoporphyria.^6^ To uptake its substrate, GlyT1 cycles through different conformational states: it binds glycine in an outward-open conformation, transitions to an occluded state, to then release glycine in the cytosol in an inward-open conformation. GlyT1 can also operate in reverse mode, binding glycine first in an inward-open conformation from the cytosolic side and transporting it outside the cell.^7^ The importance of the inward-facing state was highlighted by Shahsavar et al. in 2021, who discovered by X-Ray crystallography that the GlyT1 inhibitor **Cmpd1** (Figure 1) stabilizes the inward-open conformation binding an intracellular pocket exposed to the cytosol.^8^ Accumulating structural and pharmacological evidence now indicates that a corresponding intracellular binding site exists across multiple SLC6 transporters, for example GAT1^9, 10^ and GAT3.^11, 12^ Further structural studies have revealed that GlyT1 can be captured in distinct conformational states depending on the bound ligand. The apo transporter adopts an inward-open conformation, which is likewise stabilized by inhibitors such as **Cmpd1, iclepertin** and **ALX-5407**, whereas **SSR504734** and **PF-03463275** bind preferentially to the outward-open conformation (Figure 1). ^8, 13, 14^ These observations underscore the conformational plasticity of GlyT1 and highlight the intracellular binding pocket as a pharmacologically relevant and structurally tractable site.

**Figure 1.**
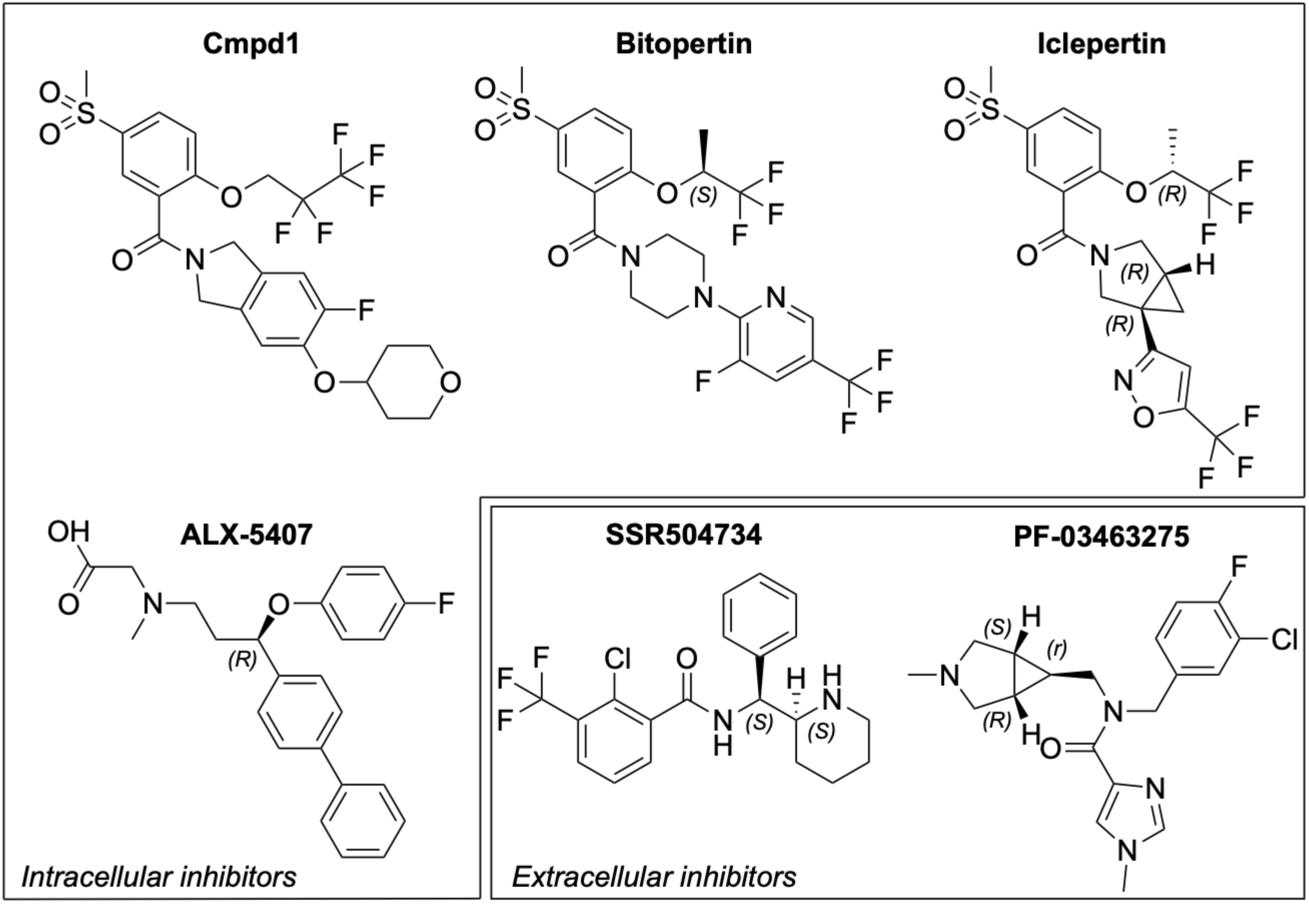
Chemical structures of GlyT1 inhibitors, classified into intracellular and extracellular inhibitors according to available X-Ray and Cryo-EM structures.

The inward-open conformation of GlyT1 and the cytosolic accessibility of its intracellular cavity suggest that GlyT1 inhibitors can be repurposed as chemical handles for the development of conformationally selective probes. Inspired by recent advances in NanoBRET-based target-engagement assays for GPCRs, where high-affinity ligands are modified with linkers and fluorophores to quantify intracellular binding in living cells,^15^ we envisioned that the GlyT1 inhibitor **Cmpd1** could be converted into a NanoBRET tracer capable of reporting on intracellular target engagement in living cells and surpassing the limitations of classic radioligand-based assays.^16^ NanoBRET (Nanoluciferase (NLuc)-based bioluminescence resonance energy transfer) relies on the energy transfer between a target fused to NLuc and the proximal fluorophore of a bound small molecule tracer. Outcompeting the tracer with an inhibitor enables quantification of target-ligand binding affinity in cells. Here, we report on the development of **GlyT1-probe**, a NanoBRET probe that provides a direct, quantitative readout of intracellular inhibitor binding to GlyT1 in a cellular environment, enabling interrogation of conformation-selective pharmacology and accelerating the discovery and characterization of ligands acting at the cytosolic site.

## RESULTS AND DISCUSSION

### Design and synthesis of test probes to identify a suitable exit vector from GlyT1

To identify a suitable exit vector from GlyT1 where a linker and a fluorophore can be attached while maintaining binding, we designed and developed a series of test probes following a structure-based approach. First, we analyzed the X-ray crystallographic data showing GlyT1 bound to Cmpd1 (PDB ID: 6ZBV).^8^ Inspection of the binding site (SiteMap)^17^ revealed a 600 Å^3^ cavity exposed towards the cytosol, characterized by a relatively narrow entrance defined by the residues Tyr116, Asn386, Asp395, Met382 and located near the tetrahydropyran moiety of Cmpd1 (Supplementary figure 1A). To facilitate downstream chemistry efforts, we docked a previously reported simplified analog of Cmpd1,^18^ where synthetic accessibility was increased by removing the fluoride substituent and by replacing the tetrahydropyran ring with a phenyl ring. The predicted binding mode of the unsubstituted compound closely aligned with that of Cmpd1, maintaining key interactions with the binding site (Supplementary figure 1B). As both the *meta*- and *para*-positions of the phenyl ring appeared to provide viable geometries for linker attachment, we designed and synthesized the *meta*-substituted compounds ***m-*1a** to ***m*-1d** and *para-*substituted ***p*-1a** to ***p*-1d** (Figure 2) to experimentally identify an exit vector to further construct the desired NanoBRET probes. The compounds were synthesized adapting a previously reported synthetic scheme (Supplementary figure 2).^18^

**Figure 2.**
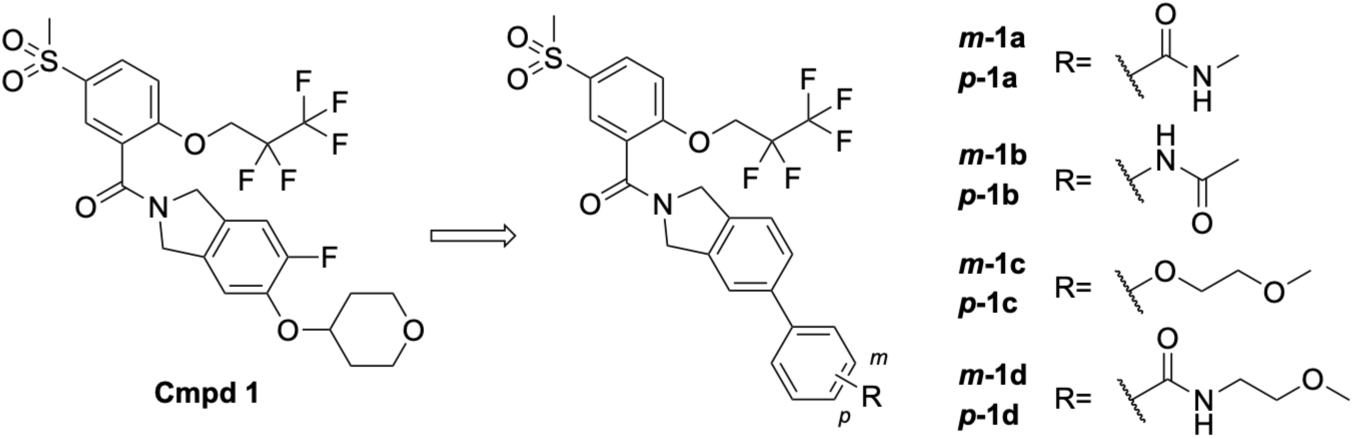
Design of test probes, exploring the *meta-*and the *para-* position as possible exit vectors from GlyT1.

### Pharmacological evaluation of test probes to identify a suitable exit vector from GlyT1

Structural studies demonstrated that the bitopertin analog iclepertin and ALX-5407 bind to the same pocket of Cmpd1 at the inward-open conformation of GlyT1.^13, 14^ In our previous study on GAT3 (SLC6A11), we observed that ligands stabilizing the inward-open conformation exhibited a time-dependent increase in apparent potency, consistent with slow-binding behavior that we attributed to membrane permeation.^11^ Based on this observation, we investigated whether pre-incubation similarly affects the apparent potency of bitopertin at GlyT1. Consistent with this hypothesis, bitopertin exhibited a time-dependent increase in apparent potency upon pre-incubation. In the absence of pre-incubation, bitopertin exhibited an incomplete inhibition, while 30 minutes of pre-incubation resulted in a IC_50_ of 11.2 nM (pIC_50_ ± SEM, *n*; 7.97 ± 0.06, *4*) (Figure 3A). Pre-incubation of 60 and 120 minutes elicited similar apparent potencies of 6.1 nM (pIC_50_ ± SEM, *n*; 8.23 ± 0.06, *4*) and 8.3 nM (pIC_50_ ± SEM, *n*; 8.08 ± 0.04, *3*), respectively, with no significant differences of pIC_50_ values between the two conditions (*p* = 0.12, two-sided Welch’s t-test) (Figure 3A). Hence, the apparent potency was considered to plateau after 60 minutes of preincubation, and therefore a 60-minute preincubation was chosen for subsequent experiments.

**Figure 3.**
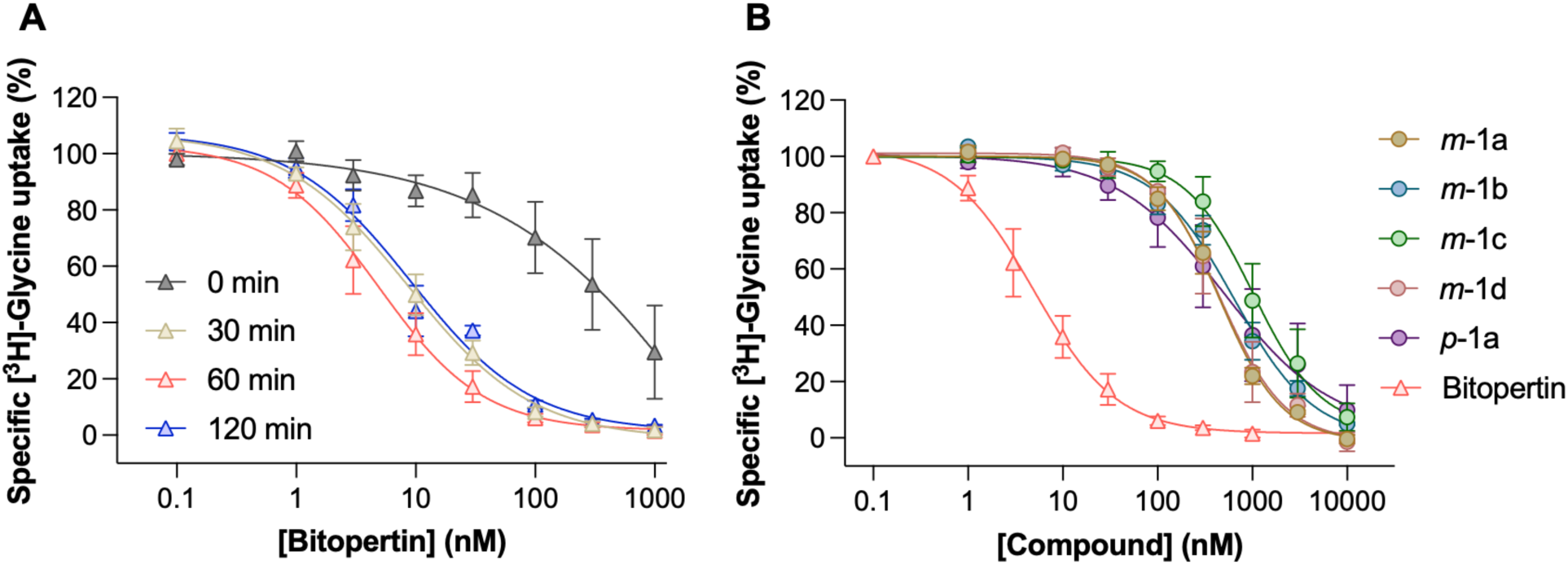
Effect of preincubation time on bitopertin-mediated inhibition of wild-type GlyT1-mediated [^3^H]glycine uptake and inhibitory activity of test-probes. A) Concentration-response curves for bitopertin in HEK293T cells expressing GlyT1 WT without preincubation (grey) and following 30 (beige), 60 (pink) and 120 minutes (blue) of preincubation. B) Concentration-response curves for the indicated test probes upon 60 minutes of preincubation in HEK293T cells expressing GlyT1 WT. The 60 minutes bitopertin concentration-response curve from A is shown as a reference for comparison in B. Data are normalized to maximal [^3^H]-Glycine uptake and presented as mean ± SEM of residual [^3^H]-Glycine uptake from *n* =3-4 biological replicates. Mean IC_50_ of test probes values are summarized in table 1.

**Table 1:**
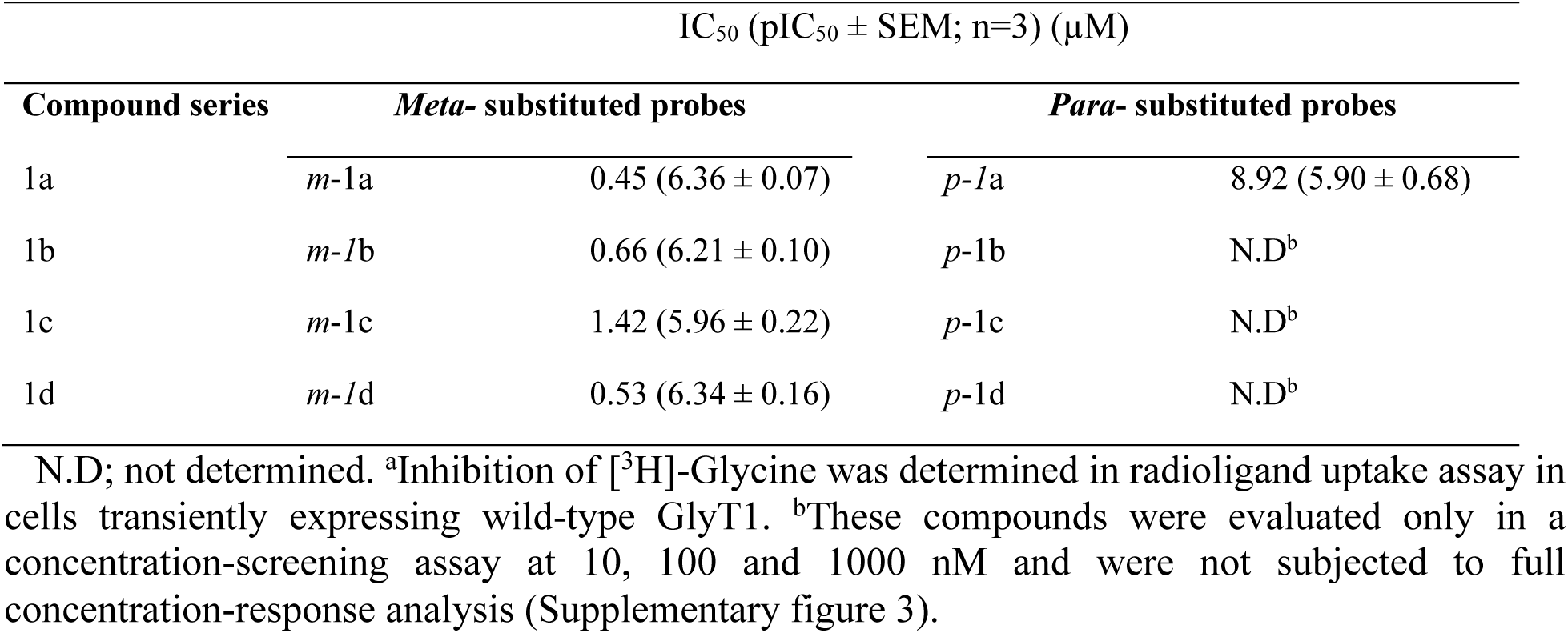
Inhibitory activity of *meta*- and *para*-substituted probes at wild-type GlyT1^a^

| IC<sub>50</sub> (pIC<sub>50</sub> ± SEM; n=3) (μM) |  |  |  |  |
| --- | --- | --- | --- | --- |
| Compound series | *Meta*- substituted probes |  | *Para*- substituted probes |  |
| 1a | *m*-1a | 0.45 (6.36 ± 0.07) | *p*-1a | 8.92 (5.90 ± 0.68) |
| 1b | *m*-1b | 0.66 (6.21 ± 0.10) | *p*-1b | N.D<sup>b</sup> |
| 1c | *m*-1c | 1.42 (5.96 ± 0.22) | *p*-1c | N.D<sup>b</sup> |
| 1d | *m*-1d | 0.53 (6.34 ± 0.16) | *p*-1d | N.D<sup>b</sup> |

N.D; not determined. <sup>a</sup>Inhibition of [<sup>3</sup>H]-Glycine was determined in radioligand uptake assay in cells transiently expressing wild-type GlyT1. <sup>b</sup>These compounds were evaluated only in a concentration-screening assay at 10, 100 and 1000 nM and were not subjected to full concentration-response analysis (Supplementary figure 3).

Having established that a 60-minute preincubation period was sufficient to reach maximal apparent bitopertin potency, we next evaluated the inhibitory activity for test probes ***m*-1a** to **-d** and ***p*-1a** to **-d**. All test probes were screened at three concentrations upon 60 minutes preincubation (Supplementary figure 3). 5 compounds exhibiting >50 % inhibition at 1000 nM were selected for further evaluation and were progressed to full concentration–response analysis, alongside bitopertin as a reference compound (Figure 3B, Table 1).

Overall, all compounds were less potent than bitopertin but displayed inhibitory potencies spanning the nanomolar to low-micromolar range. Only moderate differences in potency were observed across the compound series, with *meta*-substituted compounds ***m*-1a** to **-d** being more potent. Rather than prioritizing the most potent compounds, selection was guided by structure– activity trends related to linker elongation. In particular, *meta*-substituted analogues showed minimal loss of potency upon increasing substituent length (e.g., ***m-1*a** vs ***m*-1d**), whereas *para*-substituted analogues exhibited a more pronounced decrease in activity with elongation. On this basis, the *meta*-position was selected as the preferred exit vector for NanoBRET probe development.

### Design and preparation of NanoBRET probes and constructs

Building on the structure of ***m*-1b** and on the fluorophore BODIPY-590, selected as BRET acceptor due to its excellent spectral resolution when paired with the BRET donor NLuc,^19, 20^ we designed the three NanoBRET probes ***m*-1e** to **-g** exploring linkers of different length and nature (Figure 4). The compounds were synthesized by late-stage assembly via copper-catalyzed azide-alkyne cycloaddition (Supplementary figure 4).

**Figure 4.**
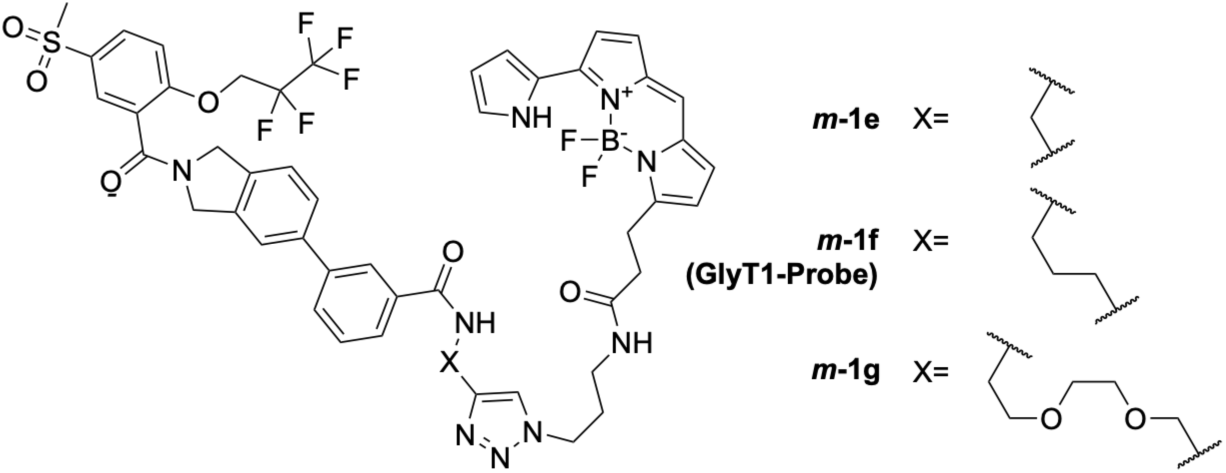
Design of NanoBRET probes.

To enable NanoBRET-based assessment of GlyT1 target engagement, we designed a GlyT1 fusion construct with the luminescent NLuc tag serving as a donor positioned on the cytosolic side of the transporter, proximal to the intracellular ligand binding site. As both the N- and C-termini of GlyT1 are intracellular, either terminus would in principle be suitable for NLuc fusion. However, the C-terminus was selected (Figure 5A and Supplementary Figure 5), based on prior evidence demonstrating that C-terminal tagging with GFP preserves GlyT1 function^13^ and that N-linked glycosylation of GlyT1 is critical for proper trafficking to the plasma membrane.^21^

**Figure 5.**
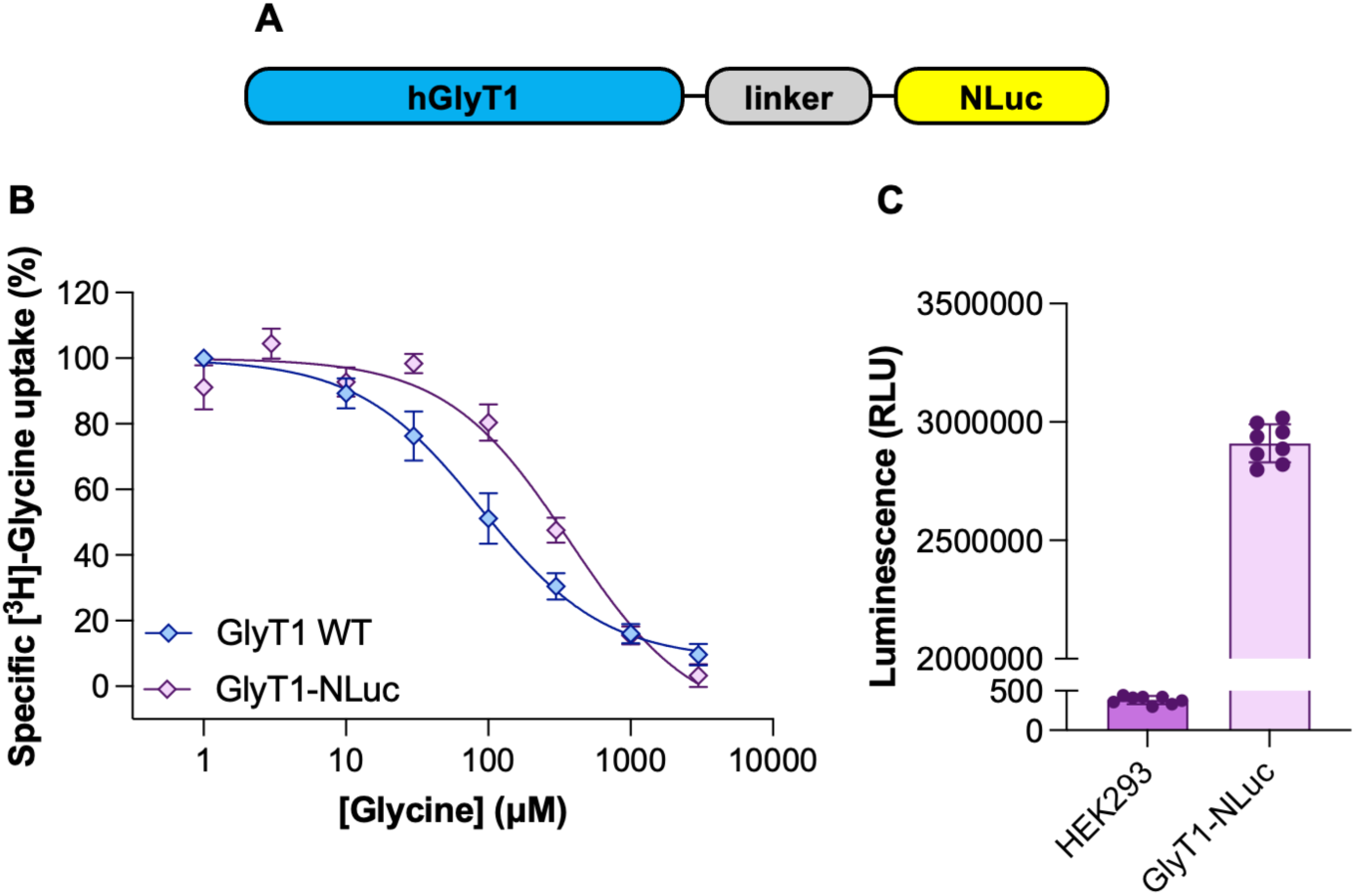
Characterization of a functional GlyT1-NLuc protein. **A)** Cartoon representation of the GlyT1-NLuc fusion construct. **B)** Concentration-response curves for glycine in HEK293T cells transiently expressing GlyT1 WT (blue) or GlyT1-NLuc (purple). Data are normalized to maximal [^3^H]-Glycine uptake and presented as mean ± SEM of residual [^3^H]-Glycine uptake from *n* =3-4 biological replicates and summarized in Table 2. pIC_50_ values for GlyT1-NLuc and GlyT1 WT were compared using an unpaired t-test with Welch’s correction, *p* = 0.0800. **C)** Representative luminescence signals representing NLuc levels in GlyT1-NLuc over-expressed (light purple bar) and untransfected HEK293cells (dark purple bar). Luminescence signal was measured in one independent experiment, and individual data points represent technical replicates. RLU, relative luminescence units.

**Table 2:** Inhibitory activity of NanoBRET probes^a^.

| Compound | $\text{IC}_{50}$ ($\text{pIC}_{50} \pm \text{SEM}$, $n$) ($\mu\text{M}$) | |
| --- | --- | --- |
|  | GlyT1-NLuc | GlyT1-WT |
| Glycine | 300 ($3.52 \pm 0.03$, 4) | 96 ($4.09 \pm 0.17$, 3) |
| ***m-1e*** | 0.040 (7.42 ± 0.08, 3) | N.D<sup>b</sup> |
| ***m-1f* (GlyT1-probe)** | 0.032 (7.52 ± 0.08, 3) | 0.033 μM (7.50 ± 0.09, 3) |
| ***m-1g*** | 0.097 (7.03 ± 0.09, 3) | N.D<sup>b</sup> |

<sup>a</sup>Potency was determined in a [<sup>3</sup>H]-Glycine radioligand uptake assay in cells transiently expressing either GlyT1-NLuc or wild-type GlyT1. Data are summarized from concentration-response curves in Figure 5B and 6A. <sup>b</sup>N.D; not determined.

Consistent with these reports, HEK293T cells transiently expressing the C-terminally NLuc-tagged GlyT1 exhibited robust [^3^H]-glycine uptake, which could be inhibited in a dose-dependent manner by unlabelled glycine with an IC_50_ of 300 µM (pIC_50_ ± SEM, *n*; 3.52 ± 0.03, *4*), not significantly different to that of untagged WT GlyT1 with an IC_50_ of 96 µM (pIC_50_ ± SEM, *n*; 4.09 ± 0.17, *3*) (*p*=0.0800, two-sided Welch’s t-test comparing pIC_50_ values) (Figure 5B, table 2). Hence, the C-terminal NLuc fusion preserves the GlyT1-mediated glycine transport. As a control for the subsequent experiments encompassing the preincubation procedure, glycine was preincubated for 120 minutes prior to [^3^H]-glycine uptake measurement. As expected, preincubation of did not alter the apparent potency of glycine at GlyT1-NLuc compared to the corresponding experiment without preincubation (Supplementary figure 6), supporting the use of preincubation for ligands in subsequent experiments.

NLuc expression levels were verified by comparing luminescence levels in HEK293 cells transiently over-expressing GlyT1-NLuc compared to untransfected cells (Figure 5C). This experiment was performed as a representative validation of NLuc activity, with the NLuc luminescence/donor signal being confirmed for each independent NanoBRET experiment.

### Pharmacological characterization of NanoBRET probes and NanoBRET assay development

The inhibitory activity of NanoBRET probes ***m*-1e** to **–g** at GlyT1-NLuc was evaluated using a [³H]glycine uptake inhibition assay in HEK293T cells transiently expressing the GlyT1–NLuc construct, following a 2 h pre-incubation protocol. All three probes exhibited concentration-dependent inhibition of glycine uptake (Figure 6A and Table 2), with IC₅₀ values of 40 nM (***m*-1e**), 32 nM (***m*-1f**), and 97 nM (***m*-1g**). The higher potency observed for probes bearing more lipophilic linkers is consistent with the predominantly lipophilic environment surrounding the selected exit vector. Potency was confirmed at untagged WT GlyT1 for the most potent compound ***m*-1f** (33 nM, Supplementary figure 7) which was renamed as **GlyT1-probe** (Figure 6B) and chosen for further NanoBRET assay optimization (Figure 6C). To estimate the apparent binding affinity of **GlyT1-probe**, we performed NanoBRET saturation binding experiments in live HEK293T cells transiently expressing GlyT1–NLuc, following a 2 h tracer pre-incubation (Figure 6D). Non-specific binding was defined in parallel wells by co-incubation with an excess (25 µM) of the reference GlyT1 inhibitor bitopertin as a competitor. **GlyT1-probe** produced a concentration-dependent increase in NanoBRET signal with a K_D_ of 11 nM (pK_i_ ± SEM, *n*; 7.96 ± 0.06, *3*) that was effectively abolished in the presence of bitopertin, indicating specific binding at the intracellular GlyT1 site.

**Figure 6.**
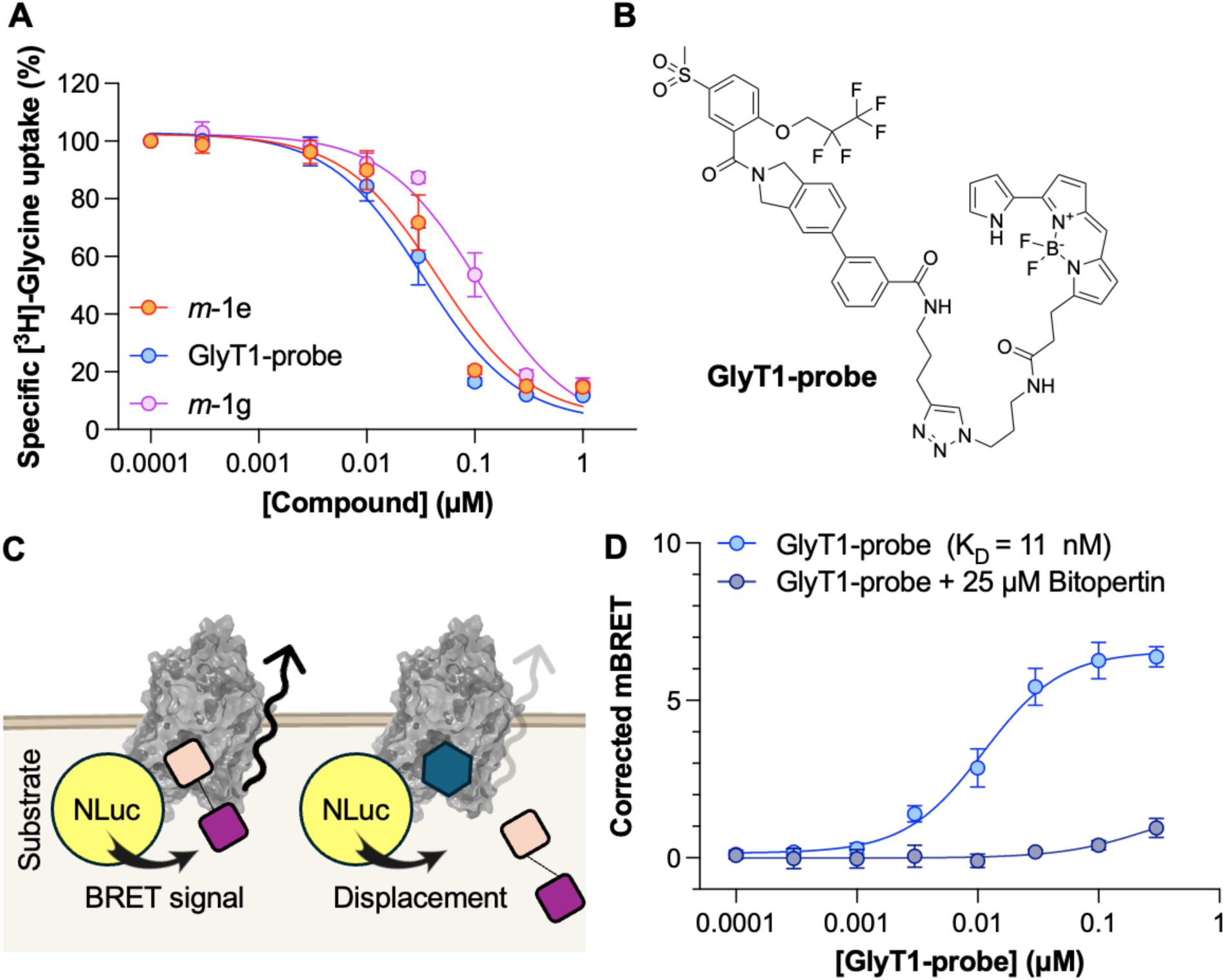
Selection of GlyT1-probe for NanoBRET assay development. A) Concentration-response curves for NanoBRET probes in HEK293T cells transiently expressing GlyT1-NLuc following 120 minutes preincubation obtained in radioligand uptake assay. Data are normalized to maximal [^3^H]-Glycine uptake and presented as mean ±SEM of residual [^3^H]-Glycine uptake from *n* = 3 biological replicates. Mean IC_50_ values are summarized in Table 2. B) Chemical structure of the probe selected for NanoBRET assay development. C) Schematic representation of the GlyT1 NanoBRET target engagement assay. The NLuc (donor, shown as a yellow sphere) emits light at 460 nm upon substrate consumption; the energy is transferred to the bound, fluorescent NanoBRET probe (acceptor, shown as a pink square connected to a purple square, representing the target binding motif and the fluorescent moiety respectively); the NanoBRET probe re-emits light at a longer wavelength; NanoBRET signal is calculated as the acceptor/donor ratio. Outcompetition with a ligand binding to the same site (illustrated with a blue hexagon) reduces the energy transfer. D) GlyT1-probe titration in HEK293T cells transiently expressing GlyT1-NLuc following 120 minutes of preincubation, and outcompetition with an excess of bitopertin. NanoBRET signal (mBU units) is corrected by background subtraction and presented as mean ±EM from *n* = 3 biological replicates (*n* = 2 for the two lowest concentrations).

### Characterizing GlyT1 affinity of GlyT1 inhibitors in live cells

Competitive displacement experiments were conducted using a fixed concentration of **GlyT1-probe** selected to provide an adequate assay windows for competition assays (EC_85_, 0.1 µM)^20^ and a panel of GlyT1 inhibitors with reported intracellular (bitopertin, ALX-5407) or extracellular (SSR504734, PF-03463275) binding modes (Figure 7A). As expected for ligands occupying the intracellular pocket, both bitopertin and ALX-5407 fully displaced **GlyT1-probe**, yielding calculated K_i_ values of 22 nM (pK_i_ ± SEM, *n*; 7.65 ± 0.09, *3*) and 5 nM (pK_i_ ± SEM, *n*; 8.27 ± 0.01, *3*), respectively. In contrast, SSR504734 and PF-03463275 produced only partial displacement and only at high concentrations (56% and 57% displacement at 25 µM respectively). To substantiate that our NanoBRET assay reports on intracellular binding affinity rather than a binding site-independent functional blockade of the transporter, we measured functional potencies of the four reference compounds using the radioligand uptake assay according to standard protocols (Figure 7B). Consistent with this hypothesis, potent inhibition of GlyT1-mediated uptake was measured for the extracellular inhibitors SSR504734 (IC_50_ = 180 nM, pIC_50_ ± SEM, *n*; 6.76 ± 0.07, *3*) and PF-03463275 (and IC_50_ = 41 nM, pIC_50_ ± SEM, *n*; 7.39 ± 0.08, *3*) despite showing little to no activity in the NanoBRET assay. Notably, the extracellular inhibitors SSR504734 and PF-03463275 exhibited greater functional potency than the intracellular inhibitors bitopertin (c.a. 70% inhibition at 1 µM, *n* = 4) and similar or greater than ALX-5407 (IC_50_ = 0.18 µM, pIC_50_ ± SEM, *n*; 6.68 ± 0.07, *3*), presumably because of their immediate access to the extracellular binding site. These findings highlight that functional assays are ideal for measuring overall inhibition of transporter activity, whereas the NanoBRET assay is able to discriminate compounds on the basis of intracellular target engagement. This is a key enabling step towards identifying intracellular binders, important for applications such as induced-proximity strategies and other mechanism-specific chemical tools and inhibitors.

**Figure 7.**
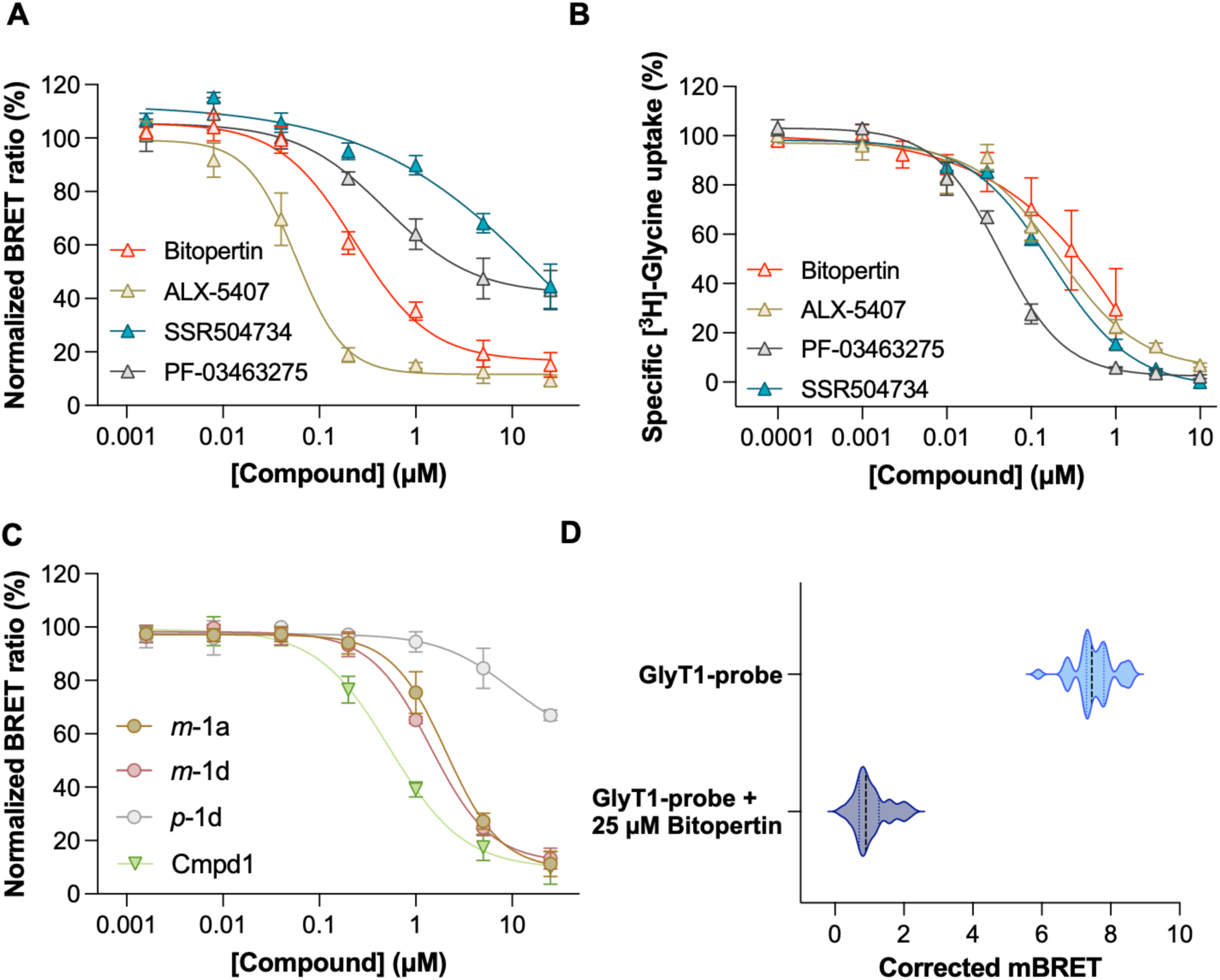
NanoBRET displacement for standard compounds to report on intracellular binding. **A)** Competition binding curves of standard intracellular and extracellular GlyT1 inhibitor in the NanoBRET assay, using a [GlyT1-probe] of 0.1 µM. NanoBRET signal (mBRET) is corrected by background subtraction and expressed as percentage relative to vehicle control. Data are presented as mean ± SEM from *n =* 3 biological replicates. **B)** Concentration-response curves for established GlyT1 inhibitors in HEK293T cells expressing WT GlyT1 in the absence of preincubation. Data are normalized to maximal [^3^H]-Glycine uptake and presented as mean ± SEM of residual [^3^H]-Glycine uptake from *n*=3-4 biological replicates. **C)** Competition binding curves of Cmpd1 and test probes in the NanoBRET assay, performed as above. Data are presented as mean ± SEM from *n =* 3 biological replicates. **D)** Z’-factor determination for the NanoBRET-based setup using GlyT1-probe (0.1 µM) and HEK293T expressing GlyT1-Nluc (32-fold determination).

To assess the ability of the NanoBRET assay to resolve structure-activity relationships resulting from minor structural variations, we measured the binding affinities of the precursor compounds ***m*-1a, *m*-1d, *p*-1d** and **Cmpd1** using NanoBRET (Figure 7C). Intracellular GlyT1 engagement was comparable for ***m*-1a** and ***m*-1d**, with calculated K_i_ values of 173 nM (pK_i_ ± SEM, *n*; 6.76 ± 0.06, *3*) and 130 nM (pK_i_± SEM, *n*; 6.88 ± 0.06, *3*) respectively, indicating that elongation at the meta position is well tolerated, as anticipated from our design strategy. In contrast, ***p*-1d** only partially displaced the tracer, even at high concentrations, demonstrating that extension at the para position is not tolerated. These findings validate our structure-based design approach. As expected, the optimized **Cmpd1** exhibited the highest affinity, with a K_i_ value of 54 nM (pK_i_ ± SEM, *n*; 7.27 ± 0.04, *3*).

To evaluate the robustness of the NanoBRET live-cell GlyT1 target-engagement assay and its suitability for screening applications, we determined the Z′ factor (Figure 7D) using untreated controls (tracer only) and bitopertin-treated controls.^22^ Despite a relatively modest assay window, the assay yielded a Z′ factor of 0.60, indicative of good assay quality and suitability for high-throughput screening.

### Effect of ionic conditions on intracellular GlyT1 engagement

The intracellular binding site of GlyT1 is accessible only when the transporter adopts an inward-facing conformation. During the transport cycle, binding of Na^+^ and Cl^−^ stabilizes outward-facing states that are relevant for substrate uptake.^23^ We therefore hypothesized that reducing the concentrations of Na^+^ and/or Cl^−^, or replacing these ions with non-transportable substitutes, could shift the conformational equilibrium toward inward-facing states, thereby increasing accessibility of the intracellular binding pocket and facilitating engagement of intracellular binders. We therefore performed direct NanoBRET saturation experiments with **GlyT1-probe** under a range of buffer conditions. Specifically, Na^+^ was replaced with gluconate, Cl^−^ was replaced with choline, the concentration of both ions was reduced to 25 mM, and both Na^+^ and Cl^−^ were substituted simultaneously using gluconate and N-methyl-D-glucamine (NMDG), respectively. In addition, a nominally salt-free condition was prepared in which pH was adjusted using gluconic acid and NMDG rather than HCl or NaOH to eliminate residual sodium ions. With a nominal NaCl concentration of 137 mM, **GlyT1-probe** showed an apparent direct binding affinity (K_D_) of 11 nM. Replacing a single ion, either Na^+^ with choline or Cl^-^with gluconate (nominally 137 mM of choline chloride or sodium gluconate), resulted in an apparent affinity of 7 and 6 nM, confirming that none of the ions is strictly required for compound binding at the inward open facing conformation (Figure 8A). We then reduced the total concentrations of NaCl to 25 mM, and observed an even further increase of apparent affinity for **GlyT1-probe** (K_D_ = 3 nM with 25 mM NaCl), indicating that high salt concentrations are not required for intracellular binding of the tracer. This effect could arise either from reduced Na^+^/Cl^−^-dependent stabilization of outward-facing states or from changes in ionic strength. To distinguish between these possibilities, NaCl was replaced with N-methyl-D-glucamine (NMDG) and gluconic acid (GA), while retaining or not Na^+^ and Cl^-^from other sources in the medium (Figure 8B). Under both conditions, ionic strength was maintained while the concentrations of Na^+^ and Cl^−^ were substantially reduced or completely removed. In both cases, **GlyT1-probe** displayed a K_D_ of approximately 3 nM.

**Figure 8.**
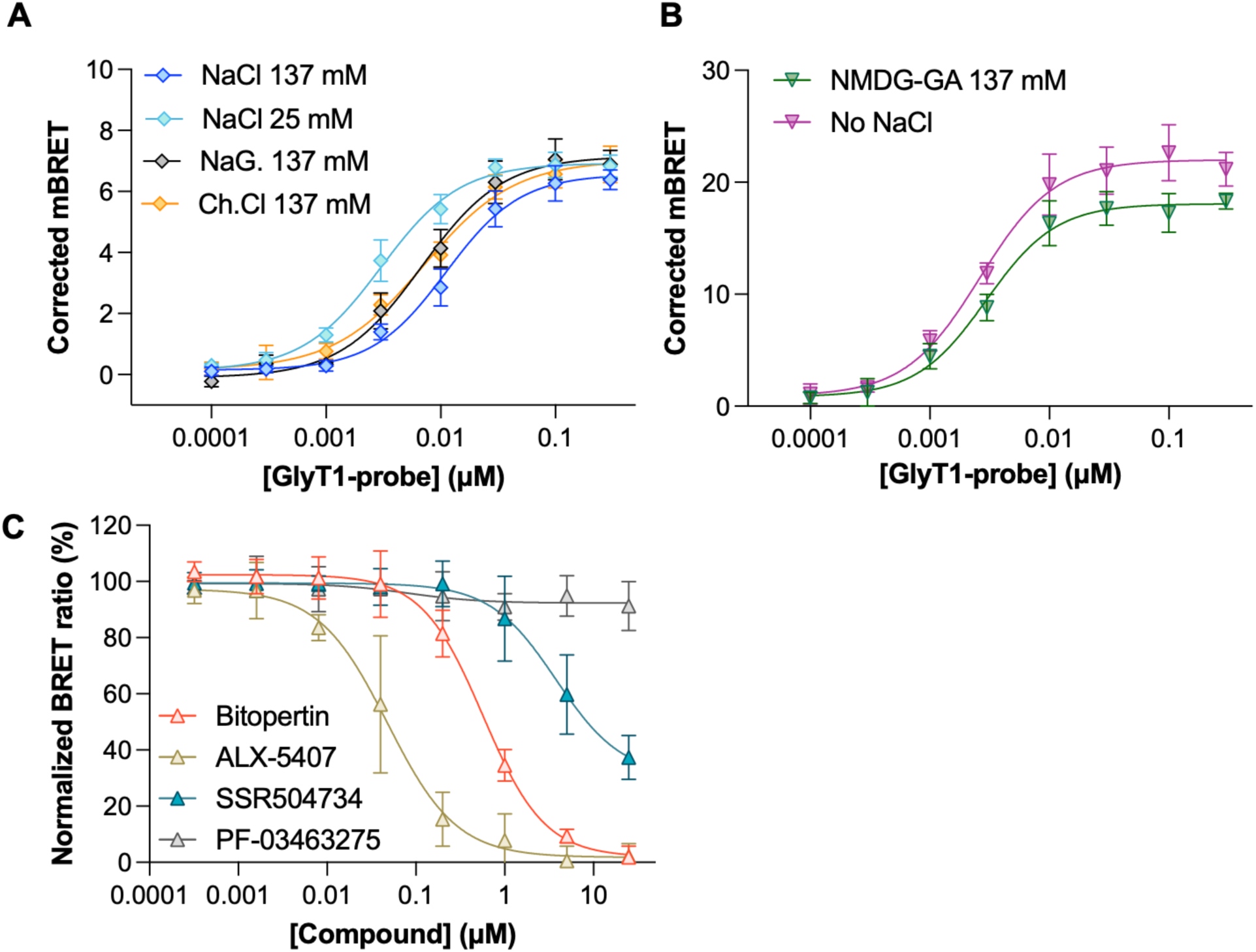
Intracellular probes and inhibitors bind GlyT1 independently from Na^+^ and Cl^-^. **A**, B) GlyT1-probe titration in different buffer conditions: normal NaCl (K_D_ = 11 nM, pK_D_ ± SEM; 7.96 ± 0.06), low sodium concentration (K_D_ = 3 nM, pK_D_ ± SEM; 8.54 ± 0.06), sodium gluconate (K_D_ = 6 nM, pK_D_ ± SEM; 8.22 ± 0.07), choline chloride (K_D_ = 7 nM, pK_D_ ± SEM; 8.17 ± 0.03), NMDGA-gluconic acid (K_D_ = 3 nM, pK_D_ ± SEM; 8.55 ± 0.04), and without any source of NaCl (K_D_ = 2 nM, pK_D_ ± SEM; 8.62 ± 0.04). NanoBRET signal (mBU units) is corrected by background subtraction and presented as mean ± SEM from *n* = 3 biological replicates (*n* = 2 for the two lowest concentrations). C) Competition binding curves of standard intracellular and extracellular GlyT1 inhibitors in the NanoBRET assay using a NaCl-free buffer and a [GlyT1-probe] of 0.01 µM. NanoBRET signal (mBU) is corrected by background subtraction and expressed as percentage relative to vehicle control. Data are presented as mean ± SEM from *n =* 3-4 biological replicates.

Although we cannot conclude on a possible link between increased apparent potency of **GlyT1-probe** to a conformational shift of GlyT1 toward inward-facing states induced by the absence of NaCl, our findings demonstrate that the intracellular **GlyT1-probe** does not require Na^+^ nor Cl^-^ to bind GlyT1. We then investigated whether **GlyT1-probe** could be outcompeted by intracellular and extracellular inhibitors under nominally NaCl-free conditions. Because **GlyT1-probe** exhibited higher affinity under these conditions, the tracer concentration was reduced to 0.01 μM to maintain an appropriate assay window. The intracellular inhibitors bitopertin and ALX-5407 displayed K_i_ values of 110 nM (pK_i_ ± SEM, *n*; 6.96 ± 0.03, *4*) and 9 nM (pK_i_ ± SEM, *n*; 8.03 ± 0.14, *3*) respectively, whereas the K_i_ values could not be calculated for SSR504734 and PF-03463275 due incomplete or no displacement (*n* = 3). Compared to the most potent binder ALX-5407, both extracellular inhibitors SSR504734 and PF-03463275 display right-shifted curves in the absence of NaCl than in normal NaCl conditions.

## CONCLUSIONS

In summary, we developed and validated the first NanoBRET-based target engagement assay for GlyT1 in living cells. The assay reports on intracellular GlyT1 binding affinity and discriminates between intracellular and extracellular ligands, providing a new layer of information that complements traditional radioligand uptake assays. The successful conversion of an intracellular GlyT1 inhibitor into the potent and high affinity NanoBRET tracer **GlyT1-probe** demonstrates that this binding mode tolerates extension toward the cytosol, revealing a viable exit vector that could be exploited for the development of additional chemical probes and induced-proximity modalities.^24–26^ The functional GlyT1-NLuc construct developed in this study may also enable additional bioluminescence-based applications beyond target-engagement measurements.^16^ Additionally, the ability of **GlyT1-probe** and intracellular GlyT1 inhibitors to engage the transporter in the absence of Na^+^ and Cl^−^ demonstrates that intracellular ligand binding is not dependent on these ions and provides a rationale for future studies of GlyT1 conformational dynamics and state-dependent pharmacology. This work lays an important foundation for ongoing induced-proximity approaches targeting GlyT1, and for the development, characterization and optimization of additional probes targeting the intracellular binding site.

## MATERIALS AND METHODS

### Computational methods

The X-ray co-crystal structure of GlyT1 bound to Cmpd1 (PDB ID: 6ZBV) was imported as a .pdb file in Maestro (Schrodinger) and preprocessed using the Protein Preparation Workflow^27^ with default settings. SiteMap was centered on the bound Cmpd1, and a single binding site region was evaluated with default settings. The docking grid was centered on the bound ligand, and docking was performed using Glide Standard Precision.^28^ Docked compounds were previously prepared using LigPrep with default settings.

### Chemistry: materials

Reference compounds (Cmpd1, bitopertin, ALX-5405, SSR504734, PF-03463275), reagents and solvents were purchased from commercial suppliers (Ambeed, MedChemExpress, BLDpharm, Sigma-Aldrich) and used without further purification. The solvents used were of standard HPLC-grade quality. Anhydrous DMF, THF and DCM were obtained from a Glass Contour Solvent System (SG Water USA). Other anhydrous solvents were drying over fresh molecular sieves (3 Å, Sigma-Aldrich) for at least 1 days prior to use. All liquid reagents were had reached room temperature before addition. The heating of reaction mixtures was performed by oil bath and the temperature stated in the experimental section was the temperature of the oil bath.

### Chemistry: general information

Air-sensitive reactions were carried out under nitrogen or argon atmosphere using syringe-septum cap technique, and moisture-sensitive reactions were performed using flame-dried glassware. Thin Layer Chromatography (TLC) was carried out using Merck silica gel 60 F254 aluminium sheets and the compounds were visualized by UV (254 nm). The eluent system is specified in the experimental procedure. Automated Flash Column (FC) Chromatography was carried out on a Combi Flash NextGen 300+. For normal phase (NP) and reverse phase chromatography, RediSep Normal Phase Silver (40-60 micron)/Gold (20-40 micron) Silica columns and RediSep Silver (40-60 micron)/Gold (20-40 micron) C18 columns were employed, respectively. Dry loading was used for sample loading. Samples are dissolved, mixed with celite and dried *in vacuo*. The solid mixture obtained is loaded to a Teledyne ISCO RediSep Rf Cartridge and connected to the flash chromatography system. The eluents A and B were utilized with a gradient elution (isocratic elution upon collection of peaks). For NP chromatography, the specific eluent system is specified in the experimental procedures. Unless otherwise stated, the eluent system for RP chromatography consisted of eluent A (95:5:0.1 Milli-Q H_2_O/ACN/TFA) and B (95:5:0.1 ACN/Milli-Q H_2_O/TFA). Analytical High-performance liquid chromatography (HPLC) was performed on an UltiMate HPLC system (Thermo Scientific) consisting of an LPG-3400A pump (1 mL/min), a WPS-3000SL autosampler, and a DAD-3000D diode array detector using a Gemini-NX C18 column (4.6 × 250 mm, 3 μm, 110 Å). Samples were diluted in 1.0mL of H_2_O/MeCN 1:1 solution and sample volumes 0.5-20μL were injected via autosampler. Gradient elution from 0 to 100% B (MeCN/H_2_O/TFA, 90:10:0.1) in solvent A (H_2_O/TFA, 100:0.1) over 20 min was employed. Data were acquired and processed using Chromeleon Software v. 6.80.

Preparative High-performance liquid chromatography (HPLC) was performed on a Dionex Ultimate 3000 HLPC system comprising a LPG-3200BX pump (20 mL/min), a Rheodyne 9725i injector, a 10 mL loop, an MWD-3000SD detector (200, 210, 254, and 281 nm), and an AFC-3000SD automated fraction collector using a Gemini-NX C18 column (21.2 × 250 mm, 5 μm, 110 Å). Data were acquired and processed using Chromeleon Software v. 6.80. The eluents A (H_2_O/TFA, 100:0.1) and B (MeCN/ H_2_O/TFA, 90:10:0.1) were employed. The conditions (eluent ratio and duration) are stated separately for each case. Nuclear Magnetic Resonance (NMR) Spectroscopy (^1^H, ^13^C, ^19^F, COSY, HSQC, HMBC and ROESY) was acquired using a 400 MHz Bruker Avance III which had a 5 mm broad band probe at 400.09 MHz or a Bruker Avance 600 MHz spectrometer equipped with a 5-mm cryogenically cooled ^13^C/^1^H DCH probe. Proton spectra were acquired at 400.09 MHz or 599.78 MHz, carbon spectra at 100.60 MHz or 150.82 MHz and fluorine spectra at 376.45 MHz. The samples were dissolved in Chloroform-d, Methanol-d_4_, Acetonitrile-d_3_ DMSO-d_6_ and analyzed at 300K. The solvent peaks were used as an intern standard. Data collecting and processing were done using MestReNova software, version 14.3.3. NMR data is reported as chemical shift: d value (ppm), multiplicity: s = singlet, d = doublet, t = triplet, q = quartet, dd = double doublet, ddd = doublet of doublet of doublets, dt = doublet of triplets, td = triplet of doublets, q = quartet, p = pentet, m = multiplet, br = broad signal, coupling constants: *J*-value (Hz) and integral. Liquid Chromatography–Mass Spectrometry (LC–MS) was performed on an Agilent 6130 Mass Spectrometer instrument using electron spray ionization (ESI) coupled to an Agilent 1200 HPLC system (ESI-LC-MS) with a C18 reverse phase column (Zorbax Eclipse XBD-C18, 4.6 mm × 50 mm), autosampler and diode array detector, using a linear gradient of the binary solvent system of buffer A (95:5:0.1 Milli-Q H2O/ACN/formic acid) to buffer B (100:0.1 ACN/formic acid) with a flow rate of 1 mL⁄min. If no LC–MS data are reported for an individual procedure, this indicates that the compound was not UV active or did not ionize. High-resolution mass spectrometry (HRMS) was performed on a QExactive Orbitrap mass spectrometer (Thermo Fisher, Bremen, Germany) equipped with a SMALDI5 ion source (TransMIT GmbH, Giessen, Germany). As a as MALDI matrix, the analytes were mixed with a 30 mg/mL solution of 2,5-dihydroxybenzoic acid in MeOH, and 2 µL of the mixture was deposited on a 30-well glass plate (Electron Microscopy Sciences, Hatfield, PA, USA). The samples were analyzed in the positive ion mode using a peak from the 2,5 dihydroxybenzoic acid matrix for internal mass calibration whereby a mass accuracy of 3 ppm or better was achieved. All final compounds were >95% pure as determined by HPLC. Synthetic routes, experimental procedures and characterization data for all the compounds, and HPLC chromatograms of the most relevant compounds are provided in the Supplementary Information.

### Pharmacology: materials

Dulbecco’s Phosphate Buffered Saline (DPBS), Dulbecco’s modified Eagle medium (DMEM) with GlutaMAX-I, Hanks Balanced Salt Solution (HBSS), Opti-MEM, fetal bovine serum (FBS), penicillin-streptomycin and tryspin-EDTA were purchased from Life Technologies (Paisley, UK). Poly-D-lysine (PDL), Glycine, CaCl_2_, MgCl_2_, 4-(2-hydroxyethyl)piperazine-1ethanesulfonic acid) (HEPES), Dimethyl sulfoxide (DMSO), N-methyl-D-glucamine (NMDG) and choline chloride were purchased from Sigma-Aldrich (St. Louis, MO, USA). D-Gluconic acid 50% in water was purchased from BLDpharm. Polyfect transfection reagent was purchased from Qiagen (West Sussex, UK). [^3^H]glycine (specific activity 42.6 Ci/mmol) and MicroScint^TM^20 were purchased from PerkinElmer. Standard assay buffer was prepared by supplementing HBSS (nominal NaCl concentration of 137 mM) with 20 mM HEPES, 1 mM CaCl_2_ and 1 mM MgCl_2_, and by adjusting the pH to 7.4 using NaOH 2M, for a final calculated concentration of Na^+^ and Cl^-^is 143 mM and 147 mM, respectively. All other assay buffers were prepared in milliQ water from 1M or 0.1M stock solutions of the appropriate salts, and sterilized by filtration through 0.22 µm precision syringe filter (VWR). Calculated concentrations of Na^+^ and Cl^-^ were: 25 mM and 25 mM (for NaCl 25 mM), 143 mM and 9 mM (for NaGluconate buffer), 5 mM and 147 mM (for choline chloride buffer), 5 mM and 9 mM (for NMDG-gluconate buffer) and 0 mM and 0 mM (for no NaCl buffer).

### DNA construct

The human GlyT1 complementary DNA sequence was codon optimized and synthetized by GenScript for expression in mammalian cells. For NLuc experiments, the pBiT3.1-N-CMV vector was modified to carry the sequence of GlyT1 followed by a linker (EFAGSSGGSSG) and the coding sequence for NLuc. For uptake experiments in HEK293T cells, wild-type human GlyT1 construct cloned into a pcDNA3.1 vector was used. All constructs were verified by full plasmid DNA sequencing. The cell lines were regularly tested for mycoplasma contamination and tested negative for contamination. No misidentified cell lines were used. HEK-293T cell line was purchased from ATCC (293T-ATCC; #CRL-3216; authenticated to be mycoplasma-free).

### Cell culture

HEK293T cells were maintained and transfected as previously described.^11^ In brief, HEK293T cells were maintained in DMEM containing GlutaMAX-I, supplemented with 10% FBS and 1% penicillin-streptomycin and incubated at 37 °C with 5% CO2 and routinely subcultured at 80% confluency.

### Transient transfection for radioligand uptake assays

Cells were subcultured one day prior to transfection. At approximately 60% confluency cells were transfected in 10 cm dishes with 4 µg of plasmid, using PolyFect as the transfection reagent according to the manufacturer’s protocol (Qiagen), except that half the volume of transfection reagent was used. Approximately 20 h post-transfection cells were plated in PDL-coated white polystyrene plates (CulturPlate, Perkin Elmer) at a density of 50.000 cells/well. Cells were left overnight and all assays were performed approximately 48 h post-transfection.

### Transient transfection for NanoBRET assays

One day before transfection, HEK293T were plated in a 6-well plate at a density of 5x10^5^ cells/well. 2 mL of DMEM were added, and the cells were incubated for 24 h. On the day of the transfection, a transfection mix of 1 µg total DNA/well in 150 µL of Opti-MEM was prepared, consisting of 900 ng of carrier DNA (1 µg/µL, Promega, E4881) and 100 ng of GlyT1-NLuc DNA (0.1 µg/µL). FuGENE HD transfection reagent was added to the mixture in a 3:1 FuGENE to total DNA ratio, the tube was mixed by inversion and incubated at room temperature for 20 minutes. Afterwards, 150 µL of this mix were added dropwise onto the cells in each well. After 24 hours of incubation, the media was replaced, and after further 24 hours the NanoBRET experiment was performed.

### Radioligand uptake assay

The [^3^H]glycine competition uptake assay was performed as previously described.^11, 29^ In brief, cells were washed with 100 µL/well assay buffer (HBSS, supplemented with 20 mM HEPES, 1 mM CaCl_2_ and 1 mM MgCl_2_, pH 7.4). Subsequently, cells were incubated with 75 µL/well assay buffer containing 30 nM [^3^H]glycine in the presence or absence of test compounds for 3 minutes at 37 °C. Uptake experiments in the presence of preincubation were performed similarly, except that cells were additionally incubated with 75 µL/well compounds in assay buffer for the indicated time at 37 °C before proceeding to the uptake step. Uptake was terminated by three washes with 100 µL/well ice-cold assay buffer. Finally, 150 µL/well scintillation liquid was added and the plate was shaken for 1 hour before scintillation counting was performed in a TopCount NXT Microplate Scintillation & Luminescence Counter (PerkinElmer). All plates contained internal controls for full inhibition (3 mM glycine in radioligand solution) and full uptake (plain radioligand uptake buffer).

### NanoBRET experiments

Tracers were dissolved in DMSO at a final concentration of 1 mM. Inhibitors were dissolved in DMSO at a final concentration of 10 mM. 48 hours post-transfection, the media was removed, cells were trypsinized, re-suspended in DMEM and pelleted by centrifugation. The media was replaced with the appropriate assay buffer, cells were counted and the cell density was adjusted to 3x10^5^ cells/mL in assay buffer. Using a multichannel automatic pipette, 85 µL of cell suspension per well were seeded in a white 96-well OptiPlate (Revvity, 6055299). 10X working solutions of the desired inhibitors or DMSO vehicle (compared to the highest final concentration of 25 µM) were prepared by diluting the 10 mM DMSO stock solution in the appropriate assay buffer. 20X working solution of GlyT1-probe or DMSO vehicle (compared to the highest final concentration) was prepared in a tube by diluting the stock DMSO solution in tracer dilution buffer. *For direct binding experiments*, a template 96-well plate with 1:3 dilutions of the GlyT1-probe in tracer dilution buffer was prepared. *For competition binding experiments*, a template 96-well plate with 1:5 dilutions of the inhibitors (or vehicle) in the appropriate buffer was prepared. Using a multichannel pipette, 10 µL of the 10X working solution of inhibitor (or vehicle) was added to the cells, followed by the addition of 5 µL the 20X working solution of GlyT1-probe. The plate was shaken for 10 seconds and placed in the incubators for 2 hours. 10 minutes before reading, a detection mixture (Promega, N2160) composed of composed of NanoBRET Nano-Glo substrate (1:400) and extracellular NLuc inhibitor (1:1000) was prepared in the same buffer used for the assay. 50 µL of detection mixture was added to each well and the plate was orbitally shaken for 10 seconds at 300 rpm. Dual emission of each well was measured using a GloMax Discover (Promega) using emission filters of 450 nm (bandwidth 80 nm) for the donor and of 600 nm (long pass) for the acceptor, or an EnVision Multimode Plate Reader (PerkinElmer) using emission filters of 470 nm (bandwidth 24 nm) for the donor and of 590 nm (bandwidth 20 nm) for the acceptor.

### Pharmacological data analysis

All radioligand uptake experiments were performed in triplicate in at least three independent experiments. Data were normalized to maximal uptake after subtraction of the nonspecific uptake measured in the presence of 3 mM glycine. Concentration-response curves were fitted by nonlinear regression using the four-parameter concentration-response model:

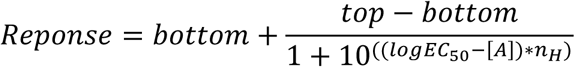

For concentration-inhibition experiments, IC_50_ values were determined by fitting concentration-response curves using the same four-parameter model. Data analysis was perfomed on pooled, normalized data using GraphPad Prism 11.0. For statistical analysis, a two-sided Welch’s t-test for comparison of mean between groups was used.

All NanoBRET experiments were performed in technical duplicate in at least three independent experiments. MilliBRET (mBRET) ratios were calculated as the ratio between acceptor and donor emission, multiplied for 1000. Corrected mBRET were calculated by background subtraction: the mBRET signal of untreated wells (no tracer) was removed from the mBRET value of each well. For competition assays, normalized BRET ratios were calculated as the corrected mBRET value of each compound-treated well (compound + tracer) divided for the corrected mBRET of vehicle-treated wells (tracer only). Data were analysed using the non-linear regression algorithms for dose-response (four parameters) in Prism 11.0.2 (GraphPad, USA). The IC_50_ values of inhibitors were converted to K_i_ values using the Cheng-Prusoff equation, and the Źfactor was calculated using the equation:

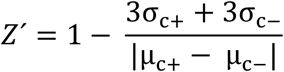

Where σ_c+_ and σ_c-_ are the standard deviation of the positive and negative control, and µ_c+_ and µ_c-_the means of the positive and negative control.

## ASSOCIATED CONTENT

### Supporting Information

The following files are available free of charge.

Supplementary figures 1-7, synthetic procedures, HPLC chromatograms and NMR spectra of selected compounds (PDF)

## AUTHOR INFORMATION

### Author Contributions

F.B. conceptualized and designed the study, coordinated the experiments, performed most of NanoBRET experiments, designed and synthesized some compounds. M.H.J. performed most of the uptake inhibition experiments. I.A. synthesized most of the compounds. M.H.J, I.A., and F.B wrote the first draft of the manuscript. A.M. Performed some uptake inhibition experiment. W.F., N.M.K. and J.S.H.M. contributed to the study design. N.M.K performed some NanoBRET experiments. J.S.H.M. designed plasmids, performed some uptake inhibition experiment. W.F., B.F., P.W. revised the manuscript. The manuscript was written through contributions of all authors. All authors have given approval to this version of the manuscript.

## ABBREVIATIONS

BRET: bioluminescence resonance energy transfer
DAD: diode array detector
DCM: dichloromethane
DMEM: Dulbecco’s modified Eagle medium
DMF: N,N-dimethylformamide
DMSO: dimethyl sulfoxide
DPBS: Dulbecco’s phosphate-buffered saline
ESI: electrospray ionization
FBS: fetal bovine serum
FC: flash chromatography
GABA: γ-aminobutyric acid
GA: gluconic acid
GAT1: γ-aminobutyric acid transporter 1 (SLC6A1)
GAT3: γ-aminobutyric acid transporter 3 (SLC6A11)
GlyT1: glycine transporter 1 (SLC6A9)
GPCR: G protein-coupled receptors
HBSS: Hanks’ balanced salt solution
HEK293T: human embryonic kidney 293T cells
HEPES: 4-(2-hydroxyethyl)-1-piperazineethanesulfonic acid
HPLC: high-performance liquid chromatography
HRMS: high-resolution mass spectrometry
LC-MS: liquid chromatography-mass spectrometry
MALDI: matrix-assisted laser desorption/ionization
mBRET: milliBRET units
NLuc: NanoLuciferase
NMDA: N-methyl-D-aspartate
NMDG: N-methyl-D-glucamine
NMR: nuclear magnetic resonance
Opti-MEM: reduced-serum cell culture medium
PDB: Protein Data Bank
PDL: poly-D-lysine
P/S: penicillin-streptomycin
RLU: relative luminescence units
RP: reverse phase
SLC: solute carrier
SLC6: solute carrier family 6
TFA: trifluoroacetic acid
THF: tetrahydrofuran
TLC: thin-layer chromatography
WT: wild type

