## Supplementary Information for "Development of a NanoBRET assay platform to detect intracellular target engagement in living cells for the Glycine Transporter 1 (SLC6A9)"

**Table of contents**


| | Title | Page |
|---|---|---|
| Supplementary Figure 1 | **Identification of the exit vector from the GlyT1 intracellular site.** | 2 |
| Supplementary Figure 2 | **Synthetic scheme for the synthesis of test probes *m*-1a to -1d and *p*-1a to -1d.** | 3 |
| Supplementary Figure 3 | **Preliminary screening of test probes for inhibition of [$^3$H]-Glycine uptake in HEK293T cells transiently expressing wild-type GlyT1.** | 5 |
| Supplementary Figure 4 | **Synthetic scheme for the synthesis of NanoBRET probes *m*-1e to *m*-1g.** | 6 |
| Supplementary Figure 5 | **Plasmid map of the GlyT1-NLuc.** | 7 |
| Supplementary Figure 6 | **Conserved apparent potency of glycine at GlyT1-nanoLuc in the absence and presence of 120 minutes preincubation.** | 8 |
| Supplementary Figure 7 | **Conserved inhibitory activity of 1-*m*-f (GlyT1-probe) at wild-type GlyT1.** | 9 |
| Supplementary Figures 8-21 | **HPLC traces for all final compounds.** | 10-15 |
| Synthetic procedures | **Synthesis of compounds** | 16-34 |

**Supplementary Figure 1. Identification of the exit vector from the GlyT1 intracellular site.** A) The intracellular pocket of GlyT1 (grey and green surface) where Cmpd1 (orange) binds is exposed to the cytosol, and delimited by the amino acids Y116, M382, N386 and D396 (green surface). B) Overlay of Cmpd1 (orange) with its simplified analog (pink) identifies the meta and para positions of the phenyl ring as suitable exit vectors from the GlyT1 intracellular site (illustrated as yellow arrows). GlyT1 backbone is displayed in faded green, with the four amino acids defining the cytosolic opening of the pocket in dark green.

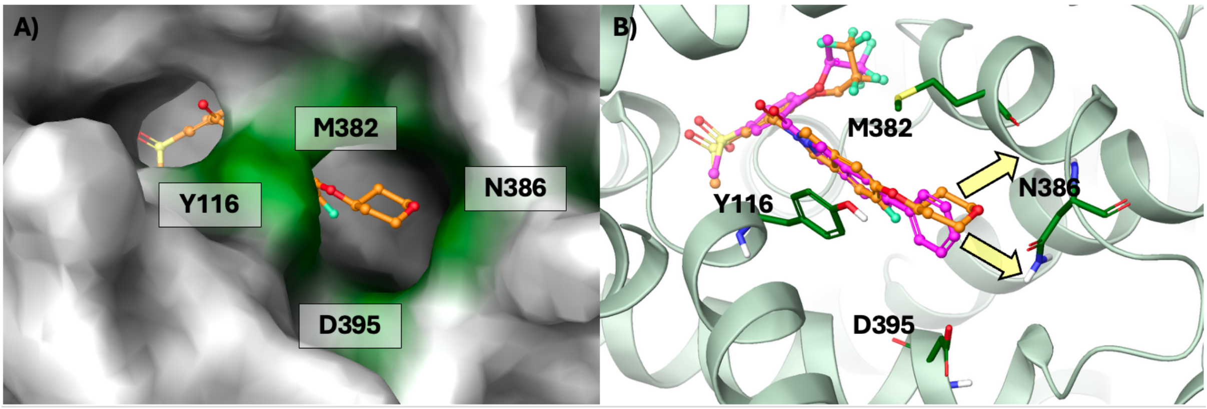

**Supplementary Figure 2. Synthetic scheme for the synthesis of test probes *m*-1a to -1d and *p*-1a to -1d**. $BBr_3$-mediated methyl ether hydrolysis of 2-methoxy-5-(methylsulfonyl)benzoate and methyl ester re-formation, followed by nucleophilic substitution with 2,2,3,3,3-pentafluoropropyl trifluoromethanesulfonate and subsequent methyl ester hydrolysis led to intermediate compound **3**. Suzuki-Miyaura cross coupling of *m*- or *p*-substituted aryl boronic acids with *tert*-butyl 5-bromoisoindoline-2-carboxylate led to compounds ***m*-4a** to ***m*-4d** and ***p*-4a** to ***m*-4c**, which upon *N*-Boc deprotection and amide coupling with **3** gave the desired GlyT1 probes ***m*-1a** to **-d** and ***p*-1a** to **-c**. In a modified version of the scheme, initial *N*-Boc deprotection of *tert*-butyl 5-bromoisoindoline-2-carboxylate, followed by amide coupling with **3** and subsequent Suzuki-Miyaura cross coupling, gave the GlyT1 probe ***p*-1d**.

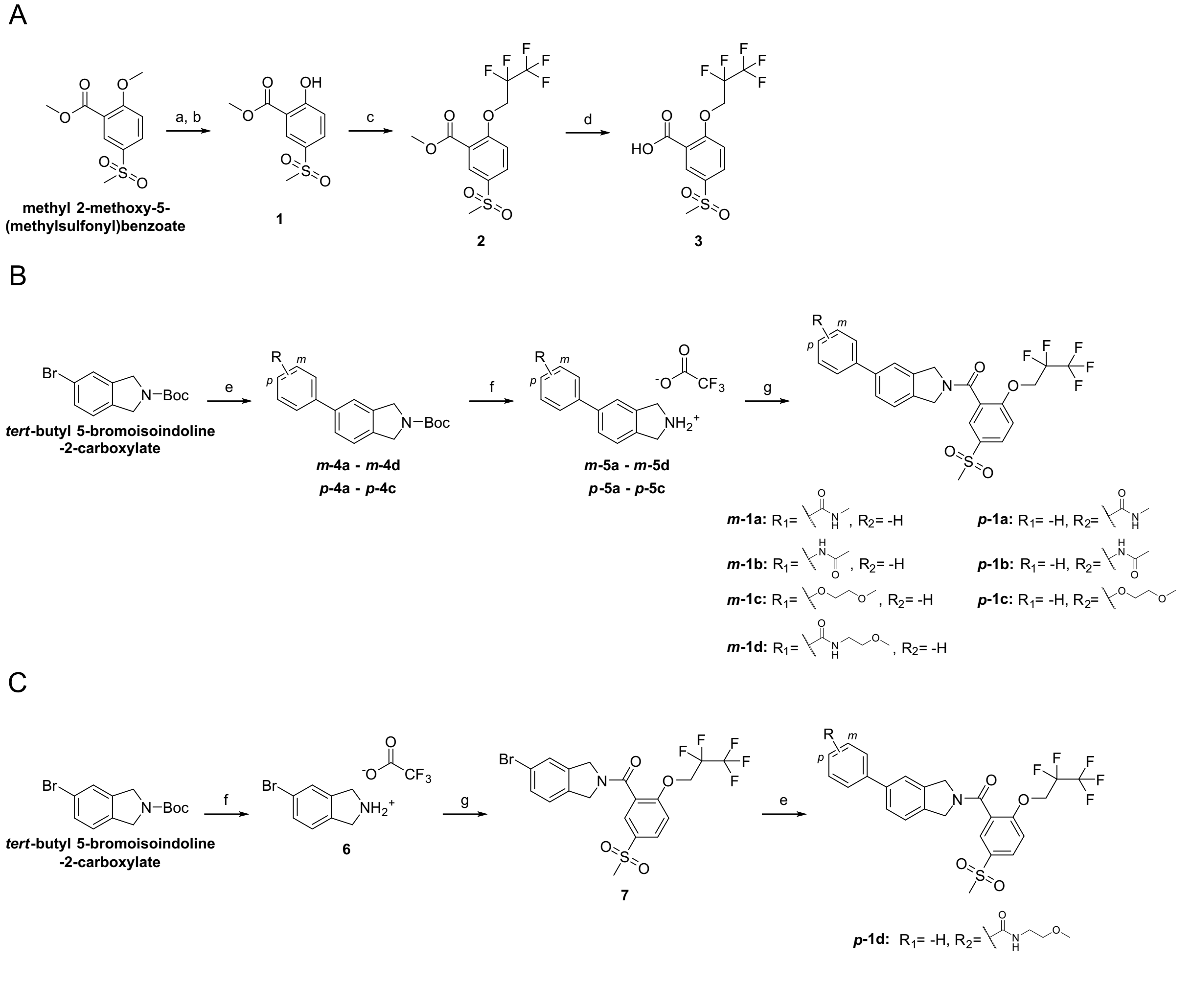


Reactions and conditions: **a)** 1 M $BBr_3$ in DCM, anhydrous DCM, -78 °C–rt, 18 h; **b)** conc. $H_2SO_4$, trimethyl orthoformate, anhydrous MeOH, reflux, 20 h, 93% over two steps; **c)** 2,2,3,3,3-pentafluoropropyl trifluoromethanesulfonate, $K_2CO_3$, butan-2-one, reflux, 42 h, 62%; **d)** LiOH·H2O, THF/$H_2O$ 1:1, rt, 1 h, 95%; **e)** HCl 4M in dioxane, or neat TFA, or TFA in $CHCl_3$/MeOH 1:1, 0–60 °C, 2-42 h, 30–87%; **f)** $Cs_2CO_3$, $Pd(PPh_3)_4$, 1,4- Dioxane/$H_2O$/EtOH 1:1:1 or DME/ $H_2O$ 1:1 (degassed), 50–80 °C, 18 h, 23–87%; **g) 3**, HBTU, DIPEA, anhydrous DMF, 18 h, rt, 7–92%.

**Supplementary Figure 3. Preliminary screening of test probes for inhibition of [$^3$H]-Glycine uptake in HEK293T cells transiently expressing wild-type GlyT1.** Compounds were tested at 10, 100 and 1000 nM following 60 minutes of preincubation. Data are normalized to [3H]-Glycine uptake in the absence of inhibitor and presented as mean residual uptake from n=3 (panel A) or n=2 (panel B) independent experiments. Error bars represent SEM (panel A) or SD (panel B). Bitopertin was included as a reference inhibitor.

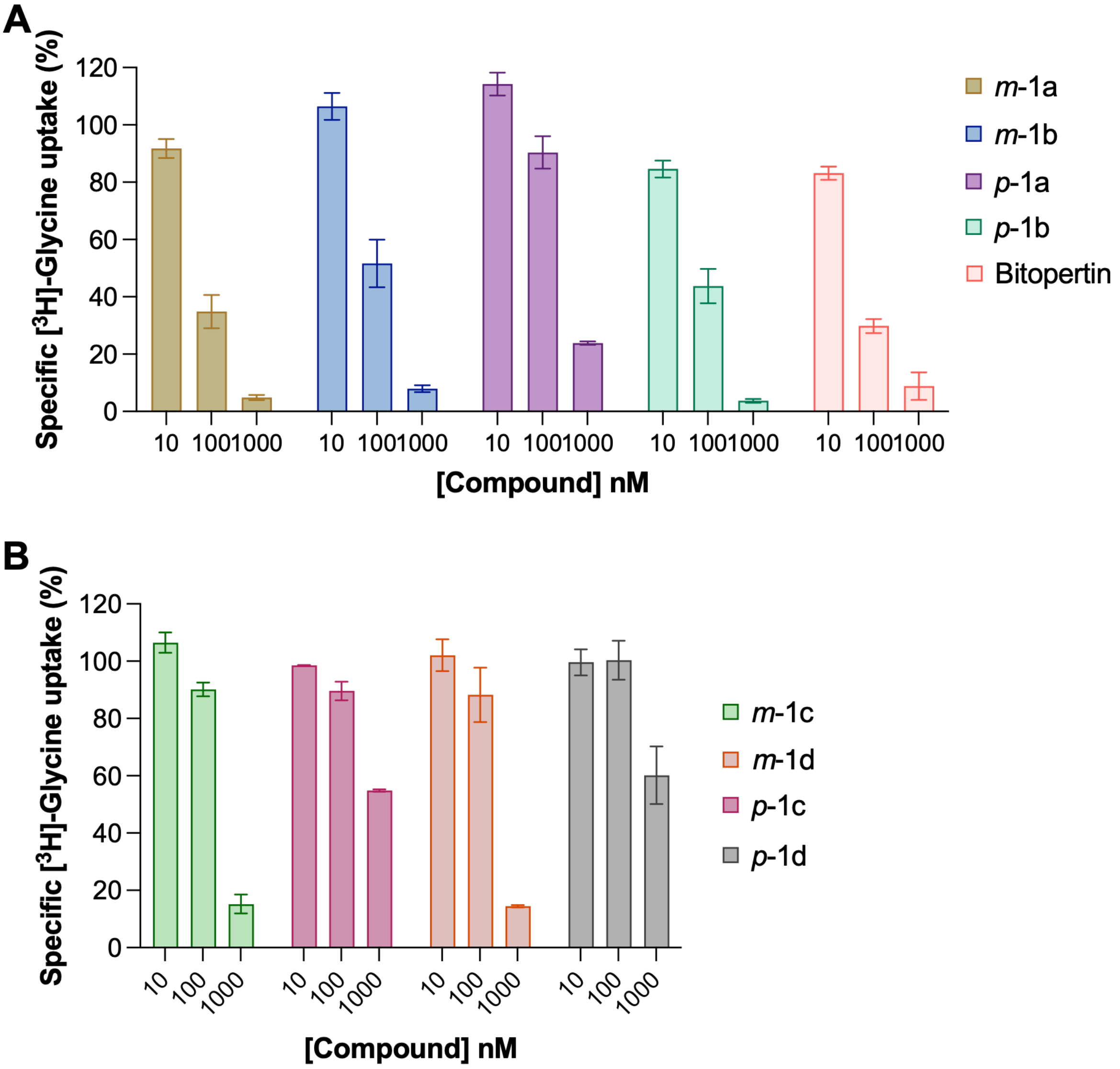

**Supplementary Figure 4. Synthetic scheme for the synthesis of NanoBRET probes *m*-1e to *m*-1g.** Suzuki-Miyaura cross coupling of intermediate **7** with (3-bromophenyl)boronic acid provided **8**, that was coupled with propargyl amine, pent-4-yn-1-amine or 2-(2-(prop-2-yn-1-yloxy)ethoxy)ethan-1-amine to give the intermediates ***m*-9e** to **-g** respectively. These intermediates feature an alkyne handle attached to the benzamide moiety with spacers of increasing length and were conjugated via copper-catalyzed azide–alkyne cycloaddition (CuAAC) to the azide-containing BODIPY-590 providing the NanoBRET probes ***m*-1e** to ***m*-1g**.

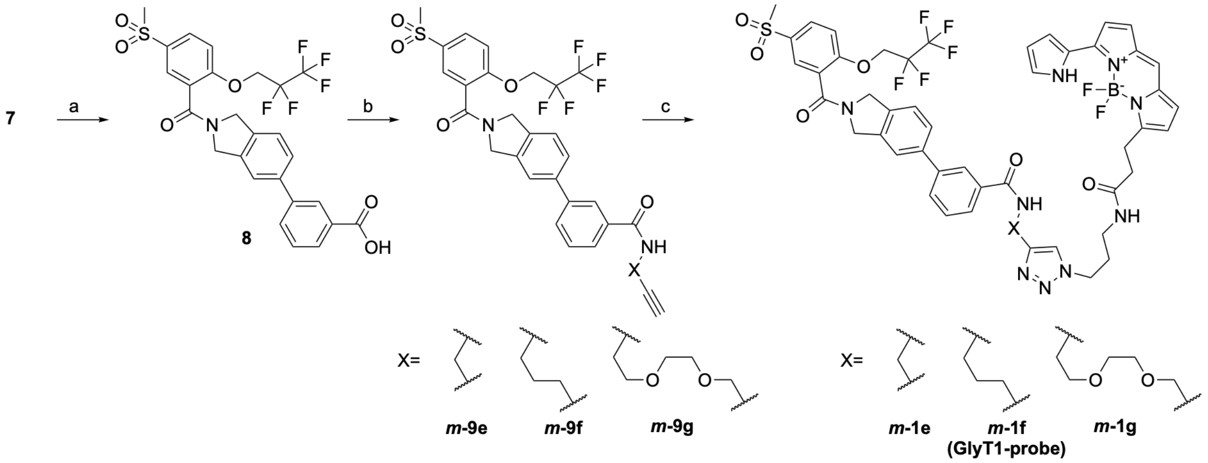


Reactions and conditions: **a)** $Cs_2CO_3$, $Pd(PPh_3)_4$, 1,4- Dioxane/$H_2O$/EtOH 1:1:1 (degassed), 50 °C, 18 h, 79%; **b)** HBTU, *N*-ethyl-diisopropylamine, anhydrous DMF, rt, 20 min; then propargyl-amine (for ***m*-9e**) or pent-4-yn-1-amine (for ***m*-9f**) or 2-(2-(prop-2-yn-1-yloxy)ethoxy)ethan-1-amine (for ***m*-9g**), 18 h, rt, 61-80%; **c)** BODIPY 590-azide in degassed DMF, Tetrakis(acetonitrile)copper(I) triflate and 3,3',3''-((nitrilotris(methylene))tris(1H-1,2,3-triazole-4,1-diyl))tris(propan-1-ol) in degassed milliQ water, 18h, rt, 82-91%.

**Supplementary figure 5. Plasmid map of the GlyT1-NLuc.** pBiT3.1-N-CMV vector was modified to carry the sequence of GlyT1 followed by a linker (EFAGSSGGSSG) and the coding sequence for NLuc.

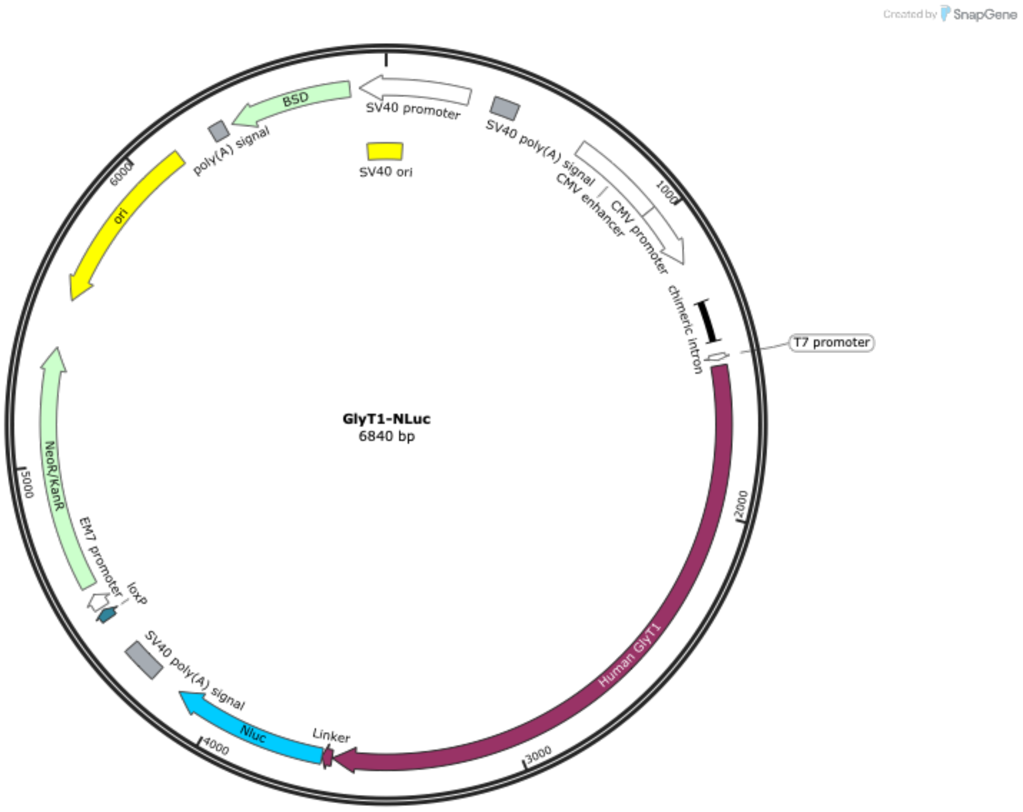

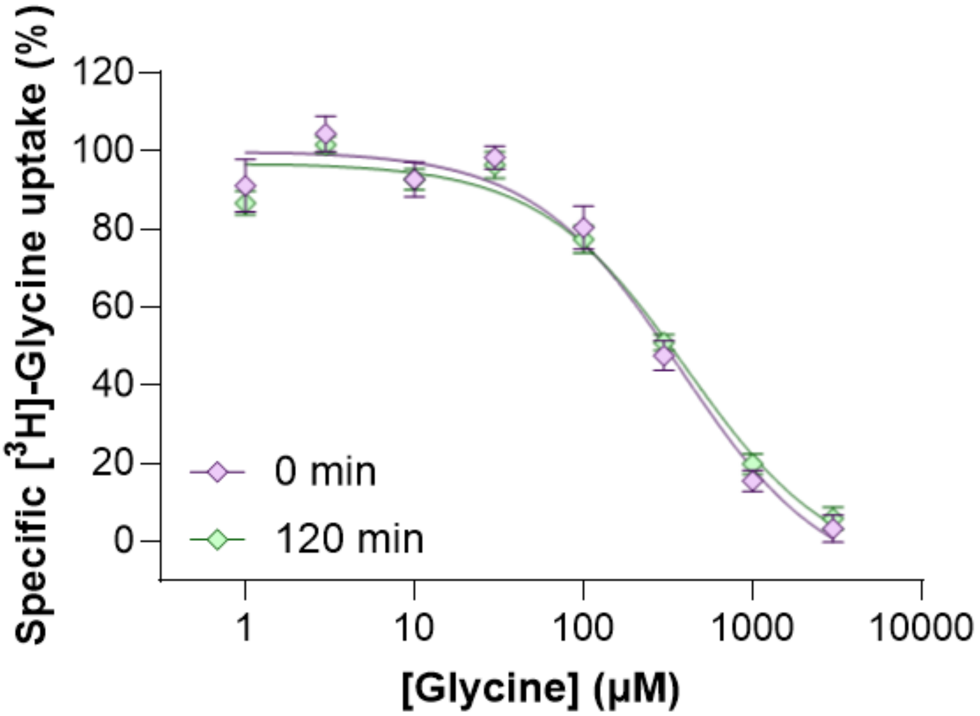


**Supplementary figure 6. Conserved apparent potency of glycine at GlyT1-nanoLuc in the absence and presence of 120 minutes preincubation.** [$^3$H]-glycine uptake in HEK293T cells transiently expressing GlyT1-NLuc. Data are normalized to maximal [$^3$H]-glycine uptake and presented as mean ± SEM of residual [$^3$H]-glycine uptake from $n$ = 3-4 biological replicates. $pIC_{50}$ values for glycine potency in the absence ($IC_{50}$: $pIC_{50}$ ± SEM, *n*; 299.4 µM: 3.53 ± 0.03, *4)* and presence of 120 minutes preincubation ($IC_{50}$: $pIC_{50}$ ± SEM, *n*; 328.8 µM: 3.49 ± 0.04, *4)* were not statistically different ($p$=0.4121, two-sided Welch's t-test)

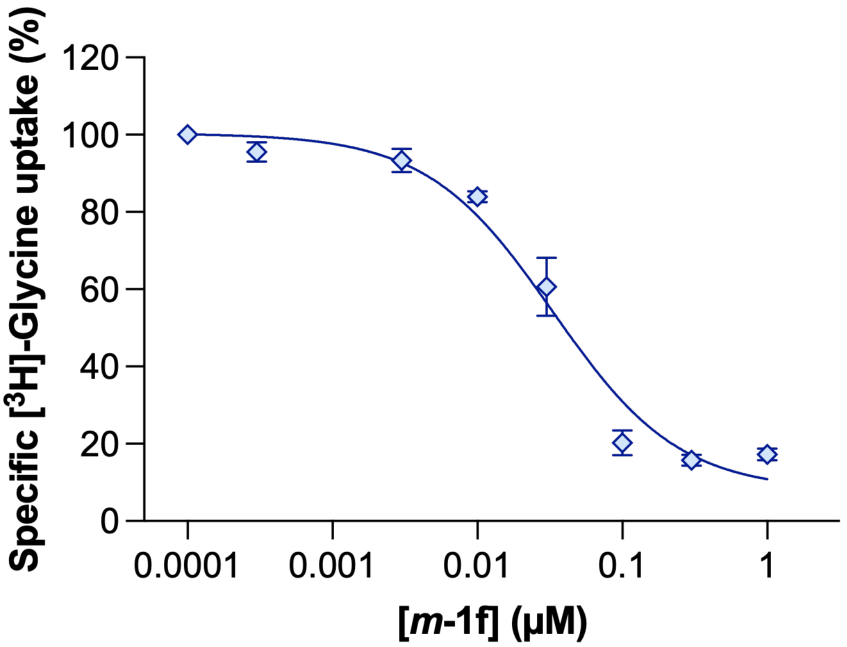


**Supplementary Figure 7**. **Conserved inhibitory activity of *m*-1f (GlyT1-probe) at wild-type GlyT1.** [$^3$H]-glycine uptake in HEK293T cells transiently expressing GlyT1-WT. Data were normalized to maximal [$^3$H]-glycine uptake and presented as mean ± SEM of residual [$^3$H]-glycine uptake from $n$ = 3 biological replicates. Resulting potency is summarized in table 2.

**Supplementary Figure 8**. **HPLC trace of compound *m*-1a.**

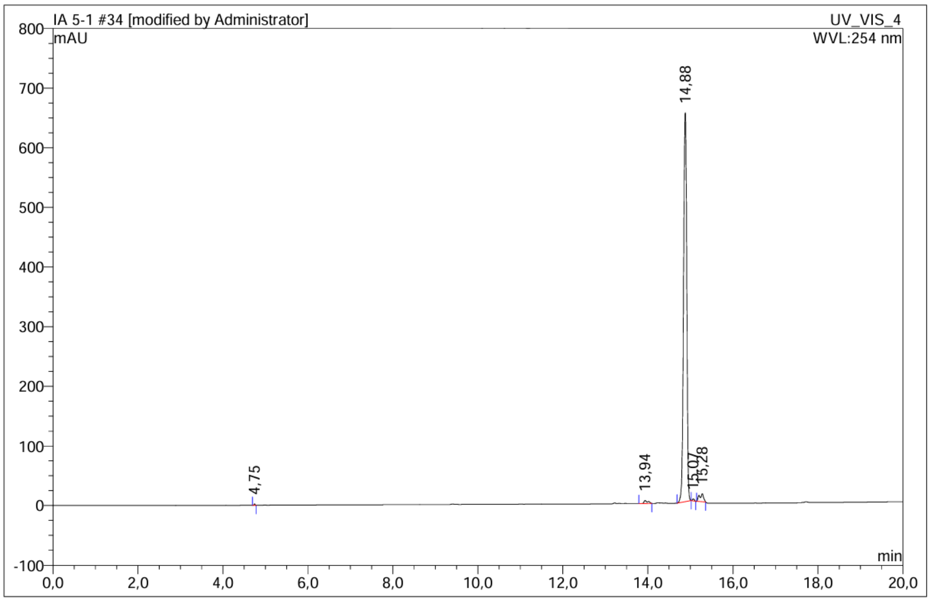


| No. | Ret.Time min | Peak Name | Height mAU | Area mAU*min | Rel.Area % | Amount | Resolution(EP) |
|---|---|---|---|---|---|---|---|
| 1 | 4,75 | n.a. | 2,579 | 0,064 | 0,11 | n.a. | 112,81 |
| 2 | 13,94 | n.a. | 5,094 | 0,540 | 0,88 | n.a. | 7,05 |
| 3 | 14,88 | n.a. | 652,406 | 58,952 | 96,27 | n.a. | 1,58 |
| 4 | 15,07 | n.a. | 3,527 | 0,192 | 0,31 | n.a. | 1,35 |
| 5 | 15,28 | n.a. | 13,585 | 1,490 | 2,43 | n.a. | n.a. |
| Total: | | | 677,191 | 61,238 | 100,00 | 0,000 | |

**Supplementary Figure 9**. **HPLC trace of compound *m*-1b.**

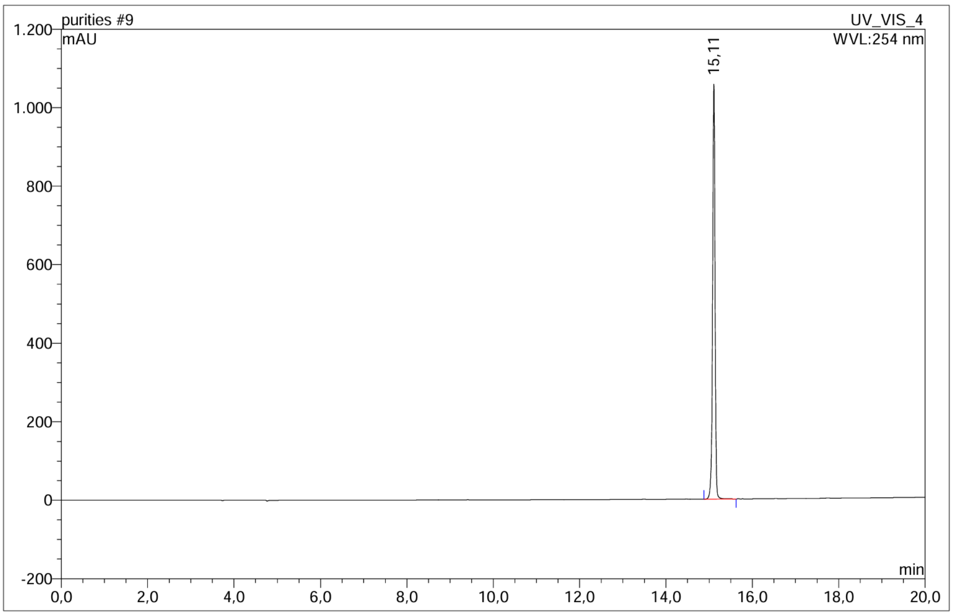


| No. | Ret.Time min | Peak Name | Height mAU | Area mAU*min | Rel.Area % | Amount | Resolution(EP) |
|---|---|---|---|---|---|---|---|
| 1 | 15,11 | n.a. | 1056,924 | 70,895 | 100,00 | n.a. | n.a. |
| Total: | | | 1056,924 | 70,895 | 100,00 | 0,000 | |

**Supplementary Figure 10. HPLC trace of compound *m*-1c.**

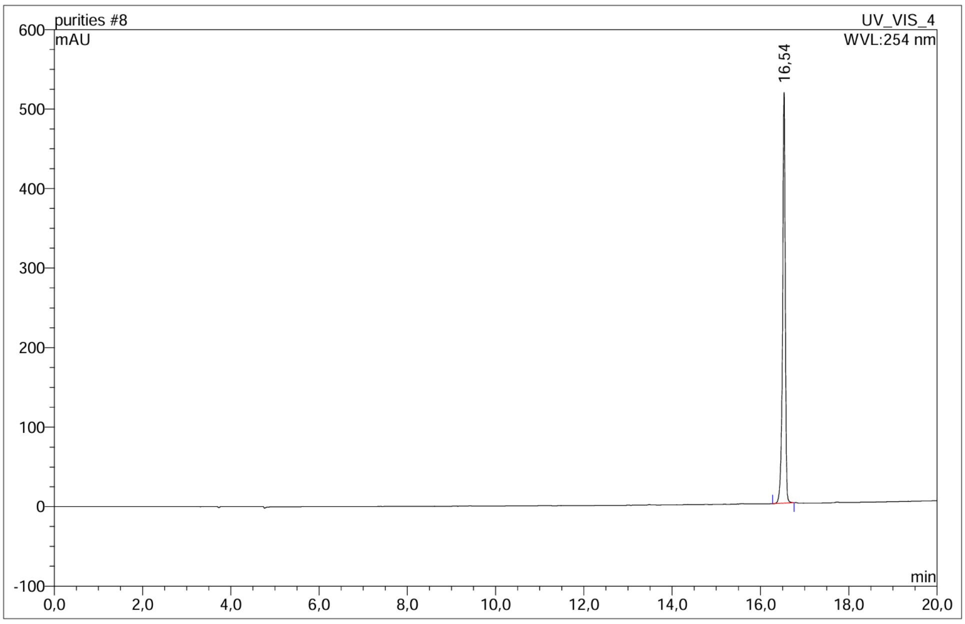


| No. | Ret.Time min | Peak Name | Height mAU | Area mAU*min | Rel.Area % | Amount | Resolution(EP) |
|---|---|---|---|---|---|---|---|
| 1 | 16,54 | n.a. | 516,783 | 36,795 | 100,00 | n.a. | n.a. |
| Total: | | | 516,783 | 36,795 | 100,00 | 0,000 | |

**Supplementary Figure 11. HPLC trace of compound *m*-1d.**

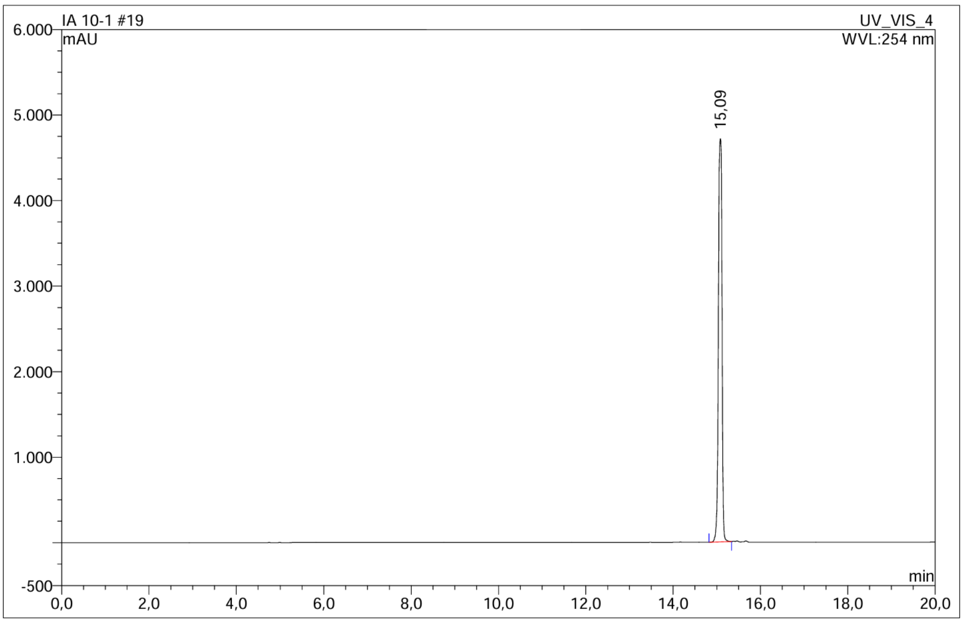


| No. | Ret.Time min | Peak Name | Height mAU | Area mAU*min | Rel.Area % | Amount | Resolution(EP) |
|---|---|---|---|---|---|---|---|
| 1 | 15,09 | n.a. | 4714,947 | 460,391 | 100,00 | n.a. | n.a. |
| Total: | | | 4714,947 | 460,391 | 100,00 | 0,000 | |

**Supplementary Figure 12**. **HPLC trace of compound *p*-1a.**

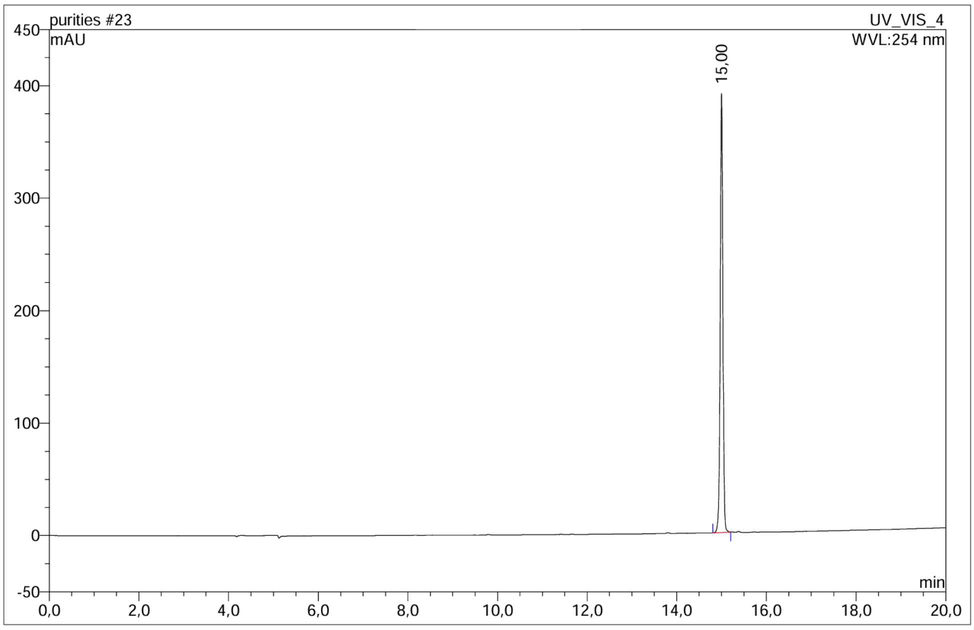


| No. | Ret.Time min | Peak Name | Height mAU | Area mAU*min | Rel.Area % | Amount | Resolution(EP) |
|---|---|---|---|---|---|---|---|
| 1 | 15,00 | n.a. | 390,522 | 26,743 | 100,00 | n.a. | n.a. |
| Total: | | | 390,522 | 26,743 | 100,00 | 0,000 | |

**Supplementary Figure 13**. **HPLC trace of compound *p*-1b.**

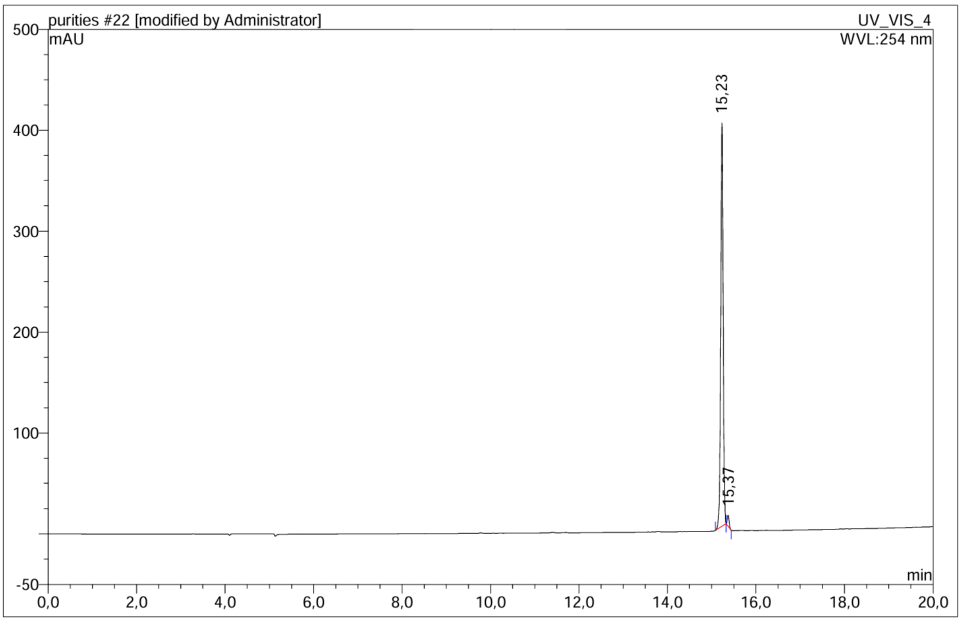


| No. | Ret.Time min | Peak Name | Height mAU | Area mAU*min | Rel.Area % | Amount | Resolution(EP) |
|---|---|---|---|---|---|---|---|
| 1 | 15,23 | n.a. | 399,877 | 25,633 | 97,98 | n.a. | 1,51 |
| 2 | 15,37 | n.a. | 10,837 | 0,529 | 2,02 | n.a. | n.a. |
| Total: | | | 410,714 | 26,162 | 100,00 | 0,000 | |

**Supplementary Figure 14**. **HPLC trace of compound *p*-1c.**

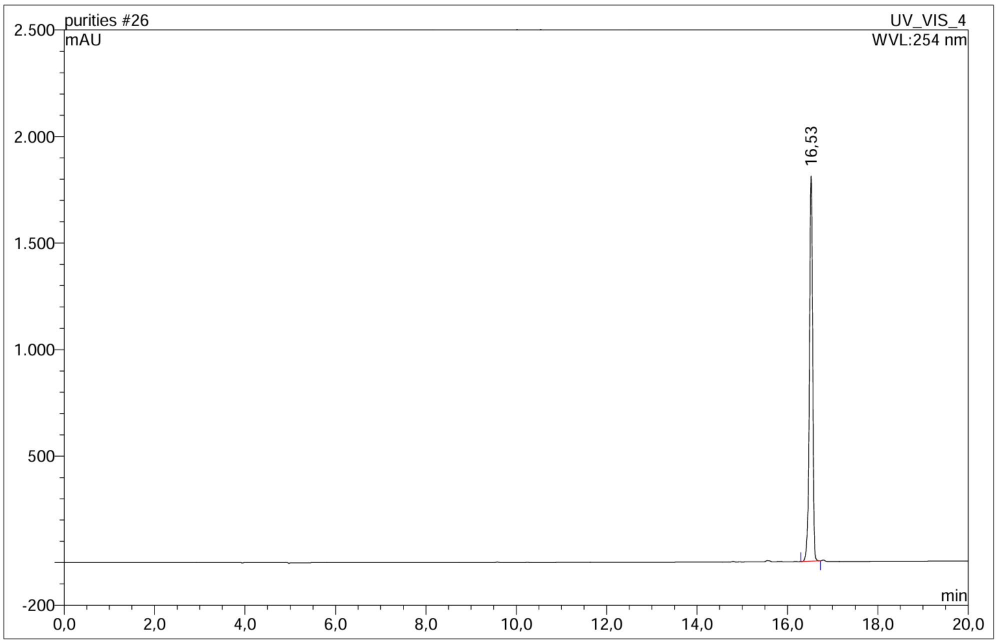


| No. | Ret.Time min | Peak Name | Height mAU | Area mAU*min | Rel.Area % | Amount | Resolution(EP) |
|---|---|---|---|---|---|---|---|
| 1 | 16,53 | n.a. | 1807,582 | 153,635 | 100,00 | n.a. | n.a. |
| Total: | | | 1807,582 | 153,635 | 100,00 | 0,000 | |

**Supplementary Figure 15**. **HPLC trace of compound *p*-1d.**

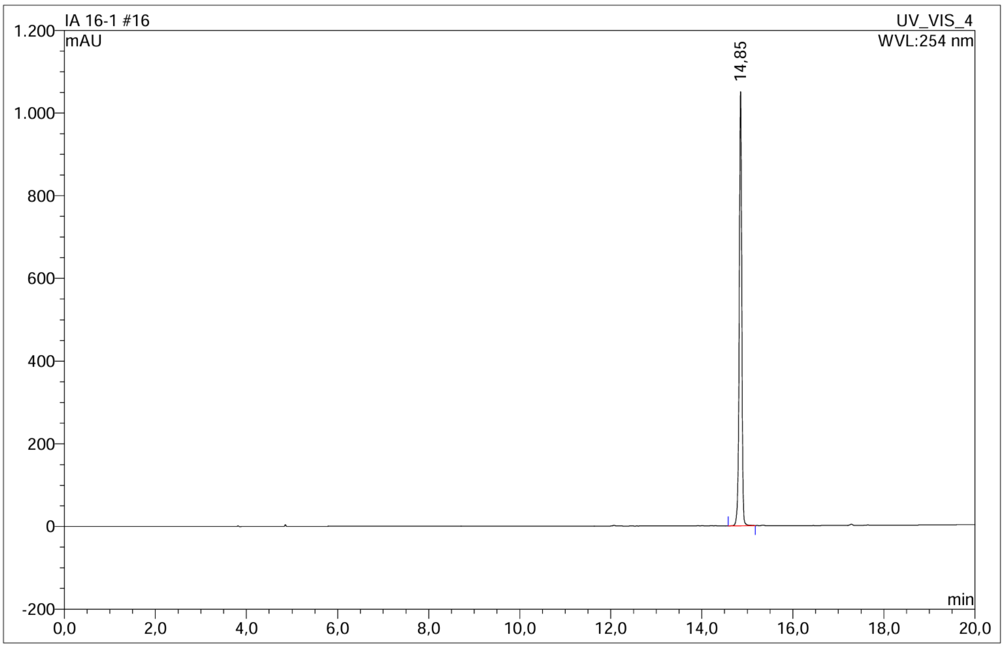


| No. | Ret.Time min | Peak Name | Height mAU | Area mAU*min | Rel.Area % | Amount | Resolution(EP) |
|---|---|---|---|---|---|---|---|
| 1 | 14,85 | n.a. | 1050,324 | 69,746 | 100,00 | n.a. | n.a. |
| Total: | | | 1050,324 | 69,746 | 100,00 | 0,000 | |

**Supplementary Figure 16**. **HPLC trace of compound 1-*m*-e (254 nm).**

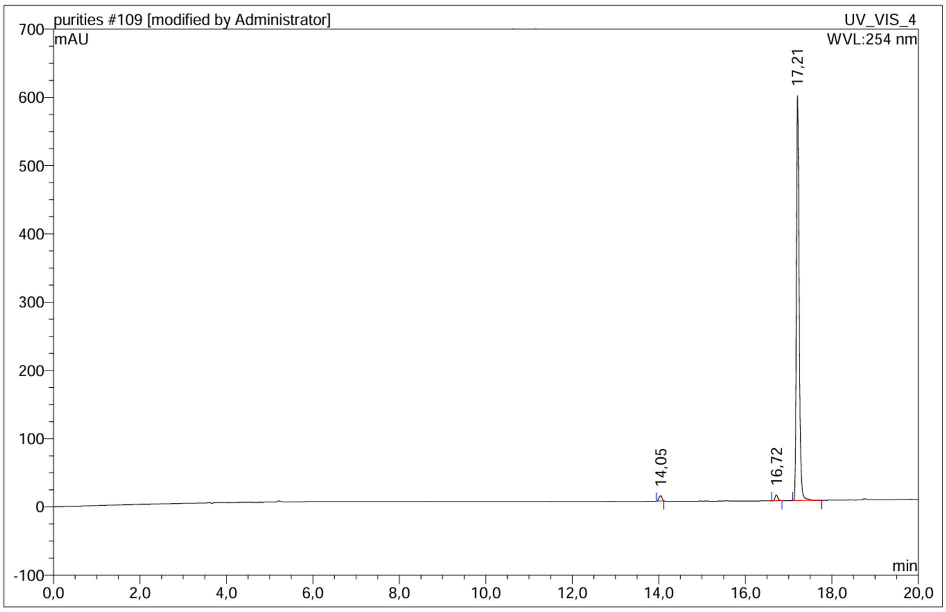


| No. | Ret.Time min | Peak Name | Height mAU | Area mAU*min | Rel.Area % | Amount | Resolution(EP) |
|---|---|---|---|---|---|---|---|
| 1 | 14,05 | n.a. | 7,617 | 0,603 | 1,40 | n.a. | 22,40 |
| 2 | 16,72 | n.a. | 8,480 | 0,589 | 1,37 | n.a. | 4,68 |
| 3 | 17,21 | n.a. | 593,681 | 41,815 | 97,23 | n.a. | n.a. |
| Total: | | | 609,778 | 43,007 | 100,00 | 0,000 | |

**Supplementary Figure 17**. **HPLC trace of compound 1-*m*-e (575 nm).**

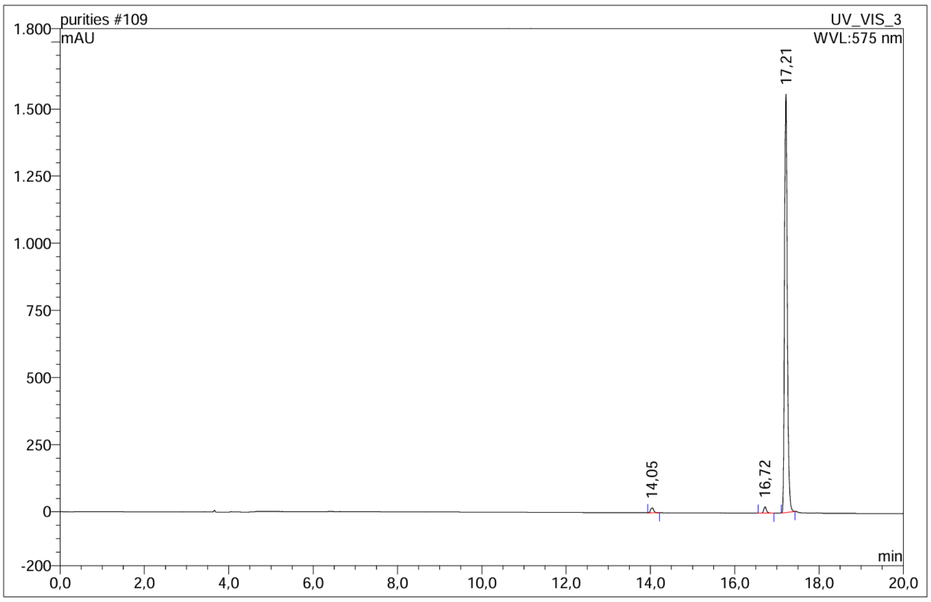


| No. | Ret.Time min | Peak Name | Height mAU | Area mAU*min | Rel.Area % | Amount | Resolution(EP) |
|---|---|---|---|---|---|---|---|
| 1 | 14,05 | n.a. | 18,147 | 1,573 | 1,34 | n.a. | 21,83 |
| 2 | 16,72 | n.a. | 23,720 | 1,710 | 1,45 | n.a. | 4,52 |
| 3 | 17,21 | n.a. | 1558,971 | 114,412 | 97,21 | n.a. | n.a. |
| Total: | | | 1600,838 | 117,695 | 100,00 | 0,000 | |

**Supplementary Figure 18. HPLC trace of compound 1-*m*-f (254 nm).**

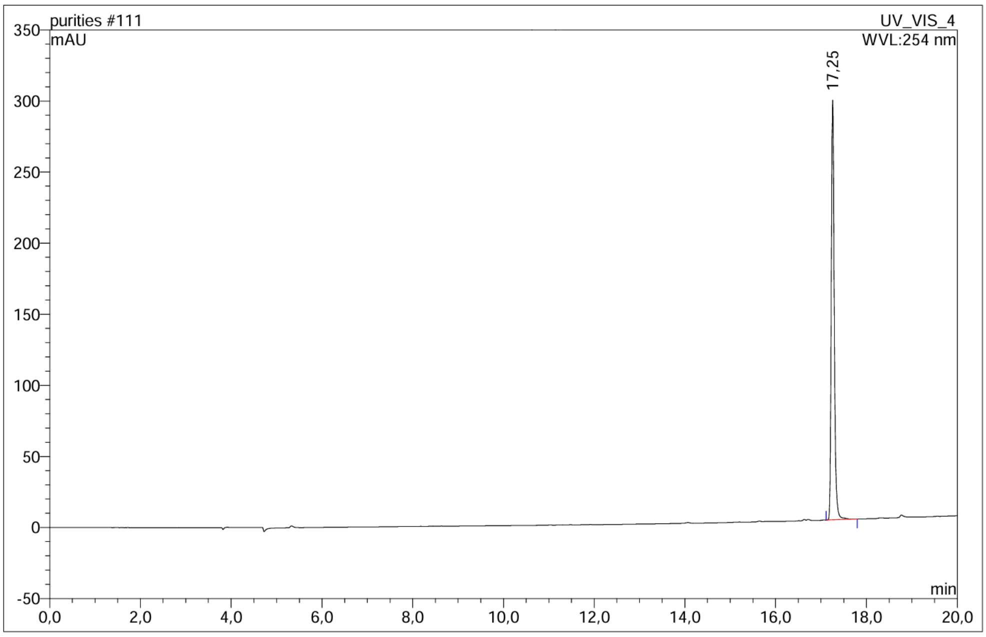


| No. | Ret.Time min | Peak Name | Height mAU | Area mAU*min | Rel.Area % | Amount | Resolution(EP) |
|---|---|---|---|---|---|---|---|
| 1 | 17,25 | n.a. | 295,291 | 21,651 | 100,00 | n.a. | n.a. |
| Total: | | | 295,291 | 21,651 | 100,00 | 0,000 | |

**Supplementary Figure 19. HPLC trace of compound 1-*m*-f (575 nm).**

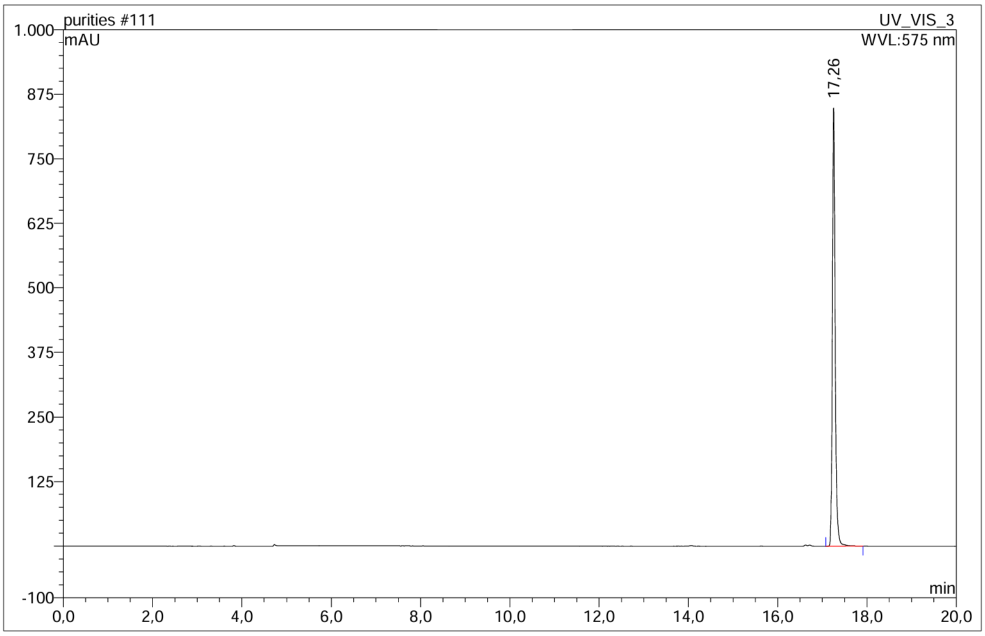


| No. | Ret.Time min | Peak Name | Height mAU | Area mAU*min | Rel.Area % | Amount | Resolution(EP) |
|---|---|---|---|---|---|---|---|
| 1 | 17,26 | n.a. | 848,778 | 62,574 | 100,00 | n.a. | n.a. |
| Total: | | | 848,778 | 62,574 | 100,00 | 0,000 | |

**Supplementary Figure 20. HPLC trace of compound 1-*m*-g (254 nm).**

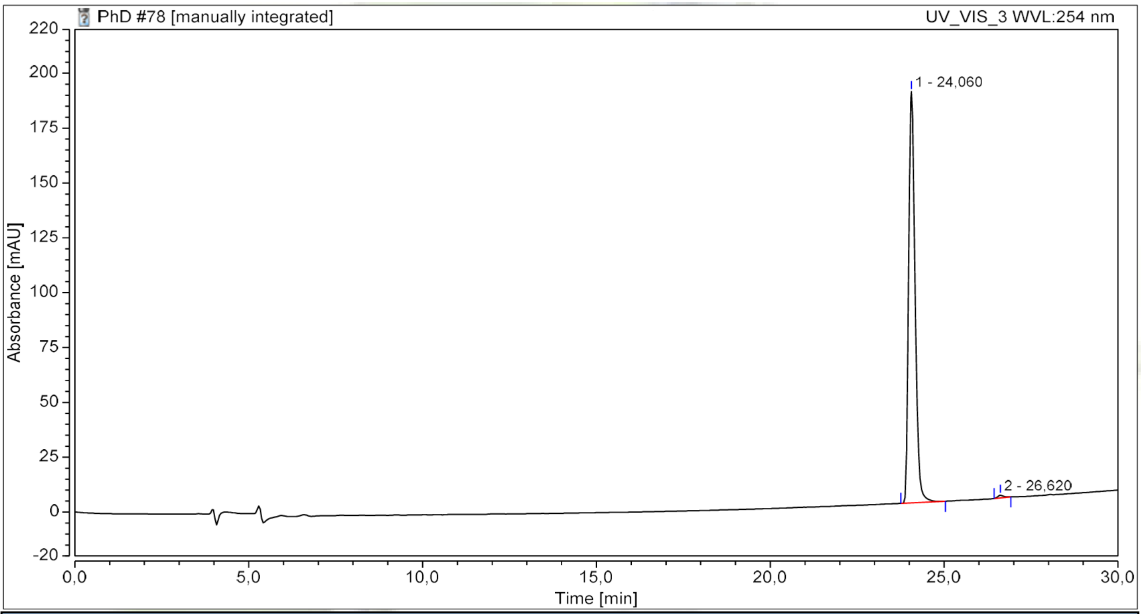


| Integration Results | | | | | | | |
|---|---|---|---|---|---|---|---|
| No. | Peak Name | Retention Time min | Area mAU*min | Height mAU | Resolution (EP) | Rel.Area % | Amount n.a. |
| 1 | | 24,060 | 40,133 | 187,446 | 8,05 | 99,42 | n.a. |
| 2 | | 26,620 | 0,235 | 1,188 | n.a. | 0,58 | n.a. |
| **Total:** | | | **40,368** | **188,633** | **8,05** | **100,00** | |

**Supplementary Figure 21. HPLC trace of compound 1-*m*-g (550 nm).**

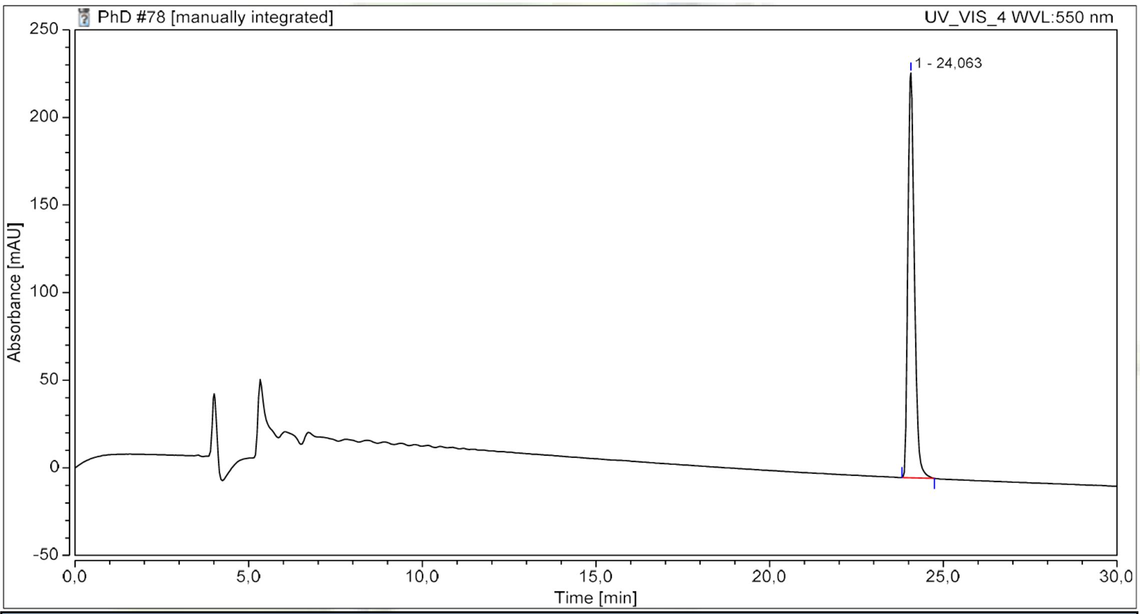


| Integration Results | | | | | | | |
|---|---|---|---|---|---|---|---|
| No. | Peak Name | Retention Time min | Area mAU*min | Height mAU | Resolution (EP) | Rel.Area % | Amount n.a. |
| 1 | | 24,063 | 50,036 | 231,016 | n.a. | 100,00 | n.a. |
| **Total:** | | | **50,036** | **231,016** | **0,00** | **100,00** | |

**Synthesis of compounds**

*Methyl 2-hydroxy-5-(methylsulfonyl)benzoate (**1**)*

In a flame-dried flask containing a dry-ice-cooled solution of methyl 2-(methyloxy)-5 (methylsulfonyl)benzoate (5 g, 20 mmol, 1 eq.) in dichloromethane (40 mL) under nitrogen atmosphere, boron tribromide (7.692 g, 30 mmol, 1.5 eq.) was added dropwise. Upon stirring overnight, ice was poured into the solution. Upon completion of ice melting, the two phases were transferred to a separatory funnel and the aqueous phase was extracted with EtOAc (x4). Combined organic phases were dried over anhydrous $Na_2SO_4$, filtered and the solvent was removed *in vacuo*. The residue was transferred to a flame-dried flask under inert conditions, dissolved in methanol (75 mL), sulphuric acid (5 mL) and trimethyl orthoformate (10 mL) and stirred under reflux for 20 hours. Upon completion, the resulting solution was cooled, evaporated, dissolved in EtOAc/$H_2O$ and the organic phase was washed with saturated aqueous $K_2CO_3$ solution, dried over $Na_2SO_4$, filtered and dried *in vacuo* to obtain the title compound as a white solid (4.28 g, 93%). LC-MS (ESI) $R_t$=3.42 min, m/z calculated for $C_9H_{11}O_5S^+$ $[M + H]^+$ = 231.03, found 231.0. $^1H$ NMR (600 MHz, $CDCl_3$) δ 11.31 (s, 1H), 8.47 (d, *J* = 2.4 Hz, 1H), 7.98 (dd, *J* = 8.8, 2.4 Hz, 1H), 7.14 (d, *J* = 8.8 Hz, 1H), 4.00 (s, 3H), 3.05 (s, 3H). $^{13}C$ NMR (151 MHz, $CDCl_3$) δ 169.6, 165.6, 134.3, 131.5, 130.9, 119.2, 112.8, 53.1, 44.9.

*Methyl 5-(methylsulfonyl)-2-(2,2,3,3,3-pentafluoropropoxy)benzoate (**2**)*

In a flame-dried flask equipped with a magnetic stirrer under inert conditions, methyl 2-hydroxy 5-(methylsulfonyl)benzoate (4.50 g, 19.5 mmol, 1 eq.), 2,2,3,3,3-pentafluoropropyl trifluoromethanesulfonate (6.06 g, 21.5 mmol, 1.1 eq.), $K_2CO_3$ (5.4 g, 39 mmol, 2 eq.) and 2-butanone (65mL) were added. The reaction was stirred under reflux for 2 days. Upon completion, the reaction mixture was concentrated *in vacuo* and the residue was dissolved in EtOAc. The solution was transferred to a separatory funnel and washed with saturated aqueous $K_2CO_3$ solution. The organic phase was dried over anhydrous $Na_2SO_4$, filtered and the solvent was removed *in vacuo*. The residue was purified by automated silica gel flash column chromatography using a gradient elution (*n*-Heptane/EtOAc). The title compound was obtained as a white solid (4.38 g, 62%). LC-MS (ESI) $R_t$=4.15 min, m/z calculated for $C_{12}H_{12}F_5O_5S^+$ $[M + H]^+$ = 363.03, found 363.1. $^1H$ NMR (400 MHz, $CDCl_3$) δ 8.35 (d, *J* = 2.4 Hz, 1H), 8.01 (dd, *J* = 8.8, 2.4 Hz, 1H), 7.05 (d, *J* = 8.8 Hz, 1H), 4.50 (t, *J* = 11.8 Hz, 2H), 3.86 (s, 3H), 3.00 (s, 3H). $^{13}C$ NMR (151 MHz, $CDCl_3$) δ 164.6, 160.1, 134.9, 122.9, 121.7, 121.4, 121.4, 121.2, 119.8, 119.5, 119.3, 117.9, 117.6, 117.4, 116.0, 115.7, 115.5, 114.3, 114.0, 113.8, 113.5, 112.6, 112.3, 112.1, 111.8, 110.9, 110.6, 110.4, 110.1, 66.4, 66.2, 66.0. $^{19}F$ NMR (376 MHz, $CDCl_3$) δ -83.3, -122.9.

*5-(methylsulfonyl)-2-(2,2,3,3,3-pentafluoropropoxy)benzoic acid **(3)***

In a flask equipped with magnetic stirrer contraining a solution 5-methanesulfonyl-2-(2,2,3,3,3-pentafluoro-propoxy)-benzoic acid methyl ester (4300 mg, 11.87 mmol, 1 eq.) in THF (55 mL), a solution of lithium hydroxide monohydrate (747 mg, 17.8 mmol, 1.5 eq.) in water (55 mL) was added. The reaction was stirred at room temperature for 1 hour. Upon completion, 17.5 mL of 1M aqueous hydrochloric acid solution were added dropwise at 0ºC. The reaction mixture was then concentrated *in vacuo* and the resulting precipitate was then filtered and washed with water several times to afford the title compound as a white solid (3924 mg, 95%). LC-MS (ESI) $R_t$ = 3.57 min, m/z calculated for $C_{11}H_{10}F_5O_5S^+$ $[M + H]^+$ = 349.02, found 349.0. $^1H$ NMR (400 MHz, DMSO) δ 13.29 (s, 1H), 8.17 (d, *J* = 2.4 Hz, 1H), 8.08 (dd, *J* = 8.8, 2.5 Hz, 1H), 7.52 (d, *J* = 8.9 Hz, 1H), 5.05 (t, *J* = 12.9 Hz, 2H), 3.24 (s, 3H). $^{13}C$ NMR (101 MHz, DMSO) δ 165.5, 158.7, 134.0, 131.8, 129.9, 122.8, 114.9, 64.9, 64.6, 64.3, 43.6. $^{19}F$ NMR (376 MHz, DMSO) δ -82.4, -122.2

*tert-butyl 5-(3-(methylcarbamoyl)phenyl)isoindoline-2-carboxylate **(m-4a)***

In a flame-dried microwave vial equipped with a magnetic stirring bar under inert atmosphere, *tert*-butyl 5-bromoisoindoline-2 carboxylate (200 mg, 0.67 mmol, 1 eq.), (3-(methylcarbamoyl)phenyl)boronic acid (143 mg, 0.80 mmol, 1.2 eq.), $Cs_2CO_3$ (873 mg, 2.68 mmol, 4 eq.) and tetrakis(triphenylphosphine)palladium (23.1 mg, 0.02 mmol, 0.03 eq.) were added. Degassed DME/$H_2O$ 1:1 was added and the reaction was left stirring overnight at 80ºC. Upon completion, the reaction mixture was diluted with EtOAc and aqueous HCl 1M was added dropwise until pH 4-5. The two phases are transferred in a separatory funnel and the aqueous phase was extracted with EtOAc. The combined organic layers were dried over anhydrous $Na_2SO_4$ and reduced *in vacuo*. Purification of the residue by automated siliga gel flash column chromatography using a gradient elution (*n*-Heptane/EtOAc), followed by automated C18 flash column chromatography using a gradient elution, yielded the product as a yellow oil (54 mg, 23%). LC-MS (ESI) $R_t$ = 4.54 min, m/z calculated for $C_{21}H_{25}N_2O_3^+$ $[M + H]^+$ = 353.2 found 353.2. $^1H$ NMR (600 MHz, $CDCl_3$) δ 7.98 (s, 1H), 7.72 – 7.67 (m, 2H), 7.53 – 7.46 (m, 3H), 7.34 – 7.30 (m, 1H), 6.23 (s, 1H), 4.72 (m, 4H), 3.05 (d, *J* = 4.3 Hz, 3H), 1.53 (s, 9H). $^{13}C$ NMR (151 MHz, $CDCl_3$) δ 168.3, 154.7, 141.6, 139.9, 135.4, 131.4, 130.2, 129.2, 126.7, 126.0, 125.7, 123.3, 121.6, 119.0, 80.0, 52.3, 34.6, 29.2, 28.7, 27.1, 26.9.

*tert-butyl 5-(3-acetamidophenyl)isoindoline-2-carboxylate **(m-4b)***

In a flame-dried microwave vial equipped with a magnetic stirring bar under inert atmosphere, *tert*-butyl 5-bromoisoindoline-2 carboxylate (100 mg, 0.34 mmol, 1 eq.), (3-acetamidophenyl)boronic acid (72 mg, 0.40

mmol, 1.2 eq.), $Cs_2CO_3$ (436 mg, 1.34 mmol, 4 eq.) and tetrakis(triphenylphosphine)palladium (11.6 mg, 0.01 mmol, 0.03 eq.) were added. Degassed 1,4-dioxane/$H_2O$/EtOH 1:1:1 was added and the reaction was left stirring for 48 hours at 80ºC. Upon completion, the reaction mixture was purified by automated silica gel flash column chromatography using a gradient elution (*n*-Heptane/EtOAc), yielding the product as a pale yellow oil (48 mg, 41%). $R_f$ (EtOAc) = 0.62. LC-MS (ESI) $R_t$ = 4.64 min, m/z calculated for $C_{21}H_{25}N_2O_3^+$ $[M + H]^+$ = 353.2 found 297.1 [M + H - *t*Bu]. $^1H$ NMR (600 MHz, MeOD) δ 7.83 (t, *J* = 2.0 Hz, 1H), 7.55 – 7.48 (m, 3H), 7.39 – 7.33 (m, 3H), 4.74 – 4.66 (m, 4H), 2.15 (s, 3H), 1.54 (s, 9H). $^{13}C$ NMR (151 MHz, MeOD) δ 171.8, 156.4, 142.9, 141.9, 140.4, 139.2, 138.6, 137.7, 137.1, 130.3, 127.64, 127.62, 124.15, 124.13, 123.8, 122.33, 122.31, 120.1, 119.8, 81.4, 53.4, 53.2, 53.0, 52.8, 28.8, 23.9.

*tert-butyl 5-(3-(2-methoxyethoxy)phenyl)isoindoline-2-carboxylate* ***(m-4c)***

In a flame-dried microwave vial equipped with a magnetic stirring bar under inert atmosphere, *tert*-butyl 5-bromoisoindoline-2 carboxylate (100 mg, 0.34 mmol, 1 eq.), (3-(2-methoxyethoxy)phenyl)boronic acid (3-(2-methoxyethoxy)phenyl)boronic acid (3-(2-methoxyethoxy)phenyl)boronic acid (78.4 mg, 0.40 mmol, 1.2 eq.), $Cs_2CO_3$ (436 mg, 1.34 mmol, 4 eq.) and tetrakis(triphenylphosphine)palladium (11.6 mg, 0.01 mmol, 0.03 eq.) were added. Degassed 1,4-dioxane/$H_2O$/EtOH 1:1:1 was added and the reaction was left stirring for 48 hours at 50ºC. Upon completion, the reaction mixture was purified by automated silica gel flash column chromatography using a gradient elution (*n*-Heptane/EtOAc), yielding the product as a pale yellow oil (20 mg, 16%). $R_f$ (*n*-Heptane/EtOAc 7:3) = 0.32. LC-MS (ESI) $R_t$ = 5.57 min, m/z calculated for $C_{22}H_{28}NO_4^+$ $[M + H]^+$ = 370.2 found 314.2 [M + H - *t*Bu]. $^1H$ NMR (400 MHz, MeOD) δ 7.51 – 7.46 (m, 2H), 7.34 – 7.28 (m, 2H), 7.18 – 7.14 (m, 1H), 7.13 (t, *J* = 2.1 Hz, 1H), 6.91 (ddd, *J* = 8.2, 2.6, 1.0 Hz, 1H), 4.70 – 4.59 (m, 4H), 4.20 – 4.11 (m, 2H), 3.80 – 3.70 (m, 2H), 3.43 (s, 3H), 1.53 (s, 9H). $^{13}C$ NMR (151 MHz, MeOD) δ 160.7, 156.4, 143.7, 141.99, 141.97, 139.1, 138.5, 137.5, 137.0, 130.9, 127.6, 124.06, 124.04, 122.32, 122.30, 120.7, 114.5, 114.4, 81.3, 72.3, 68.4, 59.3, 53.3, 53.1, 53.0, 52.8, 28.8.

*tert-butyl 5-(3-((2-methoxyethyl)carbamoyl)phenyl)isoindoline-2-carboxylate* ***(m-4d)***

In a flame-dried microwave vial equipped with a magnetic stirring bar under inert atmosphere, tert-butyl 5-bromoisoindoline-2 carboxylate (250 mg, 0.84 mmol, 1 eq.), (3-(2-methoxyethoxy)phenyl)boronic acid (197 mg, 1.01 mmol, 1.2 eq.), $Cs_2CO_3$ (1095 mg, 3.36 mmol, 4 eq.) and tetrakis(triphenylphosphine)palladium (34.7 mg, 0.03 mmol, 0.03 eq.) were added. Degassed 1,4-dioxane/$H_2O$/EtOH 1:1:1 was added and the reaction was left stirring for 48 hours at 50 °C. Upon completion, the reaction mixture was purified by automated C18 flash column chromatography using a gradient elution, yielding the product as a pale red oil (170 mg, 55%). LC-MS (ESI) $R_t$ = 4.48 min, m/z calculated for $C_{23}H_{29}N_2O_4^+$ $[M + H]^+$ = 397.2 found 397.2. $^1$H NMR (600 MHz, $CDCl_3$) δ 7.99 (s, 1H), 7.74 – 7.69 (m, 2H), 7.51 (dd, J = 7.5, 3.4 Hz, 2H), 7.48 (d, J = 18.0 Hz, 1H), 7.38 – 7.28 (m, 1H), 6.63 (s, 1H), 4.75 (d, J = 11.2 Hz, 2H), 4.71 (d, J = 6.5 Hz, 2H), 3.72 – 3.66 (m, 2H), 3.59 (t, J = 5.0 Hz, 2H), 3.40 (s, 3H), 1.54 (s, 9H). $^{13}$C NMR (151 MHz, $CDCl_3$) δ 167.7, 154.7, 141.5, 141.5, 139.8, 139.8, 138.2, 137.8, 136.9, 136.6, 135.0, 130.2, 129.1, 126.6, 126.6, 126.0, 126.0, 125.7, 125.6, 123.3, 123.0, 121.6, 121.4, 80.1, 71.1, 58.9, 52.3, 52.2, 52.0, 51.8, 39.9, 28.6.

*tert-butyl 5-(4-(methylcarbamoyl)phenyl)isoindoline-2-carboxylate* ***(p-4a)***

In a flame-dried microwave vial equipped with a magnetic stirring bar under inert atmosphere, *tert*-butyl 5-bromoisoindoline-2 carboxylate (100 mg, 0.34 mmol, 1 eq.), (4-(methylcarbamoyl)phenyl)boronic acid (71.5 mg, 0.40 mmol, 1.2 eq.), $Cs_2CO_3$ (436 mg, 1.34 mmol, 4 eq.) and tetrakis(triphenylphosphine)palladium (11.6 mg, 0.01 mmol, 0.03 eq.) were added. Degassed DME/$H_2O$ 1:1 was added and the reaction was left stirring overnight at 80ºC. Upon completion, the reaction mixture was diluted with EtOAc and aqueous HCl 1M was added dropwise until pH 4-5. The two phases are transferred in a separatory funnel and the aqueous phase was extracted with EtOAc. The combined organic layers were dried over anhydrous $Na_2SO_4$ and reduced *in vacuo*. Purification of the residue by automated C18 flash column chromatography using a gradient elution yielded the product as a yellow oil (43 mg, 36%). LC-MS (ESI) $R_t$ = 4.47 min, m/z calculated for $C_{21}H_{25}N_2O_3^+$ $[M + H]^+$ =353.2 found 353.2. $^1$H NMR (600 MHz, $CDCl_3$) δ 7.85 – 7.81 (m, 2H), 7.65 – 7.61 (m, 2H), 7.51 (d, *J* = 7.1 Hz, 2H), 7.33 (dd, *J* = 29.8, 8.0 Hz, 1H), 6.17 (s, 2H), 4.74 (d, *J* = 9.9 Hz, 2H), 4.70 (d, *J* = 7.7 Hz, 2H), 3.05 (d, *J* = 4.7 Hz, 4H), 1.53 (s, 9H). $^{13}$C NMR (151 MHz, $CDCl_3$) δ 168.3, 154.7, 141.6, 139.9, 135.4, 131.4, 130.2, 129.2, 126.7, 126.0, 125.7, 123.3, 121.6, 119.0, 80.0, 52.3, 34.6, 29.2, 28.7, 27.1, 26.9.

*N-(4-(isoindolin-5-yl)phenyl)acetamide TFA* ***(p-4b)***

In a flask equipped with magnetic stirrer compound ***p*-4b** (100 mg, 0.28 mmol, 1 eq.) was dissolved in 1 mL of methanol and 1 mL of a 4M HCl solution in dioxane was added dropwise. The reaction was stirred for for 24 hours at room temperature. Upon completion, the reaction mixture was concentrated *in vacuo*, and the crude was purified by reverse phase chromatography, providing the compound as a white amorphous solid. affording the title compound as a white amorphous solid (51 mg, 71% yield). LC-MS (ESI) $R_t$ = 2.60 min, m/z calculated for $C_{16}H_{17}N_2O^+$ $[M + H]^+$ = 253.1 found 253.1. $^1$H NMR (600 MHz, MeOD) δ 7.67 – 7.61 (m, 4H), 7.58 (d, *J* = 8.7

Hz, 2H), 7.46 (d, $J$ = 7.9 Hz, 1H), 4.67 (s, 2H), 4.65 (s, 2H), 2.15 (s, 3H). $^{13}$C NMR (151 MHz, MeOD) δ 171.8, 143.0, 139.7, 137.0, 136.5, 134.3, 128.5, 128.4, 124.3, 122.1, 121.5, 51.9, 51.8, 23.8. $^{19}$F NMR (376 MHz, Methanol-$d_4$) δ -77.1.

*tert-butyl 5-(4-(2-methoxyethoxy)phenyl)isoindoline-2-carboxylate* ***(p-4c)***

In a flame-dried microwave vial equipped with a magnetic stirring bar under inert atmosphere, *tert*-butyl 5-bromoisoindoline-2 carboxylate (100 mg, 0.34 mmol, 1 eq.), (4-(2-methoxyethoxy)phenyl)boronic acid (78.4 mg, 0.40 mmol, 1.2 eq.), $Cs_2CO_3$ (436 mg, 1.34 mmol, 4 eq.) and tetrakis(triphenylphosphine)palladium (11.6 mg, 0.01 mmol, 0.03 eq.) were added. Degassed 1,4-dioxane/$H_2O$/EtOH 1:1:1 was added and the reaction was left stirring for 48 hours at 50ºC. Upon completion, the reaction mixture was purified by automated siliga gel flash column chromatography using a gradient elution (*n*-Heptane/EtOAc), followed by automated C18 flash column chromatography using a gradient elution, yielded the product as a yellow oil (107 mg, 87%). LC-MS (ESI) $R_t$ = 5.26 min, m/z calculated for $C_{22}H_{28}NO_4^+$ $[M + H]^+$ = 370.2 found 314.2 $[M + H - t\text{–}Bu]^+$. $^1$H NMR (600 MHz, MeOD) δ 7.60 – 7.54 (m, 2H), 7.53 (dd, $J$ = 4.2, 1.6 Hz, 1H), 7.52 – 7.50 (m, 1H), 7.36 (dd, $J$ = 15.1, 7.8 Hz, 1H), 7.06 – 7.02 (m, 2H), 4.75 – 4.66 (m, 4H), 4.20 – 4.16 (m, 2H), 3.82 – 3.78 (m, 2H), 3.47 (s, 3H), 1.57 (s, 9H). $^{13}$C NMR (151 MHz, MeOD) δ 158.5, 155.1, 140.5, 140.4, 137.6, 137.0, 135.3, 134.7, 133.5, 127.7, 125.8, 122.6, 122.6, 120.4, 120.4, 114.6, 80.0, 70.8, 67.1, 57.8, 52.0, 51.7, 51.6, 51.4, 27.4.

*3-(isoindolin-5-yl)-N-methylbenzamide TFA* ***(m-5a)***

In a flask equipped with magnetic stirrer containing a 0°C solution of ***m*-4a** (190 mg, 0.54 mmol, 1 eq.) in 10 ml of $CHCl_3$/MeOH 1:1, 8.5 ml of TFA was added dropwise. The reaction was stirred for 5 minutes at 0°C, 24 hours at 40ºC, 2 hours at 50ºC and 2 hours at 60ºC. Upon completion, the reaction mixture was concentrated *in vacuo*. The residue was dissolved in water/ diethyl ether and aqueous HCl 1 M was added dropwise at 0°C until pH 3-4. The aqueous phase was extracted with diethyl ether , then basified with addition of aqueous NaOH 5 M until pH 10-11. The aqueous phase was extracted with EtOAc and $CHCl_3$. The combined EtOAc and $CHCl_3$ layers were dried over $Na_2SO_4$, filtered and concentrated *in vacuo* to afford the title compound as a pale brown oil (172 mg, 87% yield). LC-MS (ESI) $R_t$ = 2.52 min, m/z calculated for $C_{16}H_{17}N_2O^+$ $[M + H]^+$ = 253.1 found 253.1. $^1$H NMR (600 MHz, MeOD) δ 8.07 (t, $J$ = 1.9 Hz, 1H), 7.81 – 7.76 (m, 2H), 7.64 (d, $J$ = 1.6 Hz, 1H), 7.61 (dd, $J$ = 7.9, 1.6 Hz, 1H), 7.55 – 7.51 (m, 1H), 7.44 (d, $J$ = 7.9 Hz, 1H), 4.44 (s, 2H), 4.42 (s, 2H), 2.94 (d, $J$ = 2.5 Hz, 3H).

$^{13}C$ NMR (151 MHz, $CDCl_3$) δ 168.3, 168.2, 154.7, 141.6, 139.9, 135.5, 131.4, 130.2, 129.2, 126.7, 126.0, 125.7, 123.3, 121.6, 119.0, 79.0, 77.4, 77.2, 76.0, 52.3, 34.6, 32.0, 29.2, 28.7, 27.1, 27.0, 14.4.

*N-(3-(isoindolin-5-yl)phenyl)acetamide HCl* ***(m-5b)***

In a flask equipped with magnetic stirrer compound ***m*-4b** (48 mg, 0.14 mmol, 1 eq.) was dissolved in 1 ml of HCl 4M in dioxane. The reaction was stirred for for 24 hours at room temperature. Upon completion, the reaction mixture was concentrated *in vacuo*, affording the title compound as a off-white amorphous solid (33 mg, 96% yield). LC-MS (ESI) $R_t$ = 2.65 min, m/z calculated for $C_{16}H_{17}N_2O^+$ $[M + H]^+$ = 253.1 found 253.1. $^1H$ NMR (600 MHz, MeOD) δ 7.87 (s, 1H), 7.63 (s, 1H), 7.60 (d, J = 7.7 Hz, 1H), 7.52 (d, J = 7.7 Hz, 1H), 7.46 (d, J = 7.7 Hz, 1H), 7.38 (t, J = 7.7 Hz, 1H), 7.34 (d, J = 7.7 Hz, 1H), 4.67 (s, 2H), 4.66 (s, 2H), 2.17 (s, 3H). $^{13}C$ NMR (151 MHz, Methanol-$d_4$) δ 172.1, 143.2, 142.1, 140.3, 136.5, 134.8, 130.4, 128.8, 124.4, 124.0, 122.5, 120.6, 119.9, 51.9, 51.8, 23.8.

*5-(3-(2-methoxyethoxy)phenyl)isoindoline* ***(m-5c)***

In a flask equipped with magnetic stirrer compound ***m*-4c** (35 mg, 0.09 mmol, 1 eq.) was dissolved in 1 ml of HCl 4M in dioxane. The reaction was stirred for for 24 hours at room temperature. Upon completion, the reaction mixture was concentrated *in vacuo*. The residue was diluted in diethyl ether and extracted with HCl 1 M. The aqueous phase was washed with diethyl ether, then basified with addition of aqueous NaOH 5 M until pH 10-11 and extracted 5 times with EtOAc. The organic phase was dried over $Na_2SO_4$, filtered and concentrated *in vacuo* to afford the title compound as a pale brown oil (23 mg, 90% yield). LC-MS (ESI) $R_t$ = 3.06 min, m/z calculated for $C_{17}H_{20}NO_2^+$ $[M + H]^+$ = 270.1 found 270.2. $^1H$ NMR (400 MHz, MeOD) δ 7.58 (s, 1H), 7.56 (d, J = 8.0 Hz, 1H), 7.41 (d, J = 8.0 Hz, 1H), 7.34 (t, J = 8.1 Hz, 1H), 7.18 (d, J = 8.1 Hz, 2H), 7.17 – 7.12 (m, 1H), 6.94 (dd, J = 8.1, 2.5 Hz, 1H), 4.48 (s, 2H), 4.46 (s, 2H), 4.21 – 4.14 (m, 2H), 3.81 – 3.72 (m, 2H), 3.43 (s, 3H). $^{13}C$ NMR (151 MHz, Methanol-$d_4$) δ 160.7, 143.4, 142.6, 139.2, 137.5, 131.0, 128.2, 124.1, 122.3, 120.7, 114.6, 114.5, 72.3, 68.5, 59.3, 52.3, 52.1.

*3-(isoindolin-5-yl)-N-(2-methoxyethyl)benzamide TFA* ***(m-5d)***

In a flask equipped with magnetic stirrer containing a 0°C solution of ***m*-4d** (174 mg, 0.44 mmol, 1 eq.) in 10 ml of $CHCl_3$/MeOH 1:1, 8.3 ml of TFA was added dropwise. The reaction was stirred for 5 minutes at 0°C, 2 hours at room temperature, overnight at 40ºC, 2 hours at 50ºC, 2 hours at 60ºC and overnight at 40ºC. Upon completion, the reaction mixture was concentrated *in vacuo*. Purification of the residue by automated C18 flash column chromatography using a gradient elution, yielded the product as a pale red oil (TFA salt) (136 mg, 75%). LC-MS (ESI) $R_t$ = 2.64 min, m/z calculated for $C_{18}H_{21}N_2O_2^+$ $[M + H]^+$ = 297.2 found 297.2. $^1$H NMR (600 MHz, MeOD) δ 8.10 (t, *J* = 1.7 Hz, 1H), 7.83 (ddd, *J* = 7.7, 1.8, 1.0 Hz, 1H), 7.81 (ddd, *J* = 7.7, 1.9, 1.0 Hz, 1H), 7.75 – 7.74 (m, 1H), 7.74 – 7.71 (m, 1H), 7.56 (t, *J* = 7.5 Hz, 1H), 7.53 (d, *J* = 8.0 Hz, 1H), 4.70 (s, 2H), 4.68 (s, 2H), 3.60 (s, 4H), 3.39 (s, 3H). $^{13}$C NMR (151 MHz, MeOD) δ 170.1, 142.7, 141.8, 136.7, 136.4, 135.1, 131.2, 130.3, 129.0, 127.5, 127.1, 124.6, 122.6, 72.0, 59.0, 52.0, 51.8, 40.8.

*4-(isoindolin-5-yl)-N-methylbenzamide TFA* ***(p-5a)***

In a flask equipped with magnetic stirrer containing a 0°C solution of ***p*-4a** (162 mg, 0.46 mmol, 1 eq.) in 10 ml of $CHCl_3$/MeOH 1:1, 8.5 ml of TFA was added dropwise. The reaction was stirred for 5 minutes at 0°C and 24 hours at room temperature. Additional portions of TFA (0.35 mL) were added every two hours, for a total of 1 mL over 6 hours, and then stirred for 6 h at 35ºC. The temperature was further increased to 40ºC and the reaction was left stirring overnight. Upon completion, the reaction mixture was dried *in vacuo* and the crude residue was purified by automated C18 flash column chromatograpy using a gradient elution, which yielded the product as a brown oil (92 mg, 55%). LC-MS (ESI) $R_t$ = 2.49 min, m/z calculated for $C_{16}H_{17}N_2O^+$ $[M + H]^+$ = 253.1 found 253.1. $^1$H NMR (400 MHz, MeOD) δ 7.91 (d, *J* = 8.4 Hz, 2H), 7.79 – 7.67 (m, 4H), 7.53 (d, *J* = 7.9 Hz, 1H), 4.70 (s, 2H), 4.68 (s, 2H), 2.95 (s, 3H). $^{13}$C NMR (151 MHz, MeOD) δ 170.2, 144.5, 142.5, 136.7, 135.4, 134.9, 129.1, 128.9, 128.2, 124.6, 122.7, 52.0, 51.8, 26.9.$^{13}$C NMR (151 MHz, MeOD) δ 170.2, 144.5, 142.5, 136.7, 135.4, 134.9, 129.1, 128.9, 128.2, 124.6, 122.7, 52.0, 51.8, 26.9.

*N-(4-(isoindolin-5-yl)phenyl)acetamide TFA* ***(p-5b)***

In a flask equipped with magnetic stirrer compound ***p*-4b** (100 mg, 0.28 mmol, 1 eq.) was dissolved in 1 mL of methanol and 1 mL of a 4M HCl solution in dioxane was added dropwise. The reaction was stirred for for 24 hours at room temperature. Upon completion, the reaction mixture was concentrated *in vacuo*, and the crude was purified by reverse phase chromatography, providing the compound as a white amorphous solid. affording the title compound as a white amorphous solid (51 mg, 71% yield). LC-MS (ESI) $R_t$ = 2.60 min, m/z calculated for $C_{16}H_{17}N_2O^+$ $[M + H]^+$ = 253.1 found 253.1. $^1$H NMR (600 MHz, MeOD) δ 7.67 – 7.61 (m, 4H), 7.58 (d, *J* = 8.7 Hz, 2H), 7.46 (d, *J* = 7.9 Hz, 1H), 4.67 (s, 2H), 4.65 (s, 2H), 2.15 (s, 3H). $^{13}$C NMR (151 MHz, MeOD) δ 171.8,

143.0, 139.7, 137.0, 136.5, 134.3, 128.5, 128.4, 124.3, 122.1, 121.5, 51.9, 51.8, 23.8. $^{19}$F NMR (376 MHz, Methanol-$d_4$) δ -77.1.

*5-(4-(2-methoxyethoxy)phenyl)isoindoline TFA (**p-5c**)*

In a flask equipped with magnetic stirrer containing a 0°C solution of ***p*-4c** (162 mg, 0.46 mmol, 1 eq.) in 3 ml of $CHCl_3$/MeOH 1:1, 2.5 ml of TFA was added dropwise. The reaction was stirred for 5 minutes at 0°C and 2 hours at room temperature. Upon completion, the reaction mixture was dried *in vacuo* and the residue was dissolved in water/diethyl ether. Aqueous HCl 1M was added dropwise at 0°C until pH 3-4 and the aqueous phase was extracted with diethyl ether, then basified with addition of aqueous NaOH 5M until pH 10-11. The aqueous phase was extracted with EtOAc (4 x15 ml) and $CH_2Cl_2$ (3 x15 ml). Combined $CH_2Cl_2$ and EtOAc layers were dried over $Na_2SO_4$, filtered and concentrated *in vacuo*. Purification of the residue by automated C18 flash column chromatography using a gradient elution, yielded the product as a yellow oil (53 mg, 30%). LC-MS (ESI) $R_t$ =3.05 min, m/z calculated for $C_{17}H_{20}NO_2^+$ $[M + H]^+$ = 270.1 found 270.2. $^1$H NMR (600 MHz, MeOD) δ 7.61 (s, 1H), 7.60 (d, $J$ = 8.0 Hz, 2H), 7.58 – 7.53 (m, 2H), 7.44 (d, $J$ = 8.0 Hz, 1H), 7.04 – 6.99 (m, 2H), 4.67 (s, 2H), 4.65 (s, 2H), 4.19 – 4.10 (m, 2H), 3.78 – 3.72 (m, 2H), 3.43 (s, 3H). $^{13}$C NMR (151 MHz, MeOD) δ 160.2, 143.1, 136.4, 134.0, 133.9, 129.1, 128.3, 124.3, 121.9, 118.9, 117.0, 116.0, 72.2, 68.5, 59.2, 51.9, 51.7.

*N-methyl-3-(2-(5-(methylsulfonyl)-2-(2,2,3,3,3-pentafluoropropoxy)benzoyl)isoindolin-5-yl)benzamide (**m-1a**)*

In a flame-dried microwave vial equipped with a magnetic stirring bar under inert atmosphere, **3** (38.3 mg, 0.11 mmol, 1.00 eq.) and HBTU (49.3 mg, 0.13 mmol, 1.1 eq.), anhydrous DMF (0.8 mL) and anhydrous N,N-Diisopropylethylamine (0.1 mL, 5 eq.) were added an the mixture was stirred for 20 minutes at room temperature. After such time, ***m*-5a** (51 mg, 0.14 mmol, 1.2 eq.) was added and the reaction was stirred at room temperature overnight. Upon completion, the reaction mixture was diluted with $H_2O$ (30 mL) and lyophilized. The residue was dissolved in $H_2O$/MeCN 1:1 solution. The solution was purified by preparative HPLC, using a linear gradient from 0 to 100% B (MeCN/$H_2O$/TFA, 90:10:0.1) in solvent A ($H_2O$/TFA, 100:0.1) over 20 min, to afford the title compound as a transparent film (4.5 mg, 7%). LC-MS (ESI) $R_t$=4.13 min, m/z calculated for $[M + H]^+$ = 583.1 found 583.1. Purity by analytical HPLC 97.8% ($R_t$ = 14.9 min). $^1$H NMR (600 MHz, MeOD) δ 8.12 (dd, $J$ = 8.8, 2.3 Hz, 1H), 8.09 (t, $J$ = 1.8 Hz, 0.5H), 8.03 (dt, $J$ = 7.3, 2.0 Hz, 1.5H), 7.84 – 7.79 (m, 1H), 7.80 – 7.74 (m, 1H), 7.71 (s, 0.5H), 7.65 (dd, $J$ = 7.8, 1.8 Hz, 0.5H), 7.61 (dd, $J$ = 7.9, 1.7 Hz, 0.5H), 7.58 – 7.52 (m, 1H), 7.51 (t, $J$ = 7.7 Hz, 1H), 7.48 (dd, $J$ = 8.9, 2.2 Hz, 1H), 7.31 (d, $J$ = 8.0 Hz, 0.5H), 4.99 (d, $J$ = 15.6 Hz, 2H), 4.91 (t, $J$ = 12.2

Hz, 2H), 4.69 (d, *J* = 13.4 Hz, 2H), 3.17 (d, *J* = 1.5 Hz, 3H), 2.94 (d, *J* = 12.2 Hz, 3H). $^{13}$C NMR (151 MHz, MeOD) δ 169.2, 166.3, 156.5, 156.5, 141.0, 141.0, 140.3, 140.2, 136.9, 136.4, 135.5, 135.1, 135.1, 135.1, 134.8, 130.8, 129.8, 129.7, 128.8, 128.8, 127.5, 127.5, 127.4, 127.4, 126.7, 126.6, 125.8, 125.7, 125.5, 123.2, 122.8, 121.3, 121.0, 113.2, 113.2, 65.1, 64.9, 64.7, 53.0, 52.8, 51.7, 51.6, 43.1, 25.5, 25.5.

*N-(3-(2-(5-(methylsulfonyl)-2-(2,2,3,3,3-pentafluoropropoxy)benzoyl)isoindolin-5-yl)phenyl)acetamide* ***(m-1b)***

In a flame-dried microwave vial equipped with a magnetic stirring bar under inert atmosphere, **3** (41 mg, 0.12 mmol, 1.0 eq.) and HBTU (50 mg, 0.13 mmol, 1.1 eq.), anhydrous DMF (0.8 mL) and anhydrous N,N-diisopropylethylamine (0.11 mL, 0.59 mmol, 5 eq.) were added an the mixture was stirred for 20 minutes at room temperature. After such time, ***m*-5b** (30 mg, 0.12 mmol, 1.0 eq.) was added and the reaction was stirred at room temperature overnight. Upon completion, the reaction mixture was diluted with $H_2O$ (20 mL) and lyophilized. The residue was dissolved in $H_2O$/MeCN 1:1 solution. The solution was purified by preparative HPLC, using an gradient from 10 of B to 100% B (MeCN/$H_2O$/TFA, 90:10:0.1) in solvent A ($H_2O$/TFA, 100:0.1), to afford the title compound as a colorless film (40 mg, 58%). LC-MS (ESI) Rt=4.13 min, m/z calculated for $C_{27}H_{24}F_5N_2O_5S^+$ $[M + H]^+$ = 583.1 found 583.1. Purity by analytical HPLC 100% ($R_t$ =15.11 min). $^1$H NMR (600 MHz, MeOD) δ 8.11 (dd, *J* = 8.8, 2.3 Hz, 1H), 8.02 (dd, *J* = 2.3, 0.6 Hz, 1H), 7.86 (t, *J* = 1.9 Hz, 0.5H), 7.82 (t, *J* = 1.9 Hz, 0.5H), 7.63 (d, *J* = 1.6 Hz, 0.5H), 7.58 (dd, *J* = 7.9, 1.6 Hz, 0.5H), 7.54 (dd, *J* = 7.9, 1.6 Hz, 0.5H), 7.51 (dt, *J* = 7.3, 1.9 Hz, 0.5H), 7.49 – 7.44 (m, 2H), 7.43 (d, *J* = 1.6 Hz, 0.5H), 7.40 – 7.34 (m, 1.5H), 7.33 – 7.30 (m, 0.5H), 7.27 (d, *J* = 7.9 Hz, 0.5H), 4.97 (d, *J* = 12.8 Hz, 2H), 4.91 (t, *J* = 12.2 Hz, 2H), 4.67 (d, *J* = 11.5 Hz, 2H), 3.16 (d, *J* = 1.2 Hz, 3H), 2.15 (s, 1.5H), 2.13 (s, 1.5H). $^{13}$C NMR (151 MHz, Methanol-*d*$_4$) δ 171.78, 171.76, 167.7, 157.92, 157.91, 142.8, 142.7, 142.4, 142.2, 140.5, 140.4, 138.1, 137.7, 136.6, 136.5, 136.2, 132.2, 130.34, 130.30, 128.87, 128.85, 128.1, 127.9, 124.4, 124.1, 123.9, 123.8, 122.6, 122.2, 120.20, 120.17, 119.98 (qt, *J* = 285.6, 34.7 Hz), 119.82, 119.79, 114.59, 114.57, 113.90 (tq, *J* = 254.2, 38.7 Hz), 66.30 (t, *J* = 29.1 Hz), 54.4, 54.2, 53.2, 53.0, 49.6, 44.5, 23.88, 23.86. $^{19}$F NMR (376 MHz, Methanol-*d*$_4$) δ -76.9, -77.7, -84.8, -84.8, -124.4, -124.4.

*(5-(3-(2-methoxyethoxy)phenyl)isoindolin-2-yl)(5-(methylsulfonyl)-2-(2,2,3,3,3-pentafluoropropoxy)phenyl)methanone* ***(m-1c)***

In a flame-dried microwave vial equipped with a magnetic stirring bar under inert atmosphere, **3** (20 mg, 0.056 mmol, 1.0 eq.) and HBTU (23 mg, 0.061 mmol, 1.1 eq.), anhydrous DMF (0.8 mL) and anhydrous N,N-diisopropylethylamine (0.05 mL, 0.28 mmol, 5 eq.) were added an the mixture was stirred for 20 minutes at room temperature. After such time, ***m*-5c** (15 mg, 0.056 mmol, 1.0 eq.) was added and the reaction was stirred at room temperature overnight. Upon completion, the reaction mixture was diluted with $H_2O$ (30 mL) and lyophilized. The residue was dissolved in $H_2O$/MeCN 1:1 solution. The solution was purified by preparative HPLC, using an isocratic method of 50% B (MeCN/$H_2O$/TFA, 90:10:0.1) in solvent A ($H_2O$/TFA, 100:0.1), to afford the title compound as a colorless film (10 mg, 30%). LC-MS (ESI) Rt=4.64 min, m/z calculated for $C_{28}H_{27}F_5NO_6S^+$ [M + H]$^+$ = 600.1 found 600.1. Purity by analytical HPLC 100% ($R_t$ =16.54 min). $^1$H NMR (600 MHz, MeOD) δ 8.11 (dd, *J* = 8.8, 2.4 Hz, 1H), 8.02 (dd, *J* = 2.4, 1.0 Hz, 1H), 7.64 (d, *J* = 1.6 Hz, 0.5H), 7.58 (dd, *J* = 8.0, 1.6 Hz, 0.5H), 7.54 (dd, *J* = 8.0, 1.6 Hz, 0.5H), 7.47 (dd, *J* = 8.8, 1.0 Hz, 1H), 7.45 (d, *J* = 8.0 Hz, 0.5H), 7.43 (d, *J* = 1.6 Hz, 0.5H), 7.38 – 7.29 (m, 1H), 7.26 (d, *J* = 8.0 Hz, 0.5H), 7.23 – 7.20 (m, 0.5H), 7.19 (t, *J* = 2.3 Hz, 0.5H), 7.18 – 7.15 (m, 0.5H), 7.14 (t, *J* = 2.3 Hz, 0.5H), 6.94 (ddd, *J* = 8.2, 2.3, 0.9 Hz, 0.5H), 6.92 (ddd, *J* = 8.2, 2.3, 0.9 Hz, 0.5H), 5.02 – 4.87 (m, 4H), 4.67 (d, *J* = 11.7 Hz, 2H), 4.23 – 4.13 (m, 2H), 3.81 – 3.70 (m, 2H), 3.44 (s, 1.5H), 3.42 (s, 1.5H), 3.16 (d, *J* = 1.6 Hz, 3H). $^{13}$C NMR (151 MHz, Methanol-*$d_4$*) δ 167.7, 160.8, 160.7, 157.93, 157.91, 143.59, 143.55, 142.5, 142.4, 138.1, 137.6, 136.50, 136.47, 136.46, 136.1, 132.2, 131.0, 130.9, 128.9, 128.8, 128.1, 127.9, 124.3, 124.0, 122.6, 122.3, 120.80, 120.76, 120.0 (qt, *J* = 285.6, 34.9 Hz), 114.7, 114.59, 114.57, 114.52, 114.46, 113.8 (tq, *J* = 254.6, 37.8 Hz), 72.29, 72.27, 66.30 (td, *J* = 29.2, 2.6 Hz), 59.3, 59.2, 54.4, 54.2, 53.2, 52.9, 49.6, 44.5. $^{19}$F NMR (376 MHz, Methanol-*$d_4$*) δ -76.9, -77.5, -84.8, -124.4, -124.5.

*N-(2-methoxyethyl)-3-(2-(5-(methylsulfonyl)-2-(2,2,3,3,3-pentafluoropropoxy)benzoyl)isoindolin-5-yl)benzamide **(m-1d)***

In a flame-dried microwave vial equipped with a magnetic stirring bar under inert atmosphere, **3** (90.4 mg, 0.26 mmol, 1.0 eq.) and HBTU (110 mg, 0.29 mmol, 1.1 eq.), anhydrous DMF (1.6 mL) and anhydrous N,N-Diisopropylethylamine (0.2 mL, 5 eq.) were added an the mixture was stirred for 20 minutes at room temperature. After such time, ***m*-5d** (106.4 mg, 0.26 mmol, 1 eq.) was added and the reaction was stirred at room temperature overnight. Upon completion, the reaction mixture was diluted with $H_2O$ (30 mL) and lyophilized. The residue was dissolved in $H_2O$/MeCN 1:1 solution. The solution was purified by preparative HPLC, using an isocrat of 40% B (MeCN/$H_2O$/TFA, 90:10:0.1) in solvent A ($H_2O$/TFA, 100:0.1) over 20 min, to afford the title compound as a transparent film (11.4 mg, 7%). LC-MS $R_t$=4.01min, m/z calculated for [M + H]$^+$ = 627.1 found 627.1. Purity by analytical HPLC 98.8% ($R_t$ =15.09 min). $^1$H NMR (600 MHz, $CDCl_3$) δ 8.89 (d, *J* = 5.1 Hz, 1H), 8.49 (t, *J* = 7.4 Hz, 0.5H), 8.04 – 7.99 (m, 2H), 7.98 – 7.96 (m, 1H), 7.74 – 7.71 (m, 0.5H), 7.71 – 7.67 (m, 0.5H), 7.63 (d, *J* = 7.8 Hz, 0.5H), 7.56 – 7.53 (m, 1), 7.51 – 7.44 (m, 1.5H), 7.40 (d, *J* = 8.0 Hz, 0.5H), 7.36 (s, 0.5H), 7.20 (d, *J* = 7.9 Hz, 0.5H), 7.18 (dd, *J* = 8.7, 1.8 Hz, 1H), 6.72 (t, *J* = 5.6 Hz, 1H), 4.98 (d, *J* = 11.6 Hz, 2H), 4.60 (m, 4H), 3.70 – 3.61

(m, 2H), 3.60 – 3.49 (m, 2H), 3.36 (d, $J$ = 12.7 Hz, 3H), 3.06 (s, 3H). $^{13}$C NMR (151 MHz, $CDCl_3$) δ 167.4, 167.4, 165.1, 156.7, 146.3, 141.2, 141.1, 140.5, 140.3, 136.7, 135.5, 135.5, 135.4, 135.3, 135.3, 135.3, 130.8, 130.8, 130.2, 130.1, 129.2, 128.8, 128.8, 128.2, 127.4, 127.3, 127.1, 126.1, 126.1, 125.9, 125.9, 123.6, 123.1, 121.9, 121.5, 121.4, 121.2, 119.6, 119.3, 119.1, 117.7, 117.4, 117.2, 115.8, 115.6, 115.3, 114.1, 113.9, 113.6, 113.4, 113.3, 112.4, 112.2, 111.9, 111.7, 110.7, 110.5, 110.2, 110.0, 71.2, 71.2, 65.9, 65.7, 65.5, 58.9, 58.9, 53.2, 53.1, 52.2, 52.0, 44.8, 44.8, 39.9, 39.8.

*N-methyl-4-(2-(5-(methylsulfonyl)-2-(2,2,3,3,3-pentafluoropropoxy)benzoyl)isoindolin-5-yl)benzamide* ***(p-1a)***

In a flame-dried microwave vial equipped with a magnetic stirring bar under inert atmosphere, **3** (38.3 mg, 0.11 mmol, 1.00 eq.) and HBTU (49.3 mg, 0.13 mmol, 1.1 eq.), anhydrous DMF (0.8 mL) and anhydrous N,N-Diisopropylethylamine (0.1 mL, 5 eq.) were added an the mixture was stirred for 20 minutes at room temperature. After such time, ***p*-5a** (51.1 mg, 0.14 mmol, 1.2 eq.) was added and the reaction was stirred at room temperature overnight. Upon completion, the reaction mixture was diluted with $H_2O$ (30 mL) and lyophilized. The residue was dissolved in $H_2O$/MeCN 1:1 solution. The solution was purified by preparative HPLC, using an isocrat 40% B (MeCN/$H_2O$/TFA, 90:10:0.1) in solvent A ($H_2O$/TFA, 100:0.1) over 20 min, to afford the title compound as a transparent film (37.9 mg, 59%). LC-MS (ESI) $R_t$=3.87 min, m/z calculated for $[M + H]^+$ = 583.1 found 583.2. Purity by analytical HPLC 100% ($R_t$ = 15.0 min). $^1$H NMR (600 MHz, MeOD) δ 8.11 (dd, $J$ = 8.8, 2.3 Hz, 1H), 8.03 (t, $J$ = 2.0 Hz, 1H), 7.92 – 7.85 (m, 2H), 7.76 – 7.68 (m, 2.5H), 7.65 (dd, $J$ = 8.0, 1.7 Hz, 0.5H), 7.60 (dd, $J$ = 8.0, 1.7 Hz, 0.5H), 7.52 – 7.49 (m, 1H), 7.47 (dd, $J$ = 8.9, 1.8 Hz, 1H), 7.31 (d, $J$ = 7.9 Hz, 0.5H), 4.98 (d, $J$ = 14.6 Hz, 2H), 4.91 (t, $J$ = 12.2 Hz, 2H), 4.68 (d, $J$ = 12.7 Hz, 2H), 3.17 (d, $J$ = 1.4 Hz, 3H), 2.95 (s, 1.5H), 2.93 (s, 1.5H). $^{13}$C NMR (151 MHz, MeOD) δ 170.38, 170.35, 167.7, 157.93, 157.91, 145.14, 145.09, 141.4, 141.3, 138.3, 137.9, 137.2, 136.8, 136.48, 136.47, 134.48, 134.45, 132.2, 128.88, 128.86, 128.84, 128.81, 128.17, 128.16, 128.1, 128.0, 124.6, 124.3, 122.7, 122.4, 120.0 (qt, $J$ = 285.6, 34.7 Hz), 114.60, 114.58, 113.90 (tq, $J$ = 254.5, 38.2 Hz), 66.30 (td, $J$ = 29.3, 2.8 Hz), 54.4, 54.2, 53.1, 53.0, 44.5, 26.93, 26.92. $^{19}$F NMR (376 MHz, MeOD) δ -84.8, -124.5.

*N-(4-(2-(5-(methylsulfonyl)-2-(2,2,3,3,3-pentafluoropropoxy)benzoyl)isoindolin-5-yl)phenyl)acetamide* ***(p-1b)***

In a flame-dried microwave vial equipped with a magnetic stirring bar under inert atmosphere, **3** (38 mg, 0.11 mmol, 1.0 eq.) and HBTU (46 mg, 0.12 mmol, 1.1 eq.), anhydrous DMF (0.6 mL) and anhydrous N,N-diisopropylethylamine (0.10 mL, 0.55 mmol, 5 eq.) were added an the mixture was stirred for 20 minutes at room temperature. After such time, ***p*-5b** (40 mg, 0.11 mmol, 1.0 eq.) was added and the reaction was stirred at room temperature overnight. Upon completion, the reaction mixture was diluted with $H_2O$ (25 mL) and lyophilized. The residue was dissolved in $H_2O$/MeCN 1:1 solution. The solution was purified by preparative HPLC, using an isocratic method of 40% B (MeCN/$H_2O$/TFA, 90:10:0.1) in solvent A ($H_2O$/TFA, 100:0.1), to afford the title compound as a colorless film (8 mg, 13%). LC-MS (ESI) Rt=3.96 min, m/z calculated for $C_{27}H_{24}F_5N_2O_5S^+$ [M + H]$^+$ = 583.1 found 583.1. Purity by analytical HPLC 98% ($R_t$ =15.23 min). $^1$H NMR (600 MHz, MeOD) δ 8.11 (dd, *J* = 8.9, 2.4 Hz, 1H), 8.02 (dd, *J* = 2.4, 1.4 Hz, 1H), 7.66 – 7.56 (m, 4H), 7.56 – 7.52 (m, 1.5H), 7.47 (dd, *J* = 8.9, 1.7 Hz, 1H), 7.45 – 7.43 (m, 1H), 7.26 (d, *J* = 7.9 Hz, 0.5H), 4.96 (d, *J* = 15.0 Hz, 2H), 4.91 (t, *J* = 12.2 Hz, 2H), 4.66 (d, *J* = 13.0 Hz, 2H), 3.18 – 3.15 (m, 3H), 2.15 (s, 1.5H), 2.13 (s, 1.5H). $^{13}$C NMR (151 MHz, Methanol-*d*$_4$) δ 171.68, 171.66, 167.7, 157.93, 157.91, 142.1, 141.9, 139.48, 139.45, 138.1, 137.69, 137.65, 137.6, 136.47, 136.45, 136.1, 135.7, 132.2, 128.87, 128.86, 128.8, 128.4, 128.3, 127.7, 127.6, 124.4, 124.1, 122.2, 121.9, 121.5, 121.4, 120.00 (dt, *J* = 285.3, 34.4 Hz), 114.59, 114.57, 114.0 (tq, *J* = 254.6, 38.4 Hz), 66.30 (td, *J* = 29.2, 2.5 Hz), 54.5, 54.2, 53.2, 52.9, 44.53, 44.49, 23.84, 23.82, 19.7, 9.2. $^{19}$F NMR (376 MHz, Methanol-*d*$_4$) δ -84.82, -84.83, -124.44, -124.45.

*(5-(4-(2-methoxyethoxy)phenyl)isoindolin-2-yl)(5-(methylsulfonyl)-2-(2,2,3,3,3-pentafluoropropoxy)phenyl)methanone (**p-1c**)*

In a flame-dried microwave vial equipped with a magnetic stirring bar under inert atmosphere, **3** (45.2 mg, 0.13 mmol, 1.00 eq.) and HBTU (53.1 mg, 0.14 mmol, 1.1 eq.), anhydrous DMF (0.8 mL) and anhydrous N,N-Diisopropylethylamine (0.1 mL, 5 eq.) were added an the mixture was stirred for 20 minutes at room temperature. After such time, ***p*-5c** (53.5 mg, 0.14 mmol, 1.1 eq.) was added and the reaction was stirred at room temperature overnight. Upon completion, the reaction mixture was diluted with $H_2O$ (30 mL) and lyophilized. The residue was dissolved in $H_2O$/MeCN 1:1 solution. The solution was purified by preparative HPLC, using an isocrat 45% B (MeCN/$H_2O$/TFA, 90:10:0.1) in solvent A ($H_2O$/TFA, 100:0.1) over 20 min, to afford the title compound as a transparent film (7.8 mg, 10%). LC-MS (ESI) $R_t$=4.49 min, m/z calculated for [M + H]$^+$ = 600.1 found 600.1. Purity by analytical HPLC 99.52% ($R_t$ =16.5 min). $^1$H NMR (600 MHz, MeOD) δ 8.11 (dd, *J* = 8.8, 2.4 Hz, 1H), 8.02 (s, 1H), 7.23 (d, *J* = 7.9 Hz, 0.5H), 7.01 (dd, *J* = 16.7, 8.7 Hz, 2H), 4.95 (d, *J* = 14.8 Hz, 2H), 4.91 (t, *J* = 12.2 Hz, 2H), 4.65 (d, *J* = 13.2 Hz, 2H), 4.18 – 4.12 (m, 2H), 3.78 – 3.73 (m, 2H), 3.43 (d, *J* = 7.7 Hz, 3H), 3.16 (s, 3H). $^{13}$C NMR (151 MHz, MeOD) δ 166.3, 158.6, 158.6, 156.5, 156.5, 140.9, 140.7, 136.6, 136.1, 135.1, 135.0, 134.5, 134.2, 133.8, 133.4, 133.3, 131.9, 131.5, 130.7, 127.8, 127.7, 127.5, 127.5, 127.4, 127.4, 126.2, 126.0, 122.9, 122.6, 120.6, 120.3, 117.6, 114.6, 114.6, 114.3, 114.1, 113.2, 113.2, 113.0, 112.6, 112.4, 70.8, 70.8,

67.1, 67.1, δ 65.46 – 64.43 (m), 63.0, 57.9, 57.8, 53.1, 52.8, 51.8, 51.5, 43.1, 7.8.$^{19}$F NMR (376 MHz, MeOD) δ -84.8, -124.4.

*5-bromoisoindoline (**6**)*

In a flask containing tert-butyl 5 bromoisoindoline-2-carboxylate (3.3 g, 11.1 mmol, 1 eq.), trifluoroacetic acid (50 mL) was added. The reaction mixture was stirred for 2.5 hours at 40ºC. Upon completion, it was diluted in water and washed with EtOAc (x4). The aqueous phase was basified by dropwise addition of aqueous 2M NaOH solution and extracted with DCM (x5). The organic phase was dried over anhydrous $Na_2SO_4$, filtered and the organic solvent was removed *in vacuo*. The residue was purified by automated C18 flash column chromatography using a gradient elution. The title compound was obtained as a white solid (TFA salt) (1.93 g, 56%). LC-MS (ESI) $R_t$=2.43 min, m/z calculated for $C_8H_9BrN^+$ $[M + H]^+$ = 197.99 and 199.99, found 198.0 and 200.0. $^1$H NMR (600 MHz, MeOD) δ 7.65 (s, 1H), 7.58 (d, *J* = 8.2 Hz, 1H), 7.37 (d, *J* = 8.2 Hz, 1H), 4.64 (s, 2H), 4.60 (s, 2H). $^{13}$C NMR (151 MHz, MeOD) δ 166.4, 163.1, 161.2, 155.5, 153.9, 151.7, 79.8, 79.7.

*5-bromoisoindolin-2-yl)(5-(methylsulfonyl)-2-(2,2,3,3,3-pentafluoropropoxy)phenyl) methanone (7)*

In a flame-dried flask equipped with a magnetic stirring bar under inert atmosphere, **3** (1785 mg, 5.1 mmol, 1.0 eq.) and HBTU (2139mg, 5.6 mmol, 1.1 eq.), anhydrous DMF (17 mL) and anhydrous N,N-Diisopropylethylamine (0.2 mL, 5 eq.) were added and the mixture was stirred for 20 minutes at room temperature. After such time, **6** (1600 mg, 5.1 mmol, 1 eq.) was added and the reaction was stirred at room temperature overnight. Upon completion, the reaction mixture was diluted with EtOAc and the resulting solution was washed with saturated aqueous ammonium chloride solution. The organic phase was dried over anhydrous $Na_2SO_4$, filtered and the organic solvent was evaporated under reduced pressure. The residue was purified by automated silica gel flash column chromatography using a gradient elution (*n*-Heptane/EtOAc). The title compound was obtained as a white solid (2.48 g, 92%). LC-MS (ESI) $R_t$=4.48 min, m/z calculated for $C_{19}H_{16}BrF_5NO_4S^+$ $[M + H]^+$ = 529.99 and 527.99, found 528.0 and 530.0. $^1$H NMR (400 MHz, $CDCl_3$) δ 8.04 (dt, *J* = 8.7, 1.9 Hz, 1H), 7.98 (d, *J* = 2.3 Hz, 1H), 7.50 (s, 0.5H), 7.46 (d, *J* = 8.1 Hz, 0.5H), 7.41 (d, *J* = 8.1 Hz, 0.5H), 7.28 (s, 0.5H), 7.22 (d, *J* = 8.2 Hz, 0.5H), 7.14 (dd, *J* = 8.8, 1.9 Hz, 1H), 7.01 (d, *J* = 8.2 Hz, 0.5H), 4.93 (d, *J* = 16.7 Hz, 2H), 4.61 – 4.48 (m, 4H), 3.07 (s, 3H). $^{13}$C NMR (101 MHz, $CDCl_3$) δ 165.4, 156.6, 137.7, 137.6, 135.7, 135.7, 134.5, 134.4, 131.4, 131.1, 131.1, 128.3, 128.3, 126.4, 125.8, 124.7, 124.1, 122.0, 121.7, 113.2, 65.67 (t, J = 28.9 Hz), 52.9, 52.8, 51.9, 44.6. $^{19}$F NMR (376 MHz, $CDCl_3$) δ -83.4, -122.8.

*N-(2-methoxyethyl)-4-(2-(5-(methylsulfonyl)-2-(2,2,3,3,3 pentafluoropropoxy)benzoyl)isoindolin-5-yl)benzamide* ***(p-1d)***

In a flame-dried microwave vial equipped with a magnetic stirring bar under inert atmosphere **7** (37 mg, 0.07 mmol, 1 eq.), (4-((2-methoxyethyl)carbamoyl)phenyl)boronic acid (19 mg, 0.08 mmol, 1.1 eq.), $Cs_2CO_3$ (90 mg, 0.28 mmol, 4 eq.) and tetrakis(triphenylphosphine)palladium(0) (2.4 mg, 0.002 mmol, 0.03 eq.) were added. A degassed mixture of 1,4-dioxane/$H_2O$/EtOH 1:1:1 was added and the reaction mixture was stirred overnight at 50ºC. Upon completion, the solvent was removed *in vacuo* and the residue was purified by automated silica gel flash column chromatography using a gradient elution (DCM/Acetone). The title compound was obtained as a white solid (37.2 mg, 85%). LC-MS (ESI) $R_t$=3.95 min, m/z calculated for $C_{29}H_{28}F_5N_2O_6S^+$ $[M + H]^+$ = 627.16 , found 627.2. $R_t$ (HPLC) = 14.9 min. $^1H$ NMR (600 MHz, $CDCl_3$) δ 8.05 (dd, *J* = 8.7, 2.3 Hz, 1H), 8.01 (d, *J* = 2.3 Hz, 1H), 7.87 (d, *J* = 8.3 Hz, 1H), 7.85 (d, *J* = 8.4 Hz, 1H), 7.65 (d, *J* = 8.3 Hz, 1H), 7.60 (d, *J* = 8.3 Hz, 1H), 7.57 (s, 0.5H), 7.56 (s, 0.5H), 7.52 (d, *J* = 7.9 Hz, 0.5H), 7.44 (d, *J* = 7.9 Hz, 89 0.5H), 7.36 (s, 0.5H), 7.23 (d, *J* = 7.9 Hz, 0.5H), 7.15 (d, *J* = 8.7 Hz, 1H), 6.55 (dt, *J* = 12.2, 5.6 Hz, 1H), 5.03 (d, *J* = 9.9 Hz, 2H), 4.62 (d, *J* = 10.5 Hz, 2H), 4.58 (d, *J* = 11.8 Hz, 2H), 3.68 (dq, *J* = 10.5, 5.2 Hz, 2H), 3.58 (dt, *J* = 8.6, 5.0 Hz, 2H), 3.40 (d, *J* = 7.9 Hz, 3H), 3.08 (d, *J* = 1.5 Hz, 3H). $^{13}C$ NMR (151 MHz, $CDCl_3$) δ 167.2, 167.1, 165.0, 156.8, 156.7, 143.7, 143.7, 140.4, 140.2, 136.8, 136.7, 135.7, 135.7, 135.6, 133.6, 133.6, 130.8, 128.9, 128.3, 127.7, 127.4, 127.3, 127.3, 127.1, 123.7, 123.1, 121.9, 121.5, 121.3, 121.1, 119.6, 119.4, 119.2, 117.7, 117.5, 117.3, 115.8, 115.6, 115.4, 114.2, 113.9, 113.7, 113.4, 112.5, 112.2, 112.0, 111.7, 110.8, 110.5, 110.3, 110.0, 77.4, 77.2, 76.9, 71.3, 71.3, 65.9, 65.7, 65.5, 58.9, 58.9, 53.9, 53.2, 53.1, 52.2, 52.0, 44.8, 39.8, 39.8. $^{19}F$ NMR (376 MHz, $CDCl_3$) δ -83.3, -122.8.

*3-(2-(5-(methylsulfonyl)-2-(2,2,3,3,3-pentafluoropropoxy)benzoyl)isoindolin-5-yl)benzoic acid* ***(8)***

In a flame-dried microwave vial equipped with a magnetic stirring bar under inert atmosphere **7** (600 mg, 1.14 mmol, 1 eq.), 3-carboxyphenylboronic acid (225 mg, 1.36 mmol, 1.2 eq.), $Cs_2CO_3$ (1480 mg, 4.56 mmol, 4 eq.) and tetrakis(triphenylphosphine)palladium(0) (39.4 mg, 0.03 mmol, 0.03 eq.) were added. A degassed mixture of 1,4-dioxane/$H_2O$/EtOH 1:1:1 was added and the reaction mixture was stirred overnight at 50ºC. Upon completion, the solvent was removed *in vacuo* and the residue was purified by two rounds of automated silica gel flash column chromatography using gradient elution (DCM/Acetone followed by DCM/MeOH). The title compound was obtained as a white solid (512 mg, 79%). LC-MS (ESI) Rt=4.07 min, m/z calculated for $C_{26}H_{21}F_5NO_6S^+$ $[M + H]^+$ = 570.10, found 570.1. $^1H$ NMR (600 MHz, $CDCl_3$) δ 8.34 (t, *J* = 1.8 Hz, 0.5H), 8.28 (t, *J* = 1.8 Hz, 0.5H), 8.12

– 8.04 (m, 2H), 8.02 (d, *J* = 2.3 Hz, 1H), 7.84 (dt, *J* = 7.9, 1.4 Hz, 0.5H), 7.78 (dt, *J* = 7.9, 1.4 Hz, 0.5H), 7.62 – 7.52 (m, 2.5H), 7.45 (d, *J* = 8.0 Hz, 0.5H), 7.39 (s, 0.5H), 7.24 (d, *J* = 7.9 Hz, 0.5H), 7.16 (dd, *J* = 8.8, 2.7 Hz, 1H), 5.05 (d, *J* = 11.4 Hz, 2H), 4.64 (d, *J* = 11.0 Hz, 2H), 4.60 (t, *J* = 11.8 Hz, 2H), 3.09 (s, 3H). $^{13}$C NMR (101 MHz, $CDCl_3$) δ 165.4, 156.6, 137.7, 137.6, 135.7, 135.7, 134.5, 134.4, 131.4, 131.1, 131.1, 128.3, 128.3, 126.4, 125.8, 124.7, 124.1, 122.0, 121.7, 113.2, 66.0, 65.7, 65.4, 52.9, 52.8, 51.9, 44.6. $^{19}$F NMR (376 MHz, $CDCl_3$) δ -83.4, -122.8.

*3-(2-(5-(methylsulfonyl)-2-(2,2,3,3,3 pentafluoropropoxy)benzoyl)isoindolin-5-yl)-N-(prop-2-yn-1-yl)benzamide* ***(m-9e)***

In a flame-dried microwave vial and under inert conditions, a solution of 3-(2-(5-(methylsulfonyl)-2-(2,2,3,3,3-pentafluoropropoxy)benzoyl)isoindolin-5-yl)benzoic acid **8** (20 mg, 0.035 mmol, 1 eq.), HBTU (15 mg, 0.039 mmol, 1.1 eq.) and *N*-ethyl-diisopropylamine (23 mg, 0.18 mmol, 5 eq.) were dissolved in anhydrous DMF (0.3 mL) and stirred at room temperature for 20 minutes. A solution of propargylamine (1.9 mg, 0.035 mmol, 1 eq.) in DMF (0.3 mL) was added dropwise and the solution was stirred at room temperature for 18 hours. After completion, the reaction mixture was diluted with DCM (2 mL) and washed three times with an aqueous solution of HCl 1 M (2 mL) and then with an aqueous solution of $NaHCO_3$. The organic phase was dried over anhydrous $Na_2SO_4$, filtered, and evaporated under vacuum. The crude that was purified by RP flash column chromatography, providing the desired compound in 80% yield (17 mg, 0.028 mmol). $^1$H NMR (400 MHz, $CD_3CN$) δ 8.07 – 8.02 (m, 2H), 7.91 (d, *J* = 2.4 Hz, 1H), 7.84 – 7.74 (m, 2H), 7.70 (s, 0.5H), 7.67 – 7.59 (m, 1H), 7.59 – 7.51 (m, 1H), 7.48 (m, 1H), 7.46 – 7.35 (m, 1H), 7.32 (d, *J* = 8.8 Hz, 1H), 7.29 (d, *J* = 8.0 Hz, 0.5H), 4.91 (d, *J* = 5.9 Hz, 2H), 4.78 (t, *J* = 12.4 Hz, 2H), 4.60 (d, *J* = 5.4 Hz, 2H), 4.15 (ddd, *J* = 8.1, 5.6, 2.6 Hz, 2H), 3.09 (s, 3H), 2.47 (dt, *J* = 6.2, 2.5 Hz, 1H). $^{13}$C NMR (151 MHz, $CD_3CN$) δ 167.4, 167.4, 165.7, 157.3, 157.3, 141.8, 141.8, 140.8, 140.7, 138.6, 138.2, 137.4, 136.9, 136.1, 136.1, 135.8, 135.8, 131.3, 131.0, 131.0, 130.2, 130.1, 129.3, 128.4, 127.7, 127.6, 127.2, 127.2, 126.6, 124.6, 124.2, 122.7, 122.3, 114.3, 114.3, 81.4, 81.4, 71.8, 66.2, 66.2, 66.0, 66.0, 65.8, 65.8, 53.8, 53.6, 52.5, 52.3, 44.7, 29.6, 29.6.

*3-(2-(5-(methylsulfonyl)-2-(2,2,3,3,3-pentafluoropropoxy)benzoyl)isoindolin-5-yl)-N-(pent-4-yn-1-yl)benzamide* ***(m-9f)***

In a flame-dried microwave vial and under inert conditions, a solution of 3-(2-(5-(methylsulfonyl)-2-(2,2,3,3,3-pentafluoropropoxy)benzoyl)isoindolin-5-yl)benzoic acid **8** (20 mg, 0.035 mmol, 1 eq.), HBTU (15 mg, 0.039 mmol, 1.1 eq.) and *N*-ethyl-diisopropylamine (23 mg, 0.18 mmol, 5 eq.) were dissolved in anhydrous DMF (0.3

mL) and stirred at room temperature for 20 minutes. A solution of pent-4-yn-1-amine (2.9 mg, 0.035 mmol, 1 eq.) in DMF (0.3 mL) was added dropwise and the solution was stirred at room temperature for 18 hours. After completion, the reaction mixture was diluted with DCM (2 mL) and washed three times with an aqueous solution of HCl 1 M (2 mL) and then with an aqueous solution of $NaHCO_3$. The organic phase was dried over anhydrous $Na_2SO_4$, filtered, and evaporated under vacuum. The crude that was purified by RP flash column chromatography, providing the desired compound in 72% yield (16 mg, 0.025 mmol). $^1$H NMR (600 MHz, $CDCl_3$) δ 8.07 – 8.04 (m, 1H), 8.03 – 7.96 (m, 2H), 7.74 – 7.71 (m, 1H), 7.71 – 7.64 (m, 1H), 7.57 (dd, *J* = 8.1, 1.6 Hz, 1H), 7.55 – 7.47 (m, 1.5H), 7.43 (d, *J* = 8.0 Hz, 0.5H), 7.37 (s, 0.5H), 7.23 (d, *J* = 7.9 Hz, 0.5H), 7.16 (dd, *J* = 8.8, 1.6 Hz, 1H), 6.49 – 6.39 (m, 0.5H) 5.03 (d, *J* = 10.5 Hz, 2H), 4.65 – 4.54 (m, 4H), 3.62 (dq, *J* = 12.6, 6.5 Hz, 2H), 3.08 (d, *J* = 0.9 Hz, 3H), 2.35 (ddt, *J* = 10.5, 7.0, 3.4 Hz, 2H), 2.04 – 1.97 (m, 1H), 1.89 (dp, *J* = 11.4, 6.8 Hz, 2H). $^{13}$C NMR (151 MHz, $CDCl_3$) δ 167.6, 167.5, 165.1, 156.8, 141.3, 140.7, 140.4, 136.8, 136.7, 135.8, 135.5, 135.5, 135.4, 130.9, 130.9, 130.3, 130.2, 129.3, 129.0, 129.0, 128.4, 128.4, 127.4, 127.2, 126.1, 126.1, 125.8, 125.7, 123.7, 123.2, 122.0, 121.4, 113.3, 83.8, 77.4, 77.2, 76.9, 69.5, 66.0, 65.8, 65.6, 53.3, 53.2, 52.3, 52.1, 44.8, 39.6, 39.6, 28.2, 28.1, 16.5, 16.5.

*3-(2-(5-(methylsulfonyl)-2-(2,2,3,3,3-pentafluoropropoxy)benzoyl)isoindolin-5-yl)-N-(2-(2-(prop-2-yn-1-yloxy)ethoxy)ethyl)benzamide* ***(m-9g)***

In a flame-dried microwave vial and under inert conditions, a solution of 3-(2-(5-(methylsulfonyl)-2-(2,2,3,3,3-pentafluoropropoxy)benzoyl)isoindolin-5-yl)benzoic acid **8** (20 mg, 0.035 mmol, 1 eq.), HBTU (15 mg, 0.039 mmol, 1.1 eq.) and *N*-ethyl-diisopropylamine (23 mg, 0.18 mmol, 5 eq.) were dissolved in anhydrous DMF (0.3 mL) and stirred at room temperature for 20 minutes. A solution of 2-(2-(prop-2-yn-1-yloxy)ethoxy)ethan-1-amine (5 mg, 0.035 mmol, 1 eq.) in DMF (0.3 mL) was added dropwise and the solution was stirred at room temperature for 18 hours. After completion, the reaction mixture was diluted with DCM (2 mL) and washed three times with an aqueous solution of HCl 1 M (2 mL) and then with an aqueous solution of $NaHCO_3$. The organic phase was dried over anhydrous $Na_2SO_4$, filtered, and evaporated under vacuum. The crude that was purified by RP flash column chromatography, providing the desired compound in 61% yield (15 mg, 0.022 mmol). $^1$H NMR (600 MHz, $CDCl_3$) δ 8.08 – 8.03 (m, 1.5H), 8.03 – 7.97 (m, 1.5H), 7.77 – 7.70 (m, 1.5H), 7.66 (d, *J* = 7.7 Hz, 0.5H), 7.60 – 7.57 (m, 1H), 7.56 – 7.46 (m, 1.5H), 7.44 (d, *J* = 7.8 Hz, 0.5H), 7.38 (s, 0.5H), 7.23 (d, *J* = 7.9 Hz, 0.5H), 7.16 (dd, *J* = 8.8, 1.6 Hz, 1H), 5.03 (d, *J* = 11.3 Hz, 2H), 4.67 – 4.56 (m, 4H), 4.16 (dd, *J* = 11.0, 2.4 Hz, 2H), 3.75 – 3.67 (m, 8H), 3.08 (s, 3H), 2.39 (dt, *J* = 9.9, 2.4 Hz, 1H). $^{13}$C NMR (151 MHz, $CDCl_3$) δ 167.5, 167.4, 165.1, 156.8, 141.3, 140.7, 140.5, 136.8, 136.7, 135.8, 135.5, 135.3, 130.9, 130.9, 130.2, 130.2, 129.2, 129.1,

128.4, 128.4, 127.5, 127.2, 126.3, 126.3, 126.0, 125.9, 123.7, 123.1, 122.0, 121.5, 113.3, 79.5, 74.9, 74.9, 70.2, 70.2, 70.0, 69.9, 69.2, 69.2, 66.0, 65.8, 65.6, 58.6, 58.5, 53.3, 53.2, 52.3, 52.1, 44.8, 39.9, 39.9, 29.9.

*N-((1-(3-(3-(5,5-difluoro-7-(1H-pyrrol-2-yl)-5H-5λ4,6λ4-dipyrrolo[1,2-c:2',1'-f][1,3,2]diazaborinin-3-yl)propanamido)propyl)-1H-1,2,3-triazol-4-yl)methyl)-3-(2-(5-(methylsulfonyl)-2-(2,2,3,3,3-pentafluoropropoxy)benzoyl)isoindolin-5-yl)benzamide **(m-1e)***

In a microwave vial, a solution of ***m*-9e** (4.64 mg, 7.65 μmol, 1.5 eq.) and BODIPY 590-azide (2.10 mg, 5.10 μmol, 1 eq.) in DMF (0.5 mL) were degassed for 1 hour. A degassed mixture of 3,3',3''-((nitrilotris(methylene))tris(1H-1,2,3-triazole-4,1-diyl))tris(propan-1-ol) (8.86 mg, 20.4 μmol, 4 eq.) and Tetrakis(acetonitrile)copper(I) triflate (3.84 mg, 10.2 μmol, 2 eq.) in degassed milliQ water, was added dropwise to the vial under argon atmosphere. The reaction mixture was stirred at room temperature for 18 hours, diluted in acetonitrile, and purified by preparative HPLC (gradient from 20% to 100% B over 20 minutes) providing ***m*-1e** in 87% yield (4.5 mg, 4.4 μmol). HRMS (MALDI) calcd for $C_{48}H_{44}BF_7N_9O_6S^+$ (M+H$^+$) 1018.3110, found 1018.3119. $^1$H NMR (600 MHz, $CD_3CN$) δ 10.36 (s, 1H), 8.09 – 7.99 (m, 2H), 7.91 (d, *J* = 2.4 Hz, 1H), 7.83 – 7.72 (m, 3H), 7.68 – 7.49 (m, 3H), 7.45 (d, *J* = 8.3 Hz, 1H), 7.31 (dd, *J* = 8.8, 4.8 Hz, 1H), 7.25 (d, *J* = 7.9 Hz, 1H), 7.21 (s, 1H), 7.20 – 7.18 (m, 2H), 7.11 (s, 1H), 6.97 (d, *J* = 4.6 Hz, 1H), 6.93 (t, *J* = 3.6 Hz, 1H), 6.52 (d, *J* = 8.4 Hz, 1H), 6.38 – 6.33 (m, 1H), 6.31 (t, *J* = 4.4 Hz, 1H), 4.89 (d, *J* = 7.1 Hz, 2H), 4.77 (td, *J* = 12.4, 4.8 Hz, 2H), 4.59 (d, *J* = 26.7 Hz, 4H), 4.32 (q, *J* = 7.6 Hz, 2H), 3.18 (d, *J* = 7.8 Hz, 4H), 3.09 (d, *J* = 2.7 Hz, 3H), 2.62 – 2.49 (m, 2H), 2.01 (q, *J* = 7.1 Hz, 2H). $^{13}$C NMR (151 MHz, $CD_3CN$) δ 172.7, 167.7, 157.3, 156.7, 151.3, 145.8, 141.8, 141.7, 140.8, 140.7, 138.5, 138.3, 138.1, 137.3, 136.8, 136.1, 134.4, 133.2, 131.3, 130.9, 130.8, 130.1, 130.1, 129.2, 128.4, 128.4, 127.8, 127.7, 127.6, 127.3, 127.2, 126.6, 125.0, 124.5, 124.2, 124.2, 122.7, 122.3, 121.2, 118.9, 117.3, 114.3, 114.3, 112.3, 66.2, 66.1, 66.0, 66.0, 65.8, 65.8, 53.8, 53.6, 52.5, 52.4, 48.7, 44.7, 44.7, 37.0, 37.0, 35.9, 35.3, 30.8, 30.8, 25.1. Purity by analytical HPLC 97% (Rt =17.21 min).

*N-(3-(1-(3-(3-(5,5-difluoro-7-(1H-pyrrol-2-yl)-5H-5λ4,6λ4-dipyrrolo[1,2-c:2',1'-f][1,3,2]diazaborinin-3-yl)propanamido)propyl)-1H-1,2,3-triazol-4-yl)propyl)-3-(2-(5-(methylsulfonyl)-2-(2,2,3,3,3-pentafluoropropoxy)benzoyl)isoindolin-5-yl)benzamide **(m-1f** or **GlyT1-probe)***

In a microwave vial, a solution of ***m*-9f** (5.79 mg, 9.12 μmol,1.5 eq.) and BODIPY 590-azide (2.50 mg, 6.08 μmol, 1 eq.) in DMF (0.5 mL) were degassed for 1 hour. A degassed mixture of 3,3',3''-((nitrilotris(methylene))tris(1H-1,2,3-triazole-4,1-diyl))tris(propan-1-ol) (10.6 mg, 24.3μmol, 4 eq.) and Tetrakis(acetonitrile)copper(I) triflate (4.58 mg, 12.2 μmol, 2 eq.) in degassed milliQ water, was added dropwise to the vial under argon atmosphere. The reaction mixture was stirred at room temperature for 18 hours, diluted in acetonitrile, and purified by preparative HPLC (gradient from 0% to 100% B over 30 minutes) providing ***m*-1f** in 91% yield (5.8 mg, 5.5 µmol). HRMS (MALDI) calcd for $C_{50}H_{47}BF_7N_9O_6S$ (M+H$^+$) 1046.3423, found 1046.3442. $^1$H NMR (600 MHz, $CD_3CN$) δ 10.36 (s, 1H), 8.06 – 8.01 (m, 1.5H), 7.99 (t, *J* = 1.8 Hz, 0.5H), 7.91 (dd, *J* = 2.4, 1.0 Hz, 1H), 7.80 – 7.74 (m, 1.5H), 7.72 (ddd, *J* = 7.8, 1.9, 1.1 Hz, 0.5H), 7.70 (d, *J* = 1.7 Hz, 0.5H), 7.63 (dd, *J* = 7.9, 1.7 Hz, 0.5H), 7.62 – 7.58 (m, 1.5H), 7.52 (dt, *J* = 18.0, 7.8 Hz, 1H), 7.48 – 7.45 (m, 1H),f 7.31 (dd, *J* = 8.9, 2.1 Hz, 1H), 7.26 (d, *J* = 7.9 Hz, 0.5H), 7.22 – 7.17 (m, 3H), 7.13 – 7.10 (m, 1H), 6.97 (dd, *J* = 4.6, 2.3 Hz, 1H), 6.93 (t, *J* = 3.9 Hz, 1H), 6.60 – 6.55 (m, 0.5H), 6.54 – 6.50 (m, 0.5H), 6.35 (dt, *J* = 4.1, 2.4 Hz, 1H), 6.32 (d, *J* = 3.9 Hz, 1H), 4.90 (d, *J* = 7.9 Hz, 2H), 4.77 (t, *J* = 12.4 Hz, 2H), 4.58 (d, *J* = 11.5 Hz, 2H), 4.28 (dt, *J* = 12.5, 6.9 Hz, 2H), 3.42 (dq, *J* = 14.6, 6.6 Hz, 2H), 3.21 (td, *J* = 7.7, 5.6 Hz, 2H), 3.14 (dq, *J* = 13.0, 6.4 Hz, 2H), 3.09 (d, *J* = 1.5 Hz, 3H), 2.76 (dt, *J* = 14.0, 7.3 Hz, 2H), 2.56 (td, *J* = 7.7, 6.6 Hz, 4H), 1.99 (dt, *J* = 12.7, 6.6 Hz, 2H). $^{13}$C NMR (151 MHz, $CD_3CN$) δ 172.6, 172.6, 167.9, 167.8, 165.8, 165.7, 157.3, 157.2, 156.8, 151.3, 147.7, 141.7, 141.7, 140.9, 140.8, 138.6, 138.3, 138.1, 137.2, 136.8, 136.6, 136.6, 136.1, 136.1, 134.4, 133.2, 131.3, 130.6, 130.6, 130.1, 130.0, 129.3, 129.3, 128.4, 128.4, 127.8, 127.7, 127.6, 127.3, 127.2, 127.1, 126.5, 125.8, 125.0, 124.5, 124.2, 124.1, 123.2, 123.2, 122.7, 122.3, 121.2, 118.9, 117.3, 114.3, 114.3, 112.3, 66.1, 66.0, 66.0, 65.8, 53.8, 53.6, 52.5, 52.3, 48.5, 48.5, 44.7, 44.7, 40.0, 39.9, 36.9, 36.9, 35.3, 35.3, 30.8, 30.8, 29.7, 29.6, 25.1, 23.5, 23.5. Purity by analytical HPLC 100% (Rt =17.25 min).

*N-(2-(2-((1-(3-(3-(5,5-difluoro-7-(1H-pyrrol-2-yl)-5H-5λ4,6λ4-dipyrrolo[1,2-c:2',1'-f][1,3,2]diazaborinin-3-yl)propanamido)propyl)-1H-1,2,3-triazol-4-yl)methoxy)ethoxy)ethyl)-3-(2-(5-(methylsulfonyl)-2-(2,2,3,3,3-pentafluoropropoxy)benzoyl)isoindolin-5-yl)benzamide* ***(m-1g)***

In a microwave vial, a solution of ***m*-9g** (4.00 mg, 5.76 µmol,1.5 eq.) and BODIPY 590-azide (2.37 mg, 5.76 µmol, 1 eq.) in DMF (0.5 mL) were degassed for 1 hour. A degassed mixture of 3,3',3''-((nitrilotris(methylene))tris(1H-1,2,3-triazole-4,1-diyl))tris(propan-1-ol) (10.0 mg, 23.0 µmol, 4 eq.) and Tetrakis(acetonitrile)copper(I) triflate (4.34 mg, 11.5 µmol, 2 eq.) in degassed milliQ water, was added dropwise to the vial under argon atmosphere. The reaction mixture was stirred at room temperature for 18 hours, diluted in acetonitrile, and purified by preparative HPLC (gradient from 20% to 100% B over 30 minutes) providing ***m*-1f** in 82% yield (5.2 mg, 4.7 µmol). HRMS (MALDI) calcd for $C_{52}H_{51}BF_7N_9O_8S$ (M+H$^+$) 1106.3634, found 1106.3652. $^1$H NMR (600 MHz, $CD_3CN$) δ 10.37 (s, 1H), 8.05 – 8.01 (m, 1.5H), 7.99 (m, 0.5H), 7.91 (d, *J* = 2.4 Hz, 1H), 7.79 – 7.72 (m, 1.5H), 7.72 – 7.66 (m, 1.5H), 7.63 (s, 0.5H), 7.59 (d, *J* = 8.0 Hz, 0.5H), 7.55 (d, *J* = 7.9 Hz, 0.5H), 7.53 – 7.39 (m, 3H), 7.31 (d, *J* = 8.8 Hz, 1H), 7.24 (d, *J* = 7.9 Hz, 0.5H), 7.23 – 7.17 (m, 3H), 7.14 –

7.09 (m, 1H), 6.97 (d, *J* = 4.5 Hz, 1H), 6.95 – 6.91 (m, 1H), 6.60 (m, 1H), 6.35 (m, 1H), 6.31 (d, *J* = 4.0 Hz, 1H), 4.88 (s, 2H), 4.77 (t, *J* = 12.4 Hz, 2H), 4.56 (s, 3H), 4.54 (s, 1H), 4.27 (dt, *J* = 10.9, 6.9 Hz, 2H), 3.67 – 3.57 (m, 6H), 3.53 (dq, *J* = 11.3, 5.5 Hz, 2H), 3.20 (td, *J* = 7.8, 4.2 Hz, 2H), 3.14 – 3.09 (m, 2H), 3.09 (s, 3H), 2.55 (td, *J* = 7.7, 2.8 Hz, 2H), 2.04 – 1.96 (m, 2H). $^{13}C$ NMR (151 MHz, $CD_3CN$) δ 165.7, 157.3, 157.2, 156.8, 151.3, 145.4, 141.7, 141.6, 140.8, 140.7, 138.5, 138.3, 138.1, 137.2, 136.8, 136.1, 134.3, 133.2, 131.3, 130.7, 130.6, 130.1, 130.0, 129.3, 129.3, 128.4, 128.4, 127.8, 127.7, 127.5, 127.3, 127.2, 127.1, 126.6, 125.0, 124.5, 124.4, 124.2, 124.1, 122.6, 122.2, 121.2, 118.9, 118.3, 117.3, 114.3, 114.3, 112.3, 71.0, 71.0, 70.4, 70.2, 70.2, 66.2, 66.1, 66.0, 65.9, 65.8, 65.8, 64.8, 64.8, 53.7, 53.6, 52.5, 52.3, 48.4, 48.3, 44.7, 40.5, 40.5, 36.9, 36.9, 35.3, 32.6, 30.8, 29.6, 25.1, 23.3, 14.3, 1.7, 1.5, 1.4, 1.3, 1.1, 1.0, 0.8. Purity by analytical HPLC 99% (Rt =24.06 min, gradient over 30 min).
